# Identification, structure and function of the methyltransferase involved in the biosynthesis of the dithiolopyrrolone antibiotic xenorhabdin

**DOI:** 10.1101/2024.01.12.575338

**Authors:** Li Su, Eva M. Huber, Margaretha Westphalen, Jonas Gellner, Edna Bode, Tania Köbel, Peter Grün, Mohammad M. Alanjary, Timo Glatter, Daniel Schindler, Michael Groll, Helge B. Bode

**Author notes:** These authors contributed equally (Li Su, Eva M. Huber).

## Abstract

Xenorhabdins (XRDs) are produced by *Xenorhabdus* species and are members of the dithiopyrrolone (DTP) class of natural products that have potent antibacterial, antifungal and anticancer activity. The amide moiety of their DTP core can be methylated or not to fine-tune the bioactivity properties. However, the enzyme responsible for the amide *N*-methylation remained elusive. Here, we identified and characterized the amide methyltransferase XrdM that is encoded nearly 600 kb away from the XRD gene cluster using proteomic analysis, methyltransferase candidate screening, gene deletion, and allied approaches. In addition, crystallographic analysis and site-directed mutagenesis proved that XrdM is completely distinct from the recently reported DTP methyltransferase DtpM, and that both have been tailored in a species-specific manner for DTP biosynthesis in Gram-negative/positive organisms. Our study expands the limited knowledge of post-NRPS amide methylation in DTP biosynthesis and reveals the evolution of two structurally completely different enzymes for the same reaction in different organisms.

## Introduction

Xenorhabdins (XRDs) are members of the dithiopyrrolone (DTP) class of natural products. DTPs are structurally unique and exhibit broad-spectrum bioactivity against a variety of organisms, including Gram-positive and Gram-negative bacteria, fungi, and even eukaryotic parasites.^1^ DTPs share the bicyclic dithiopyrrolone scaffold, which is biosynthesized by a non-ribosomal peptide synthetase (NRPS) from two cysteine building blocks followed by cyclization, decarboxylation and oxidation.^2^ Tailoring enzymes, such as amino acyltransferase and amide *N*-methyltransferase (MT), further expand the structural diversity of DTPs, which according to their modifications can be grouped into three subfamilies: *N*-acyl-DTPs (*e.g.* holomycin), *N*-acyl, *N*-methyl-DTPs (*e.g.* thiolutin), and thiomarinol, a distinct type carrying the polyketide antibiotic marinolic acid as the *N*-acyl group linked to DTP (**Figure S1**).^3^

To date, only thiolutin and derivatives thereof (*e.g.* aureothricin), as well as a few members of XRDs (referring to as *N*-methylated XRDs in the following), such as XRD-243Me, XRD-271Me, XRD-285Me, and XRD-299Me were identified as naturally occurring *N*-acyl, *N*-methyl-DTPs (**Figure 1a, b and Figure S1**). In analogy to holomycin and thiolutin, which differ only in the *N*-methyl group at the endocyclic amide nitrogen position, there are non-methylated and *N*-methylated XRDs. Interestingly, while the methylated derivatives show a better antifungal activity, the non-methylated derivatives are better antibiotics against Gram-positive and Gram-negative bacteria (**Table S1**), implying that the bioactivity of DTPs is modulated by the amide methyl group.^4,5^

**Figure 1.**
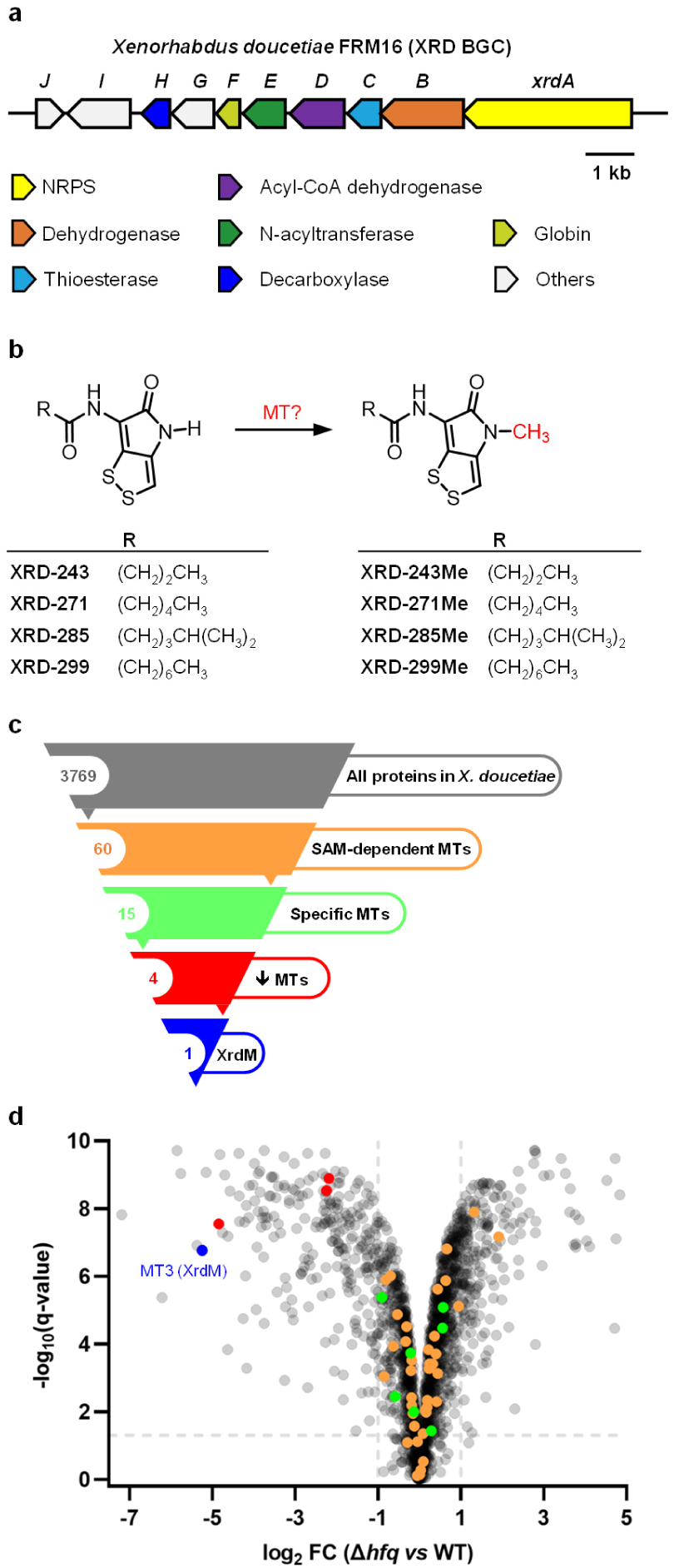
Identification of XRD *N*-methyltransferase. **a**) Biosynthetic gene cluster of XRD in *X. doucetiae* FRM16 not encoding an *N*-methyltransferase. **b**) Structures of the naturally produced non-methylated and *N*-methylated XRDs with the *N*-methyl moiety highlighted. **c**) Screening funnel for XRD-specific *N*-methyltransferase leading to 15 MT candidates for testing by protein expression, of which 4 were further checked by gene deletion. XrdM alone was found to be responsible for the *N*-methylation of XRDs at the end. **d**) Volcano plot of the comparative proteomics analysis between *X. doucetiae* Δ*hfq* mutant and its wild type (WT). Dashed lines show cutoff values q = 0.05, and FC (fold change) = 2. Dots in blue, red, green, and orange refer to MTs according to the screening funnel shown in **c**. Among them, the green and red dots were MTs specific (**Figure S4**) for *X. doucetiae* and thus were chosen for a test by protein expression in Δ*hfq* P*_BAD_*_xrdA. In addition, the blue (MT3 or XrdM) and red dots were down-regulated MTs and were tested by gene deletion in WT, respectively. Data were calculated from four biological replicates. Detailed information is given in **Figure S4**. and **Extended Excel sheet_2**.

Usually, methylation of NRPS products is accomplished by a MT domain within the NRPS or by a separate MT within the corresponding biosynthetic gene cluster (BGC). However, also MTs encoded outside the corresponding BGC can be hijacked for tailoring reactions. Prominent examples are the *S*- MT from gliotoxin biosynthesis^6,7^ and the *N*-MT converting holomycin to thiolutin^8^. Similarly, XRD MT is neither encoded within the NRPS nor as a separate enzyme within the XRD BGC^9^. To selectively generate XRD and other DTP derivatives with or without *N*-methyl groups and close the knowledge gap in DTP biosynthesis, we aimed to identify the XRD MT. It’s worth mentioning that the *N*-MT of holomycin – DtpM was indentified recently from *Sacchrothrix algeriensis* NRRL B-24137 via BLASTP similarity search querying the terpenoid *N*-MT WelM^8^. However, a similar strategy for the identification of the XRD MT in *Xenorhabdus doucetiae* FRM16 failed even when DtpM was used as the bait.

Here, we identified and characterized the amide *N*-MT XrdM that is encoded nearly 600 kb away from the XRD gene cluster using proteomic analysis, MT-candidate screening, gene deletion, and allied approaches. We further showed that XrdM and DtpM are isofunctional enzymes toward XRDs and holomycin. Additionally, X-ray crystal structures, molecular modeling and site-directed mutagenesis revealed differences between XrdM and DtpM and deliver insights into their catalytic mechanisms on DTP methylation.

## Results

### Identification of the XRD MT XrdM

Since our previous study revealed the MT involved in XRD methylation to be SAM-dependent^9^, we screened the *X. doucetiae* genome for all the SAM-dependent MTs and found 60 candidates (**Figure 1c, Table S2**). 45 of them were excluded based on their presence and high sequence identity (cutoff > 60%) to proteins encoded in *X. nematophila* ATCC 19061 and *X. szentirmaii* DSM 16338, two *Xenorhabdus* species that are evolutionarily similar to *X. doucetiae* but do not carry the XRD BGC^10^ (**Table S2**). Indeed, we could show that, in contrast to *X. doucetiae*, neither *X. nematophila* ATCC 19061 nor *X. szentirmaii* DSM 16338 were able to methylate any of the XRD derivatives (**Figure S2**). The remaining 15 MT candidates were considered to be specific in *X. doucetiae* and were therefore selected for further experiments (**Figure 1c, Table S2**).

In addition, a previously generated Δ*hfq* mutant, in which the gene encoding Hfq (an RNA chaperone mediating mRNA/sRNA interaction) was deleted, revealed a deprived ability to produce XRDs (**Figure S3**).^11^ In the easyPACiD (easy Promoter Activated Compound Identification) approach^11,12^, the Δ*hfq* mutant can be used for the selective production of specialized metabolites using promoter-exchange. Therefore, we exchanged the natural promoter of NRPS *xrdA* with an arabinose inducible promoter, resulting in strain Δ*hfq* P*_BAD_*_*xrdA* (**Table S3**). Upon induction, this strain produces mostly non-methylated XRDs (XRD-243, -271, and -299) with only trace amount of methylated XRD- 271Me, whereas in WT, the *N*-methylated XRDs (XRD-243Me, -271Me, and -299Me) are the main derivatives (**Figure S3**). These findings imply that the transcription or translation of both XRD BGC and the corresponding XRD MT were down-regulated in the Δ*hfq* mutant in comparison to the WT. Promoter exchange of *xrdA* restored the transcription or translation of XRD BGC but not of XRD MT, which accordingly makes Δ*hfq* P*_BAD_*_*xrdA* an ideal host for the expression of selected MT candidates. A comparative proteomic analysis between Δ*hfq* and WT verified the down-regulation of proteins encoded by XRD BGC (**Figure S4**) and revealed that 4 of the 15 MT candidates are also down-regulated, referred to as MT3, MT13, MT14, and MT15. For a second line of verification, these four MT candidates were individually selected for gene knockouts in WT to investigate their contribution to XRD methylation *in vivo* (**Figure 1d and Figure S4**).

MT candidates (MT1 – MT15) were separately cloned into vector pACYC and overexpressed in *X. doucetiae* Δ*hfq* P*_BAD_*_*xrdA*. Extracted ion chromatograms (EIC) for signals of XRDs from these strains indicated that only the strain Δ*hfq* P*_BAD_*_*xrdA* pMT3 (**Table S3**) yields *N*-methylated XRDs, as opposed to the non-methylated XRDs that were produced by the remaining 14 strains and the empty plasmid pACYC-containing control strain (**Figure 2a and Figure S5**). Accordingly, deletion of the *MT3* gene in WT changed the XRD production profile from methylated to non-methylated XRDs, while mutants with the deletion of *MT13, 14,* or *15* did not (**Figure 2b**). Together, these results confirmed that MT3, located 592 kb away from the XRD BGC, is the only MT in the producing strain that catalyzes the conversion of non-methylated XRDs to *N*-methylated XRDs, and thus it was named XrdM (<u>X</u>enorhab<u>d</u>in <u>m</u>ethyltransferase) (**Figure S1**).

**Figure 2.**
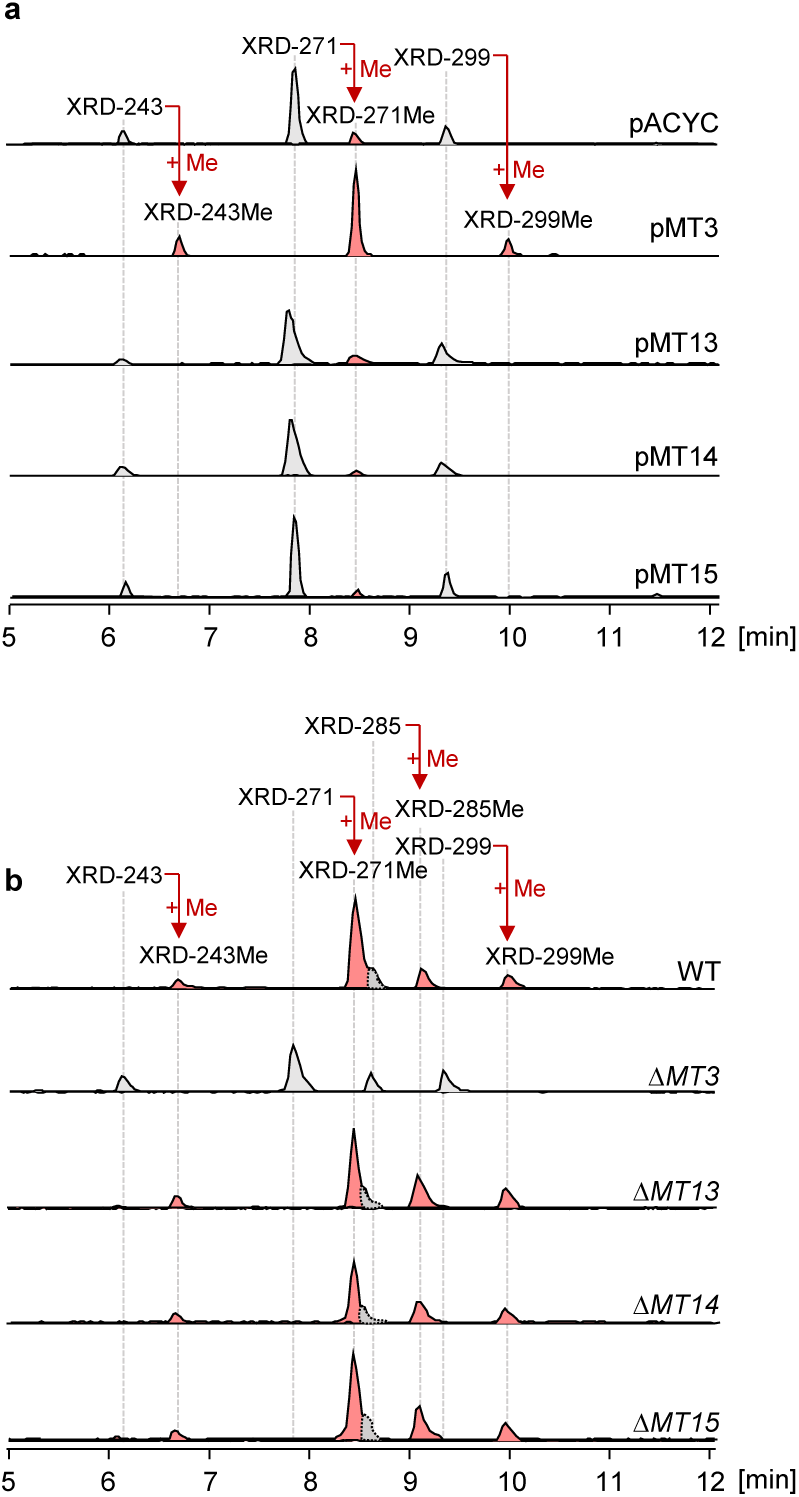
Extracted ion chromatograms (EIC) of XRDs from plasmid expression strains (**a**) and gene-deletion strains (**b**). Peaks referring to non-methylated XRDs and *N*-methylated XRDs were colored in grey and red, respectively. pACYC represents the mutant Δ*hfq* P*_BAD_*_xrdA containing an empty plasmid pACYC (see details in **Table S3**), serving as control for protein expression strains having pMT3, 13, 14, and 15, respectively. Results of all MT expression strains are shown in **Figure S5**.

### XrdM and DtpM are isofunctional enzymes

A BLASTP search of XrdM against the genome of *S. algeriensis* NRRL B-24137 led to several hits, in which the top hit CTG1_5739 (22.7% sequence identity) was the closest XrdM homolog (**Figure S6**). However, the expression of CTG1_5739 in Δ*hfq* P*_BAD_*_*xrdA* did not show any methylation towards non- methylated XRDs (**Figure S7**). We initially thought CTG1_5739, the putative MT of holomycin in *S. algeriensis* NRRL B-24137, might have substrate specificity to holomycin and thus fails to methylate the long acyl-chain decorated XRDs. To test this hypothesis, we produced holomycin in *X. doucetiae* FRM16. Since holomycin and XRDs only differ in the length of their acyl side chain, we replaced the XRD specific acyltransferase gene *xrdE* in strain Δ*hfq* P*_BAD_*_*xrdA* by either *hlmA*, the gene encoding the acetyltransferase in holomycin BGC of *S. clavuligerus* ATCC 27064^2^, or *actA*, the coding sequence of the putative acetyltransferase involved in thiolutin biosynthesis from *S. algeriensis* NRRL B-24137^13,14^. Holomycin was produced by both resulting strains, Δ*hfq* Δ*xrdE::hlmA* P*_BAD_*_*xrdA* and Δ*hfq* Δ*xrdE::actA* P*_BAD_*_*xrdA* (**Table S3**), as confirmed by high-resolution mass spectrometry (HRMS) and MS/MS fragmentation pattern analysis (**Figures S8 and S9**). The production of holomycin in the *hlmA* strain was much better than in the *actA* strain, which additionally yielded XRD-243 and XRD-271 as byproducts. These observations are in agreement with the catalytic profiles of HlmA and ActA in their native producer strains: while holomycin is the only DTP detected in *S. clavuligerus* ATCC 27064^2^, at least six thiolutin derivatives with varying acyl chain lengths could be produced by *S. algeriensis* NRRL B-24137^15^. To test whether CTG1_5739 is the genuine MT for holomycin, we further individually introduced the plasmids pCTG1_5739 and pXrdM (same as pMT3) into the *hlmA* carrying mutant (**Table S3**). Unfortunately, CTG1_5739 could not catalyze the conversion of holomycin to thiolutin (**Figure S8**), for which task the MT DtpM was identified when we prepared this manuscript^8^.

Given the above, we intended to test if XrdM and DtpM are both active to each other’s substrates. Analysis of resulting strains Δ*hfq* Δ*xrdE::hlmA* P*_BAD_*_*xrdA* pXrdM and Δ*hfq* Δ*xrdE::hlmA* P*_BAD_*_*xrdA* pDtpM revealed that XrdM is surprisingly able to convert holomycin to thiolutin, albeit with a relatively lower activity than DtpM (**Figure S8**), showcasing the substrate promiscuity of XrdM towards *N*-acyl- DTPs that have *N*-acyl side chains in different lengths and even in branched type (*e.g.* XRD-285 and XRD-285Me, **Figure 1b**). Likewise, DtpM was also able to methylate all of the XRDs as seen from the analysis of strain Δ*hfq* P*_BAD_*_*xrdA* pDtpM (**Figure S8**). Notably, these data confirm XrdM and DtpM are isofunctional enzymes for XRDs and holomycin though they both share the amino acid sequence identity of only 14.6% (**Figure S10**).

To get an overview of the similarities between XrdM and DtpM, we performed a sequence similarity network (SSN) (**Figure 3**). This analysis revealed the close association of DtpM with a number of phenolic-*O*-MTs (e.g., LaPhzM^16^, CrmM^17^, AzicL^18^, and CalO6^19^) and several *N*-MTs (e.g., WelM^20^) belonging to the Pfam00891 family. In contrast, XrdM did not cluster with those enzymes but with other MTs of the Pfam13649 family and some Pfam13847 MTs. It must be noted that MT15, which clustered with DtpM (**Figure 3**), was the sequence homolog of DtpM we found in *X. doucetiae* FRM16 (identity of 31.8% to DtpM and 34.9% to WelM, **Figure S11**) but was inactive upon XRDs as we have demonstrated (**Figure 2**), emphasizing that identification of XRD MT based on sequence BLAST/protein similarity only was not feasible.

**Figure 3.**
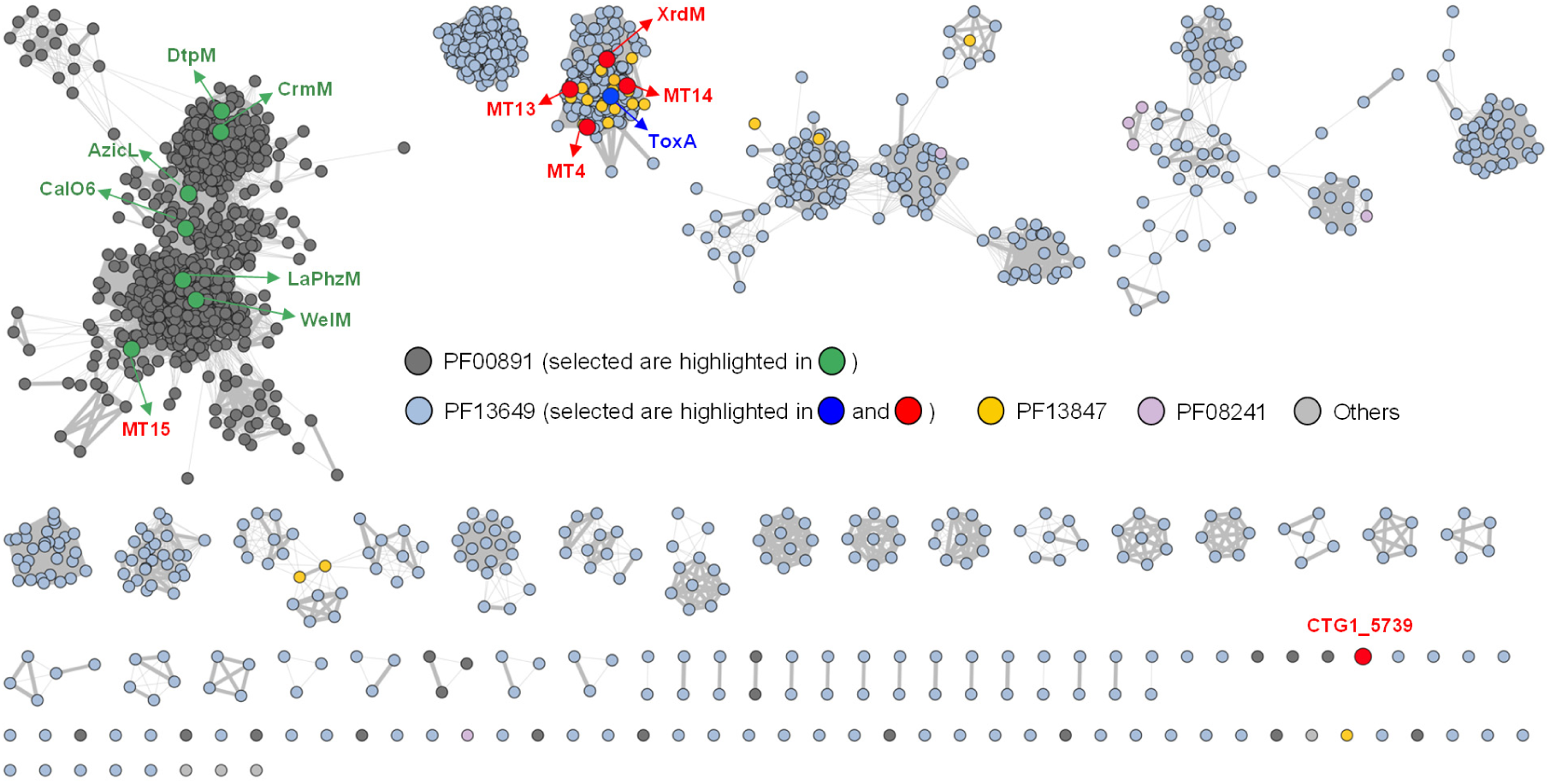
Sequence similarity network (SSN) of XrdM and DtpM. This SSN was visualized using an alignment score cutoff ≥70 (up to 331) and a sequence identity ≥30%. Nodes are colored according to which PFAM family the protein belongs to; the edge thickness is proportional to the identity (from 30% to 100%). Nodes shown in green, red, and blue were highlighted methyltransferases.

### Crystal structure of XrdM

To understand the mechanism of XRD methylation by XrdM, we determined its 3D structure using X-ray cystallography. Crystallization trials with the pure protein (**Figure S13a, b**) yielded diffraction-quality crystals that allowed us to solve the structure of XrdM (residues 4-246) in complex with the cofactor S-adenosylhomocysteine (SAH) to 2.1 Å resolution (**Table S9**) using the AlphaFold2 prediction^21^ as a molecular replacement model (**Figure S14a**). The structure of XrdM is similar to many other MTs (**Table S10**), but most related to ToxA, a dual-specificity *N*-MT that catalyzes the last two steps of toxoflavin biosynthesis^22^, as well as to glycine sarcosine *N*-MT^23^. XrdM folds into two subdomains at the interface of which a pronounced cofactor- and substrate-binding cleft is located (**Figure 4a**). The larger N-terminal domain features the canonical Rossmann motif GxGxxG which is highly conserved across class I SAM-dependent MTs^22,24,25^. The smaller domain involves residues located more C-terminally and is known as flap domain. Similar to the ToxA apo structure (PDB 5JE6^22^), the N-terminus of XrdM (residues 4-23) is disordered^22^ and parts close to the C-terminus (residues 219- 235) are not resolved in the 2F_O_-F_C_ electron density map. Due to the disorder of the N-terminal segment, the substrate binding cleft is solvent-exposed and accessible to interactions with a crystal symmetry mate. Specifically, the loop connecting residues 179-185 inserts into the substrate binding pocket of a neighboring XrdM molecule (**Figure S15**). Although the resulting interface area of 1148 Å^2^ between two XrdM monomers is classified as stable by PISA analysis, the contacts are probably non-physiological for two reasons. First, XrdM likely is a monomer in solution as determined by size exclusion chromatography (**Figure S13a**); second, the dimeric assembly in the crystal blocks the active site and probably interferes with substrate binding and conversion. Notably, such a dimerization via the flap domain has also been observed for ToxA.^22^ While in our XrdM structure SAH is bound at the active site, dimerization of ToxA even displaces the cofactor, resulting in active sites that are completely apo. In the crystal lattice two ToxA dimers further assemble into a tetramer that is supposed to be stable also in solution according to PISA analysis.^22^ By contrast, XrdM does not form homotetramers *in crystallo*. From these results, we conclude that the oligomerization of ToxA and XrdM is likely a crystallization artefact.In order to understand how XrdM methylates XRDs, we performed co-crystallization trials with XRD-271 and XRD-271Me. Unfortunately, however, all crystals obtained belonged to the same space group as the apo crystal structures and showed the same packing of molecules in the crystal lattice, suggesting that the ligands are displaced from the active site in favor of intermolecular protein contacts that drive crystal growth. Based on these results, we shortened the loop (aa 179-185) to allow ligands to bind even in the crystal lattice, but the mutant XrdM protein was not accessible to purification.

**Figure 4.**
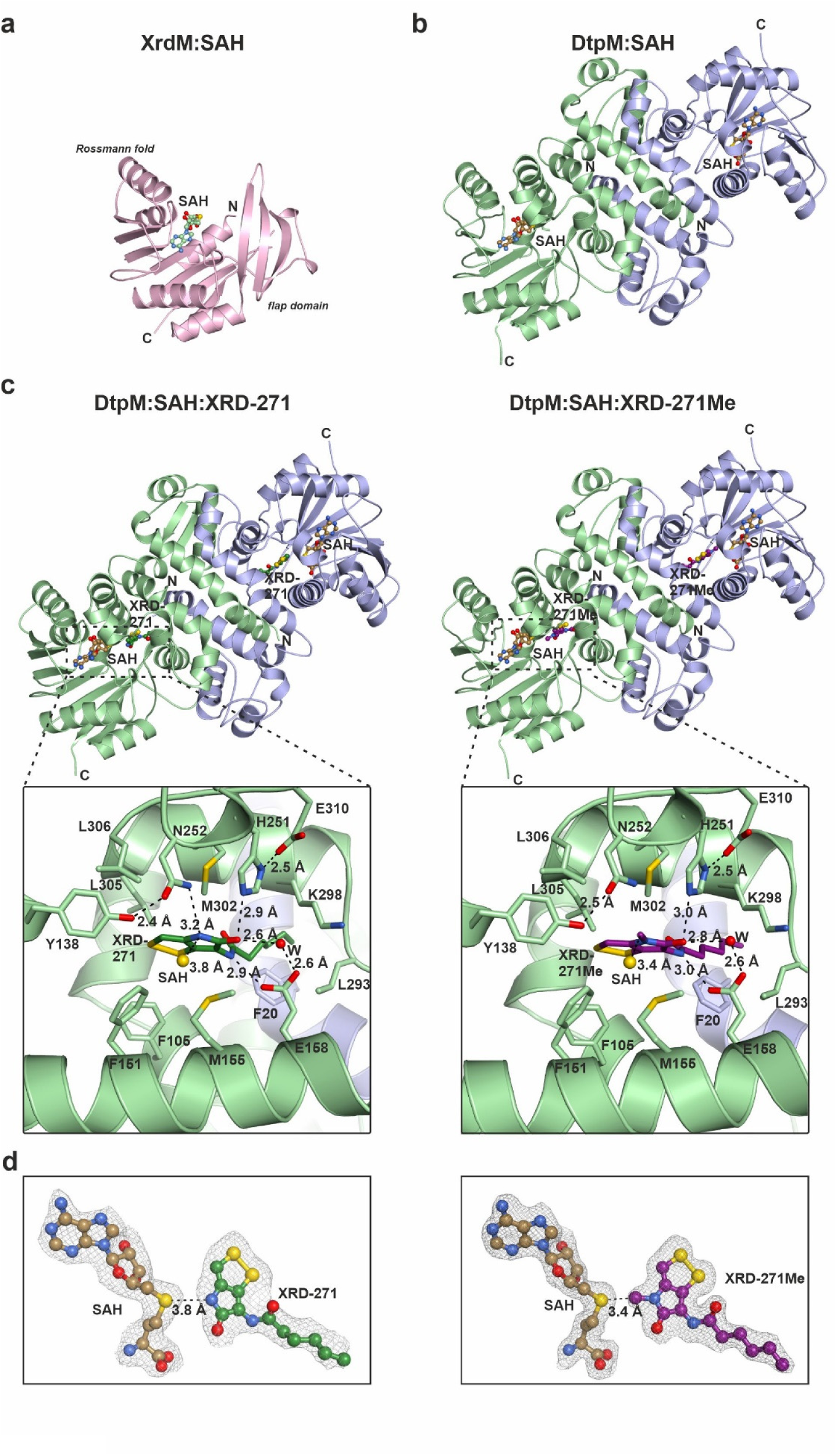
X-ray structures of XrdM and DtpM. (**a**) Ribbon illustration of monomeric XrdM with bound SAH (multicolored ball-and-stick model). N- and C-termini are labeled. (**b**) Ribbon structure of dimeric DtpM with bound SAH according to panel a. (**c**) DtpM structures with bound SAH and XRD-271 (substrate) and XRD-271Me (product), respectively. Zoom-in pictures of the active site are shown below. Amino acid residues required for ligand binding or catalysis are labeled by the one-letter code and numbered. Black dotted lines indicate important hydrogen bond interactions. Distances are given. W indicates a defined water molecule that contributes to ligand coordination. (**d**) 2F_O_-F_C_ omit electron density maps are shown for DtpM bound SAH and ligands contoured to 1σ.

### Ligand complex structures of DtpM reveal key active site residues

Considering that XRD-271 is also converted to XRD-271Me by DtpM (**Figure S12**), we crystallized DtpM and solved its structure to a resolution of 1.75 Å (**Table S9**) using the Alphafold2 prediction as a search model (**Figure S14b**). The asymmetric unit contains one DtpM molecule that forms a homodimer in the crystal lattice (**Figure 4b**). The interface area of the homodimer is 3670.8 Å^2^ large and is considered to be stable in solution by PISA analysis.^26^ Moreover, the homodimer represents a functional unit, as the N-terminus of each subunit contributes to the active site pocket of the other (**Figure 4c**). Surprisingly, DtpM is structurally related to various class I *O*-MTs.^27^ Among the ten closest structural relatives of DtpM, the Dali server^28^ highlights nine *O*-MTs and only one *N*-MT, namely the human *N*- acetlyserotonin MT (**Table S11**). The reason for this bias might be that many more *O*- than *N*-MTs are known.^29^ Each DtpM monomer adopts a two-domain fold. The N-terminal domain is required for homodimerization, the C-terminal Rossmann fold mediates cofactor binding and the pocket in between binds the substrate. Overall, DtpM shows a highly negatively charged surface (**Figure S17b**) and is most similar to the phenazine *O*-MT LaPhzM from *Lysobacter antibioticus* (PDB 6C5B, **Table S11, Figure S18**).^16^

To obtain ligand-bound structures, we performed co-crystallization trials of DtpM with SAH and either XRD-271 or XRD-271Me. This setup led to numerous yellowish crystals and diffraction data sets for both XRD-271 (2.2 Å resolution, **Table S9**) and XRD-271Me (1.75 Å resolution, **Table S9**). The structures were solved by molecular replacement with the apo DtpM coordinates and showed no conformational changes compared to the apo structure (**Figure S14d**). The experimental electron density maps however clearly visualized SAH and the respective ligand at the active site (**Figure 4d**).

The core ring stuctures of XRD-271 and XRD-271Me are stabilized in the substrate binding pocket by numerous hydrophobic interactions with surrounding amino acid residues (e.g. Phe105, Phe151, Met155, Leu293, Met 302, Leu305 and Leu306) (**Figure 4c**). The aliphatic tail of XRD-271/XRD-271Me is outstretched and points into a deep hydrophobic channel that extends to the protein surface. The amino acid side chains of Lys298, Val297, Leu293 as well as Gln293 mediate Van-der-Waals contacts with the tail of XRD-271/XRD-271Me. In addition, Phe21 and Arg23 from the neiboring DtpM monomer contribute non-polar contacts to this binding site (**Figure 4c**). Based on the pronounced depth of the channel, substrate derivatives with longer or shorter aliphatic tails (e.g. holomycin with its shorter acetyl side chain) than that of XRD-271 are likely to fit as well. Besides the numerous hydrophobic interactions, several hydrogen bonds help recognizing XRD-271 and presumably also holomycin. Glu158 for example hydrogen bonds directly to the amide nitrogen atom of the side chain of XRD-271 and indirectly via a water molecule to the amide oxygen atom of the dithiopyrrolone moiety. The latter is also coordinated by His251, which in turn is hydrogen bridged to Glu310 (**Figure 4c**). This hydrogen bond network is supposed to drain off electrons and to faciliate deprotonation of the neigboring nitrogen atom, the target site of methyl transfer. The nucleophilicity of the target nitrogen atom was speculated be further enhanced by a hydrogen bond to Tyr138 via Asn252. The distance of the sulphur atom of SAH to the acceptor nitrogen atom (3.8 Å) is ideally suited for the alkylation reaction. By modelling SAM instead of SAH into the DtpM:XRD-271 active site, the angle connecting the sulfur atom of SAH, the methyl group to be transferred and the acceptor nitrogen atom of XRD-271 was found to be 166° and thus close to the linear arrangement supposed for S_N_2 reactions.

To validate the structural data, a series of mutants (Y138F, E158A, H251A, N252A, Q301A and E310Q) were generated for DtpM and tested for their *in vitro* activity towards XRD-271 (**Figure 5 and Figure S19**). Despite mediating two hydrogen bridges to XRD-271 – one direct and one indirect – mutation of Glu158 to alanine did not affect substrate turnover, suggesting that the hydrophobic contacts are sufficient for ligand binding and catalysis. Further support for this notion is provided by the mutant Q301A. Based on the structural data, Gln301 was supposed to weakly hydrogen bridge to the amide oxygen atom of the side chain of XRD-271, but activity assays with the mutant protein showed full activity. While the mutation Y138F did not compromise substrate conversion as well, Asn252 turned out to be essential for catalytic activity. From this result we infer that Asn252 is sufficient to activate the acceptor nitrogen atom in XRD-271 for methyl transfer and no further support by Tyr138 is required.

**Figure 5.**
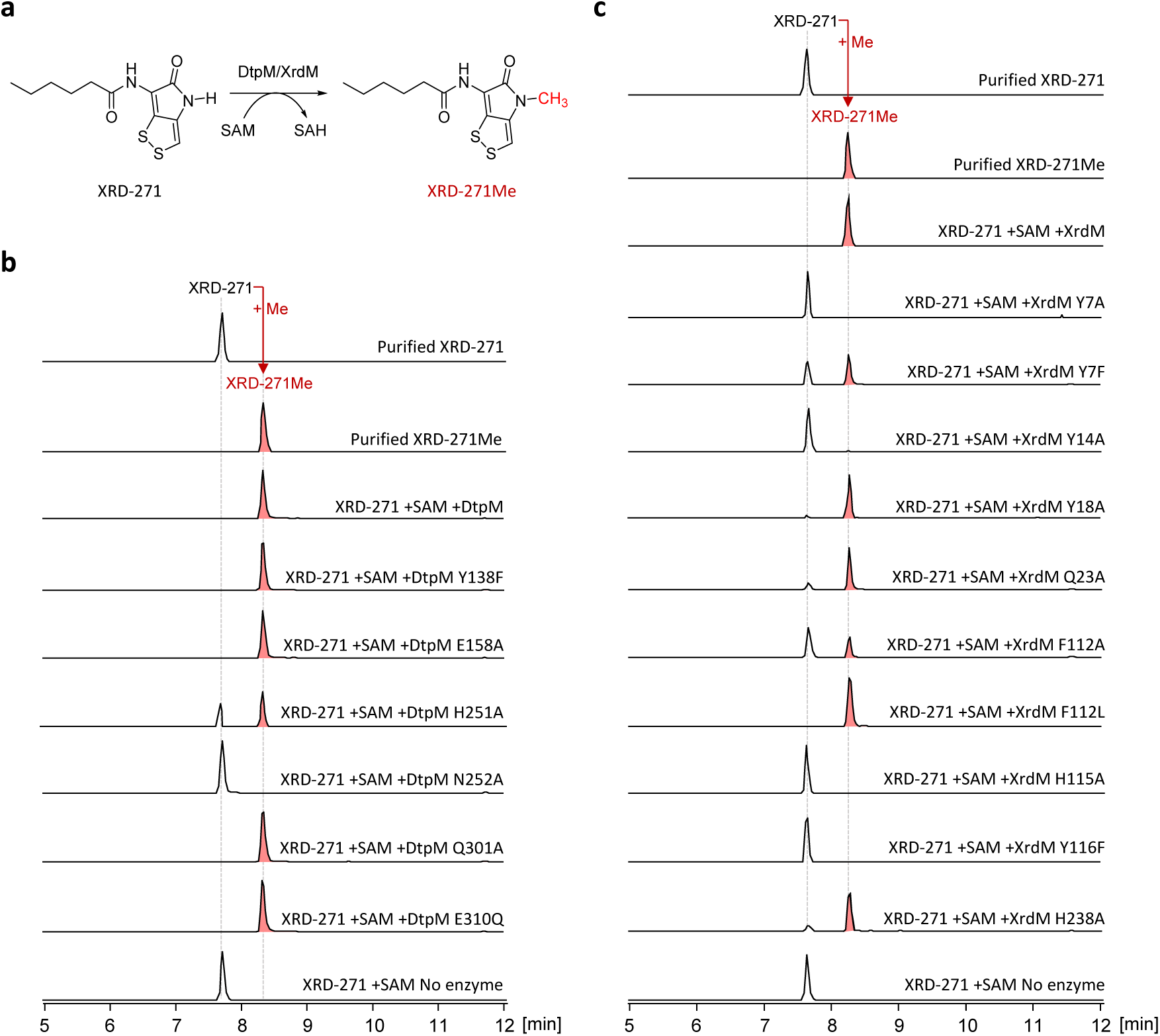
Activity assays of DtpM, XrdM and their mutants. (**a**) Methylation reaction of XRD-271 to XRD-271Me by wild type DtpM or XrdM. (**b**) Activity assay of wild type or mutant DtpM after incubation of 30 min. (**c**) Activity assay of wild type or mutant XrdM after incubation of 30min. No enzyme-added reaction was used as control and shown at the bottom, purified XRD-271 and XRD-271Me were used as standard control shown on the top. Results after incubation time of 60 min were shown in **Figure S19 and 21**.

Finally, we also analysed the relevance of the residues His251 and Glu310, which in DptM and other MTs often form a kind of dyad. While glutamine can functionally replace Glu310, probably because substitution of the side-chain carboxylate for an amide retains the hydrogen bonding pattern with His251, the H251A mutant was slowed down in substrate turnover (**Figure 5 and Figure S19**). This data is in agreement with a previous kinetic analysis of DtpM, showing that the mutations His251A and Glu310A lead to reduced catalytic activity.^8^ In summary, Asn252 is the catalytic relevant residue in DptM and His251 as well as Glu310 only contribute to a minor extent to substrate deprotonation. In agreement, sequence alignments indicate that His251, Asn252, and Glu310 are highly conserved in DtpM, LaPhzM^16^, and other related *O-*MTs underlining the results from the mutagenesis screening^8^.

### *In silico* modelling of XrdM

Since the crystal packing of XrdM prevented ligand binding *in crystallo* and the structure of DtpM was too distinct from XrdM (**Figure S14c**) to simply model substrate/product by superposition, we performed *in silico* modeling with XrdM using its AlphaFold2 prediction (**Figure S14a**). Furthermore, the ToxA structure was used as a template and several restraints were applied (**Figure S20d**). This way we obtained a structural model of XrdM with SAH and XRD-271Me (**Figure S20b, c**), based on which we generated a series of mutants (Y7A, Y7F, Y14A, Y18A, Q23A, F112A, F112L, H115A, Y116F, and H238A), targeting residues protentially involved in ligand binding and catalysis (**Figure 5 and Figure S21**). According to the model, His238 and Gln23 might hydrogen bond via a water molecule to the amide oxygen atom of the aliphatic tail of XRD-271Me. Notably, Gln23 is also involved in substrate binding by ToxA (**Figure S20e**)^22^. Mutation of either His238 or Gln23 to alanine slightly reduced XrdM activity, implying that both residues are indeed relevant for substrate recognition (**Figure 5 and Figure S21**). However, similarly to DptM, the polar contacts between the enzyme and the ligand’s tail are not dominant. We therefore probed key hydrophobic contacts with the dithiopyrrolone ring predicted by the *in silico* modelling. For instance, Tyr18 and Phe112 were supposed to position the ligand by stacking laterally and from top to the disulfide moiety of XrdM-271Me (**Figure S20c**). While mutation of Tyr18 to alanine caused only a very slight reduction in catalytic activity, suggesting that the aromatic character of residue 18 is dispensable, the F112A mutant was more severely compromised in substrate turnover (**Figure 5 and Figure S21**). To evaluate the impact of an aromatic versus an aliphatic side chain, we additionally tested a F112L mutant and found that is was fully active (**Figure 5 and Figure S21**). From these results we conclude that hydrophobic contacts between residue 112 and the dithiopyrrolone ring are crucial for the catalytic activity of XrdM. Support for this notion is also provided by ToxA, where Trp112 stacks from top onto the substrate (**Figure S20e**).^22^ Next, we investigated the impact of Tyr14 that according the model hydrogen bonds to the acceptor nitrogen atom in the dithiopyrrolone ring (**Figure S20c**). The mutation Y14A drastically restricts enzymatic activity, implying that Tyr14 is required for efficient catalysis by either activating the nitrogen atom for methyl transfer or stablizing the transition state, i.e. the positive charge of the transferred methyl-group (**Figure 5 and Figure S21**). Similarly as in DtpM, the carbonyl carbon atom of the dithiopyrrolone ring is embedded in a set of hydrogen bonds with Tyr7, His115 and Tyr116 (**Figure S20c**) and each of the mutations Y7A, H115A and Y116F completely inactivates XrdM (**Figure 5 and Figure S21**). Notably, the mutation Y7F retained significant catalytic activity, implying that the hydrogen bond provided by Tyr7 is of less importance than those mediated by His115 and Tyr116. Similar results have been reported for ToxA^22^: The residues Asn115 and Tyr116 of ToxA, corresponding to His115 and Tyr116 of XrdM, were shown to be involved in substrate coordination (**Figure S20e**). Furthermore, the mutations Y7F and Y7A cause modest and strong reductions respectively in catalytic activity of ToxA.^22^ Considering that the residues 1-23 are disordered in the experimentally determined apo XrdM structure but functionally relevant, as shown by the above listed single point mutants as well as by an catalytically inactive N-terminal truncation mutant (ΔNterm) (**Figure S21**), we suppose that these residues only adopt their correct fold upon ligand binding, similarly to an induced-fit mechanism. Notably, the catalytic importance and conservation of two tyrosine residues – one as part of the mobile N-terminal segment (Tyr7 in XrdM) and a second Tyr (Tyr116 in XrdM) at the active site – has been noted before for other *N*-MTs such as ToxA (**Figure S20e**) and glycine *N*-MT^22,30,31^.

The striking similarity of XrdM with ToxA prompted us to to investigate whether XrdM and ToxA are active against each other’s substrates. To this end, we expressed the *tox*-*like* BGC (except the ToxA- coding gene) from *X. szentirmaii* DSM 16338 in *E. coli*, resulting in the engineered *E. coli* strain to produce 1,6-DDMT (1,6-didemethyltoxoflavin), the substrate of ToxA (**Figure S22a, b**). Subsequently, the genes of *toxA* and *xrdM* were cloned and expressed in 1,6-DDMT-producing strain, respectively (**Table S3, S4, and S7**). LC-MS analysis showed that 1,6-DDMT can be methylated in the ToxA but not XrdM expressing strain (**Figure S22c, d**). Likewise, expression of ToxA in non-methylated XRD- producing strain Δ*hfq* Δ*DC* P*_BAD_*_xrdA did not yield methylated XRDs either (**Figure S12**). These findings point out that the *N*-MTs XrdM and ToxA have been “tailored to function” to participate in specialized metabolic pathways though both of them appear to share high structural identity.

## Discussion

Recent work identified the *N*-MT DtpM of *Sacchrothrix algeriensis* NRRL B-24137 that is involved in thiolutin biosynthesis via BLASTP similarity search^8^. However, a similar strategy for the identification of the XRD MT in *Xenorhabdus doucetiae* FRM16 failed even when DtpM was used as the bait. Using proteomics analysis together with overexpression and knockout studies, we finally identified XrdM located almost 600 kb way from the XRD cluster as the XRD MT. We furthermore noticed that XrdM and DtpM are isofunctional proteins toward XRDs and holomycin^8^ though they can be regarded as non-homologous proteins (14.6% identity). A sequence similarity network analysis (**Figure 3**) showed that XrdM and DtpM are mapped to different subnetworks consisting of Pfam00891 and Pfam13649 proteins, respectively, implying that XrdM and DtpM stem from different Pfam origins. Moreover, their corresponding homologs in each other’s host (namely XrdM vs. CTG1_5739; DtpM vs. MT15) have been shown to be unable to fulfill the same task. These results exemplify that identification of a certain MT simply by a sequence similarity search may be misleading, reflecting that MTs participates in specialized metabolite pathways are likely to be “tailored” in a clade- and sometimes species-specific manner^24^. This is also indicated by the occurrence of DtpM and XrdM homologs in all strains encoding a DTP-producing pathway identified in all strains listed in the antiSMASH database (**Figure S23**), where DtpM and XrdM might be specific to Gram-positive and Gram-negative bacteria, respectively.

To characterize XrdM in comparison to DtpM, we determined the structures of both by X-ray crystallography. Notably, the 3D structures of DtpM and XrdM turned out to be fairly different in their substrate binding domains (**Figure 4a and S14c**). While DtpM is most similar to LaPhzM^16^, XrdM is reminiscent of ToxA^22^. For this reason, complex structures of DtpM with XRD-271 and XRD-271Me could not be used to infer any conclusions about substrate binding in XrdM. Yet, to learn about how XrdM binds XRD-271/XRD-271Me, we performed *in silico* molecular modeling. Both the experimental ligand structures with DtpM and the modeling data for XrdM were validated by site-directed mutagenesis combined with activity assays. From these results we inferred that Tyr14 in XrdM and Asn252 in DtpM represent the key residues that either render the amide nitrogen atom of the dithiopyrrolone ring competent for methyl transfer and/or stabilize the positive charge of the transition state during the S_N_2 reaction. Furthermore, hydrogen bridging to the amide oxygen next to the target nitrogen (His251 in DtpM as well as His115 and Tyr116 in XrdM) enhance the nucleophilicity of the target nitrogen atom by draining off electrons.

With all this structural data in hand, we wondered why the sequence homologues of XrdM, ToxA as well as MT4, MT13 and M14 fail to methylate XRDs, as they all cluster in the same sequence similarity subnetwork (**Figure 3**). We therefore compared the sequences and structures of XrdM and ToxA, as well as XrdM and MT4, MT13 and MT14.

For ToxA, XRD-271 could be too large as a substrate. Considering that ToxA naturally methylates an azapteridine scaffold with no extensions, one could speculate that the aliphatic tail of XRD-271 might be too large to fit into the ToxA active site. In this regard, Val145, Phe187 and Leu237 could pose some steric hindrance to XRD-271 binding. Furthermore, Tyr14 in XrdM, proven to be important for XRD activation (**Figure S21**), is replaced by Phe14 in ToxA. Hence ToxA lacks an obvious base close to the site of methylation, the N6 atom in 1,6-didemethyltoxoflavin (1,6-DDMT).^22^ This observation might imply that either the aromatic character of residue 14, which is retained in both XrdM and ToxA, is required for catalysis by stabilizing the positively charged transition state during methyl transfer or that the electronic configuration of the substrates XRD-271 versus 1,6-DDMT determine whether the enzyme needs a catalytic base for the methyl transfer or not. 1,6-DDMT shares similarities with uracil whose N3 position, flanked by two carbonyl groups, is known to be a weak acid^32,33^. Similarly, the two carbonyl groups next to the N6 atom in 1,6-DDMT might drain off electrons and favor the disscociation of the hydrogen without the need for Tyr14. Activation of the C5 ketogroup by hydrogen bonds with Asn115 and Tyr116 further promotes the deprotonation. Even though in XrdM the amide group of XRD-271 is also activated by hydrogen bonds between the carbonyl oxygen atom and His115 as well as Tyr116, it might still not be acidic enough to allow for a S_N_2 reaction and hence, Tyr14 is required. (**Figure S20a, e-g**).

Next, we inspected the sequences and AlphaFold2^21^ predictions for MT4, MT13 and MT14, and compared them to XrdM. Since MT4, MT13 and MT14 are inactive towards XRD-271, we looked for residues that are in XrdM involved in XRD-271 binding or activation, but missing in all three homologous MTs. This way, we spotted Tyr18 that is in MT4, MT13 and MT14 replaced by Ser18. Mutation of S18Y in MT13 and MT14 however did not activate these MTs toward XRD-271 (**Figure S24**). This result agrees with the fact that exchange of Tyr18 by Ala in XrdM did not impair enzyme activity (**Figure S21**). In addition to Tyr18, MT4 lacks key other residues like His115, Tyr116 and His238. However, even the MT4 quadruple mutant S18Y-C115H-H116Y-D238H did not convert XRD-271, suggesting that other factors like altered geometries, dynamics or charge distributions might play a role. The failure to explain the inactivity of MT4, MT13 and MT14 toward XRD-271 might result from both technical limitations (resolution of X-ray structure/accuracy of the *in silico* modeling and the AlphaFold2 algorithm^34^) as well as the inability to predict activities and substrate specificities from primary or tertiary protein structures.

To summarize, we have utilized a comprehensive approach combining proteomic analysis, *in vivo* expression/screening, and gene deletion to identify the amide MT XrdM of the dithiolopyrrolone antibiotic xenorhabdin. The crystallographic analysis proved that XrdM from the Gram-negative organism *Xenorhabdus doucetiae* FRM16 is completely different from its isofunctional protein DtpM from the Gram-positive organism *Sacchrothrix algeriensis* NRRL B-24137. Structure-based mutagenesis and activity assays of XrdM suggest that this *N*-MT has been “tailored” in a species-specific manner during DTP biosynthesis. Our study provides an interesting example for the parallel/independent evolution of two different enzymes for the same reaction in different organisms and expands the limited knowledge of post-NRPS amide methylation in DTP biosynthesis.

## Methods

### General methods

All reagents and chemicals were obtained from Sigma–Aldrich, ROTH, and BD. Genomic DNA was isolated using the Monarch^®^ Genomic DNA Purification Kit. DNA purification was performed from 1% TAE agarose gel using Monarch^®^ DNA Gel Extraction Kit. Plasmids in *E. coli* were isolated by using Monarch^®^ Plasmid Miniprep Kit. DNA primers were purchased from Sigma-Aldrich. Synthetic genes of *ctg1_5739, hlmA, and actA* were purchased from Twist Bioscience. Synthetic genes of *xrdM* and *dtpM* were purchased from MWG Eurofins Genomics (Ebersberg, Germany). DNA polymerases (Taq and Q5) were purchased from New England Biolabs. Polymerases were used according to the manufacturer’s instructions. Plasmid constructions used for gene expression and deletion were assembled using the NEB Gibson assembly^®^ kit.

HPLC-UV-MS analysis was conducted on an UltiMate 3000 system (Thermo Fisher) coupled to an AmaZonX speed mass spectrometer (Bruker, ESI-MS spectra were recorded in positive-ion-mode with the mass range from 100-1200 *m*/*z,* ultraviolet at 190-800 nm) with an ACQUITY UPLC BEH C-18 column (130 Å, 2.1 mm × 50 mm, Waters) at a flow of 0.4 mL/min (5–95% acetonitrile/water with 0.1% formic acid, v/v, 18 min). Q-TOF HRMS analysis was conducted on an Agilent Infinity II HPLC coupled to an Agilent 6550 Quadrupole Time-Of-Flight (Q-TOP) Mass Spectrometer with an ACQUITY UPLC BEH C18 column (130 Å, 2.1 mm × 50 mm, Waters) at a flow of 0.4 mL/min (5–95% acetonitrile/water with 0.1% formic acid, v/v, 20 min).

Orbitrap HRMS analysis for 1,6-DDMT (1,6-didemethyltoxoflavin) and reumycin were conducted on a Kinetex Evo C18 column (100 Å, 2.1 mm × 150 mm, 1.7 μm particle size, Phenomenex) equipped with a guard column (2.1 mm × 20 mm) of similar specificity at a constant eluent flow rate of 0.2 mL/min with eluent A being 0.1% formic acid in water and eluent B being 0.1% of formic acid in MeOH using the following elution profile: 0 – 4 min constant at 0 % B; 4 – 12 min from 0 to 90 % B; 12 – 15 min constant at 90 % B; 15 – 15.1 min from 90 to 0 % B; 15.1 to 20 min constant at 0 % B. A thermos scientific ID-X Orbitrap mass spectrometer was used in negative mode with a high-temperature electrospray ionization source and the detection was performed in full scan mode using the mass analyzer at the mass resolution of 240000.

### Culture and media

Unless otherwise specified, all strains were cultured in LB medium (10 g/L tryptone, 5 g/L yeast extract, and 5 g/L NaCl) or on an LB agar plate (1.5 % agar was added) at 37 °C (for *E. coli* strains) or 28 °C (for *Xenorhabdus* strains). Antibiotics kanamycin (50 µg/mL) and chloramphenicol (34 µg/mL) were added when appropriate. For the production culture of *X. doucetiae* and its mutants, the overnight LB culture was transferred into 5 mL XPP medium^11^ with a starting OD_600_ of 0.1 and 0.2% of L-arabinose as an inducer if required, as well as selective antibiotics at 28 °C with shaking at 200 rpm for 3 days. For large production of non-methylated XRDs and *N*-methylated XRDs, the overnight LB culture was transferred into 5 x 1000 mL XPP medium (1:100, v/v) supplemented with 2% (v/v) of AmberliteTM XAD-16 resins, 0.2% of L-arabinose as an inducer, and antibiotics at 28 °C with shaking at 200 rpm for 3 days.

### Proteomic analysis

A fresh 5 mL LB culture of *X. doucetiea* wild type and Δ*hfq* were inoculated from an overnight culture at OD_600_ of 0.1. Cells were grown with shaking at 28 °C until the OD_600_ of 4, 750 µL were harvested at 10000 rpm for 1 min. The cell pellets were washed twice with cold PBS buffer at 10000 rpm for 1 min. Afterward, the cell pellets were frozen in liquid nitrogen and stored at –80 °C. Samples were prepared in biological quadruplicates. Procedures for sample processing and HPLC-MS/MS analysis were the same as previously described^35^. For the data presented in **Figure 1d and S4**, the threshold levels to assign the differentially expressed proteins between Δ*hfq* and WT were relative fold change > 2 and the *q*-value < 0.05 based on samples with four biological replicates. Each point in the plot corresponds to the relative fold change between Δ*hfq* and WT for the peptides with minimal *q*-value among all peptides quantified for a certain protein, and thus one protein is represented by a single dot in the graph. Raw data from the proteomic analysis was shown in **Extended Excel Sheet_2**.

### Construction of MT expression strains

The MT candidates (MT1–MT15, listed in **Table S2**) were amplified from the genomic DNA of *X. doucetiae FRM16* wild-type, using forward and reverse primers (listed in **Table S5**) incorporating the overlapping part to vector pACYC. The vector backbone was amplified by using primers CK1007 and CK1010 listed in **Table S5**. The PCR amplicons were ligated with the vector backbone by Gibson assembly. The resulting plasmid pMT1–pMT15 (**Table S4**) was used to transform *E. coli* DH10B, respectively. After verifying the identity of the plasmid by PCR and sequencing, it was extracted from *E. coli* DH10B and transformed into strain Δ*hfq* P*_BAD_*_xrdA^36^, affording the protein expression strain, for example, Δ*hfq* P*_BAD_*_xrdA pMT1 listed in **Table S3**. The successful transformation was verified by antibiotic selection and PCR verification. This protocol also applies to the construction of strains containing pCTG1_5739.

### Construction of gene deletion strains

The following protocol applies to construct mutant Δ*MT3,* but the overall procedure was also used to construct mutants Δ*MT13,* Δ*MT14,* Δ*MT15,* and Δ*xrdA-H.* For the deletion of *MT3*, two homologous arms HAL (<u>H</u>omologous <u>A</u>rms <u>L</u>eft) and HAR (<u>H</u>omologous <u>A</u>rms <u>R</u>ight) flanking of the target region was amplified by using primer pairs LS130 + LS131 and LS132 + LS133 (listed in **Table S5**), respectively. The vector backbone of pEB17 was amplified by using the primer pair LS142 + LS143 (**Table S5**). These 3 PCR fragments were ligated by Gibson assembly, resulting in the plasmid construct pEB_de-MT3 (**Table S4**). The plasmid was electorally transformed into *E. coli* ST18 and then transferred to *X. doucetiae* FRM16 via intergenic conjugation. The resulting conjugants were screened and selected for successful double crossover by primer LS134 + LS135 (**Table S5**). Subsequently, the plasmid pEB_de-MT3 was removed by sucrose (6%) resistance selection. Finally, the successful gene deletion mutant Δ*MT3* was verified by PCR and sequencing.

### Gene replacement of *xrdE* by *hlmA* and *actA*

The method used to construct gene-swapped mutants Δ*hfq* Δ*xrdE::hlmA* and Δ*hfq* Δ*xrdE::hlmA* is similar to the method used for gene deletion, except for one additional insertion gene fragment was inserted between HAL and HAR during the plasmid assembly (**Table S4**). The resulting plasmid was electorally transformed into *E. coli* ST18 and then transferred to strain Δ*hfq* via intergenic conjugation. After screening and plasmid removal, the right *xrdE*-swapped mutant (**Table S3**) was verified by PCR and sequencing.

### Sequence similarity network analysis

The sequences of XrdM and DtpM were used as the query for a search of the UniProt database using BLAST, respectively. An all-by-all BLAST was performed to obtain its homologs from bacteria (filter of taxonomy) with the default BLAST retrieve options (e-value: 5, the maximum number of sequences retrieved: 1000). The resulting homologs sequences for XrdM (923 sequences) and DtpM (999 sequences) were downloaded and combined for further network analysis. The network was constructed by an all-by-all blastp comparison of each sequence against each other sequence and was generated using the EFI-Enzyme Similarity Tool^37^. The resulting SSN was visualized in Cytoscape 3.9.1^38^ using an alignment score cutoff of 70 (up to 331), and sequences of >30% identity.

### MIC measurement of XRD-271 and XRD-285Me

The antimicrobial activity of XRD-271 and XRD-271Me was performed by adapting from the CLSI guidelines M7-A11 for aerobic bacteria for the determination of dilution minimum inhibitory concentrations (MICs). Suspensions equivalent to a McFarland 0.5 standard were prepared for the test strains in medium LB (for *E.coli* MG1655 Δ*tolC* Δ*bam* and *Micrococcus luteus* DSM20030) and YPD (for *Saccharomyces cerevisiae*, YPD recipe for 1 L: Yeast extract 10g, Bacto-peptone 20g, Tryptophan 0.32g, 40% Glucose 50 mL). The inoculum was further diluted 1:200 (for *E. coli*) and 1:10 (for *M. luteus* and *S. cerevisiae* CEN. PK2) in a medium. The diluted suspension was aliquoted into 96-well plates at 100 µL per well. A series of DMSO diluted solutions of XRD-271 and XRD-271Me were prepared (see details in **Extended Excel sheet_1**) and 1 µL of them were added to each well accordingly. Suspension without adding compounds and medium without inoculum was also tested as controls. Assay plates were incubated at 30 °C for 16-40 h. The MIC was defined as the lowest concentration of compound completely inhibiting visible growth after 16 to 24 h of incubation at 30°C. Raw data from the MIC test was shown in **Extended Excel Sheet_1**.

### Isolation and purification of XRD-271 and XRD-285Me

To facilitate the purification of XRDs, the decarboxylase (DC, XDD1_2132) which is responsible for the formation of amides derivatives^39^ was deleted in strain Δ*hfq* P*_BAD_*_*xrdA*, resulting in the mutant Δ*hfq* Δ*DC* P*_BAD_*_*xrdA* (**Table S3**), which has an extremely clear background for the production of non-methylated XRDs. The introduction of pXrdM into this strain resulted in mutant *Δhfq ΔDC* PBAD_xrdA pXrdM (**Table S3**), specifically producing *N*-methylated XRDs.

Mutants Δ*hfq* Δ*DC* P*_BAD_*_*xrdA* and Δ*hfq* Δ*DC* P*_BAD_*_*xrdA* pXrdM were cultured in 5 liters of XPP medium with 2% XAD-16 resins and induced by the addition of L-arabinose, respectively. After incubation at 28 °C with 200 rpm of shaking for 72 h, XAD-16 resins for non-methylated XRDs and methylated XRDs were separately harvested, washed with water, extracted with methanol, and evaporated to obtain the crude extracts. Extracts were subjected to semi-preparative HPLC (see details in **Table S8**), resulting in pure XRD-271 and XRD-271Me of 64.4 mg and 84.2 mg, respectively.

### Heterologous expression of XrdM and DtpM

Synthetic genes encoding XrdM (residues 4-246) and DtpM (**Table S6**) were respectively inserted with the restriction enzymes *Bam*HI and *Pst*I into the multiple cloning site 1 of a pET-DUET-1 derived plasmid encoding an N-terminal His_6_-SUMO-tag (MWG Eurofins Genomics). The resulting vectors served as expression plasmids for the heterologous production of proteins and as templates for in vitro site-directed mutagenesis. DtpM and XrdM mutants were created with Q5 mutagenesis. Primers were designed with the NEBase Changer tool (https://nebasechanger.neb.com/) (**Table S5**). After whole plasmid amplification with Q5 polymerase, a KLD reaction was set up to degrade the template DNA with DpnI and to ligate the amplified linear DNA fragments with the help of T4 polynucleotide kinase and T4 ligase. After 2 hours of incubation at room temperature, 1–10 µL of the KLD reaction were transformed in competent *E. coli* cells. Plasmids were re-isolated and checked for the desired mutation by sequencing (Eurofins Genomics, Ebersberg, Germany)

The corresponding expression plasmid encoding wild type or mutant XrdM or DtpM (**Table S4**) was transferred in *E. coli* BL21-Gold (DE3) and a single colony was picked for overnight cultivation in liquid LB medium supplemented with 180 mg/L ampicillin at 37°C. After inoculation of the main culture (1:50), flasks were shaken at 37 °C until an optical density (OD_600nm_) of 0.5-0.7 was reached. Prior to addition of 0.5 mM isopropyl-*β*-D-1-thiogalactopyranoside (IPTG), the cultures were cooled to 20°C. After overnight expression at 20°C, the cells were harvested by centrifugation, washed with 0.9% (w/v) NaCl, and frozen at –20°C.

### Purification of wild type and mutant XrdM

For protein purification, cell pellets were resuspended in 100 mM Tris/HCl pH 8.0, 300 mM NaCl, 20 mM imidazole and 10% (v/v) glycerol (buffer A) and lysed by sonication. Upon centrifugation at 41,000 x *g* for 30 min at 4 °C, the supernatant was loaded on a 5 mL nickel chelating Sepharose HP column (Cytiva), pre-equilibrated with buffer A (flow rate 5 mL/min). After washing with buffer A, the protein was eluted by increasing the concentration of buffer B (100 mM Tris/HCl pH 8.0, 300 mM NaCl, 500 mM imidazole, and 10% (v/v) glycerol) from 0% to 50% over 15 column volumes (CV) and further to 100% in another 5 CV. Fractions containing XrdM were supplemented with SUMO-protease to remove the His_6_-SUMO-tag and dialyzed overnight against buffer C containing 100 mM Tris/HCl, pH 8.0, 300 mM NaCl, and 10% (v/v) glycerol. The protein was again loaded on a 5 mL nickel chelating Sepharose HP column, pre-equilibrated with buffer A to remove the proteases and the cleaved tag. The flow through, containing the protein of interest, was collected, dialyzed against 20 mM Tris/HCl, pH 8.0, 100 mM NaCl, and 1 mM DTT (buffer D), concentrated, and subjected to size exclusion chromatography (Superdex 75 16/60 for preparative scale, GE Healthcare) using buffer D as running buffer.

### Purification of wild type and mutant DtpM

Cells were thawed in buffer E (100 mM Tris/HCl pH 7.5, 250 mM NaCl, 20 mM imidazole) and disrupted by sonication. After removal of cell debris by centrifugation at 41,000 x *g* for 20 min at 4°C, the cleared lysate was applied to a 5 mL nickel chelating Sepharose HP column (Cytiva), pre-equilibrated with buffer E (flow rate 5 mL/min). Washing with buffer E removed non-specifically bound proteins. The protein of interest was eluted by applying a linear gradient from 0% to 100% buffer F (100 mM Tris/HCl pH 7.5, 250 mM NaCl, 500 mM imidazole) over 10 CV, supplemented with SUMO-protease and dialyzed against buffer G (20 mM Tris/HCl pH 7.5, 200 mM NaCl) at 4°C. A reverse Ni-affinity chromatography finally separated cleaved from uncleaved, protein, tag, and protease. Cleaved protein was concentrated with a 30 kDa cutoff filter (Amicon) and further purified by size exclusion chromatography using a Superdex 200 16/60 column and buffer H (20 mM Tris/HCl pH 7.5, 100 mM NaCl).

### Crystallization and structure determination of apo XrdM

According to the results of the differential scanning fluorimetry (**Figure S16**), DtpM was 3 °C more stable in HEPES buffer pH 7.0 and with 10% glycerol. We therefore dialyzed the protein prior to crystallization trials against 20 mM HEPES pH 7.0, 100 mM NaCl and 10% (v/v) glycerol. Diffraction-quality crystals of apo XrdM were obtained by the sitting drop vapour diffusion technique at 20°C and by mixing equal volumes of protein (∼28 mg/mL, supplemented with 2 mM SAM) and reservoir solution (0.1 M Bicine pH 9.0, 30% (w/v) PEG6000. Crystals were harvested by the addition of a 1:1 (v/v) mixture of reservoir solution and 60% (v/v) glycerol and vitrified in liquid nitrogen.

Diffraction data were collected at the beamline X06SA, Swiss Light Source (SLS), Villigen, Switzerland. Reflection intensities were analyzed with the program package XDS.^40^ The structure of XrdM was solved by molecular replacement with Phaser^41^ (CCP4 suite) using the AlphaFold2^21^ prediction (https://colab.research. google.com/github/sokrypton/ColabFold/blob/main/AlphaFold2.ipynb) of XrdM as a search model. After iterative model building and refinement steps with Coot^42^ and REFMAC5,^43^ water molecules were placed with ARP/wARP solvent.^44^ Translation/libration/screw refinements finally yielded excellent values for R_crys_ and R_free_ as well as root-mean-square deviation (r.m.s.d.) bond and angle values. The models were proven to fulfill the Ramachandran plot using PROCHECK^45^ (**Table S9**) and evaluated by MolProbity.^46^

### Co-crystallization and soaking attempts of XrdM with either XRD271 or XRD-271Me

To prevent XRD-271 and XRD-271Me from reduction, XrdM was purified without reductive agent. Purified XrdM was supplemented with 2 mM SAH and 2 mM XRD-271 or XRD-271Me, both dissolved in DMSO, and crystallized by the sitting drop vapor diffusion technique at 20°C. Besides colorless crystals, several yellowish ones grew. They were vitrified as the apo crystals and several diffraction datasets were obtained up to 2.25 Å resolution, but none of them visualized the ligand.

### Crystallization and structure determination of apo DtpM

Despite high yield and purity of the DtpM sample, initial crystallization trials with SAM alone failed. Analysis of the thermal stability of DtpM under different buffer conditions using differential scanning fluorimetry indicated that the protein was robustly folded (melting temperature (T_M_) of 44.5 °C, **Figure S16**). However, the melting temperature could be further increased by lowering the pH and addition of glycerol (T_M_ of 49.5 °C). We therefore dialyzed the protein against an optimized buffer and set up new crystallization trials. Apo DtpM (∼18 mg/mL, supplemented with 2 mM SAM) crystallized from sitting drop vapor diffusion crystallization trials at 20°C and a reservoir solution of 2.1 M D/L-malic acid pH 7.0 and a protein: reservoir ratio of 1:1. Crystals were cryoprotected by the addition of a 1:1 (v/v) mixture of reservoir solution and 60% (v/v) glycerol.

Diffraction data, collected at the beamline X06SA, Swiss Light Source (SLS), Villigen, Switzerland, were evaluated with the program package XDS.^40^ Patterson search calculations using the AlphaFold2^21^ model of DtpM (https://colab.research.google.com/github/sokrypton/ColabFold/blob/main/ AlphaFold2.ipynb) initially did not yield a molecular replacement solution with Phaser^41^ (CCP4 suite). Only a truncated model (residues 97-346) could be positioned in the crystal lattice. The missing domain was afterwards automatically built using ARP/wARP classic (CCP4)^47^ and optimized by iterative model building with Coot^42^ and refinement with REFMAC5^43^ Water molecules were placed with ARP/wARP solvent^44^. Translation/libration/screw refinements finally polished the structure resulting in good geometric and quality matrices (**Table S9**). The final model was evaluated by MolProbity.^46^

### Crystallization and structure determination of DtpM:SAH:XRD-271 and DtpM:SAH:XRD-271Me

Purified DtpM was supplemented with 2 mM SAH and 2 mM XRD-271 or 2 mM XRD-271Me. Sitting drop crystallization trials at 20°C yielded yellowish crystals under several conditions. Cryoprotection was performed as for apo crystals. Diffraction data for XRD-271Me bound DtpM was collected from a crystal grown from 2.1 M D/L-malic acid pH 7.0 and for XRD-271 bound DtpM from 1 M LiCl_2_, 0.1 M MES pH 6.0, 30% (w/v) PEG6000. Reflection intensities were analyzed with XDS^48^ and the structures were solved by molecular replacement (PHASER^41^ within the CCP4 suite) using the coordinates of apo DtpM. The F_O_-F_C_ and 2F_O_-F_C_ omit maps clearly indicated electron density for the cofactor and the ligands (**Figure 4c**). The chemical structures of XRD-271 and XRD-271Me were built with SYBYL 8.0 ( Tripos International, 1699 South Hanley Rd., St. Louis, Missouri, 63144, USA) and restraints were generated using the prodrgify-this-residue tool within Coot^42^. Refinement and validation of the structural models was performed as described for the apo structure.

### Molecular modeling

To obtain an initial model, docking was performed using AutoDock Vina (version 1.2.0)^49,50^. Since the crystal structure of XrdM did not resolve the N-terminus, that likely forms important contacts with the ligand, the related ToxA structure (PDB 5JE1) was used as a receptor. The original ligand as well as water were removed, and protonation states were assigned at pH 7.4 using the PROPKA function within Maestro (version 13.3). XRD-271Me was prepared in its neutral form from the SMILES representation using RDKit (version 2023.03.1)^51^. The docking was performed using a 20x20x20 grid-box set around the center of the original ToxA-ligand and default exhaustiveness settings. Results were subsequently filtered based on the distance of methyl donor and methyl acceptor, i.e. only poses with a distance less than 6 Å were included.

The XRD-271Me pose resulting from cross-docking was then placed into the receptor. We opted to use the AlphaFold2 prediction^21^ of XrdM in our model to account for the disordered N-terminus as well as possible defects in the flap domain compared to related enzymes such as ToxA due to artificial dimerization in the XrdM crystal. Next, the model was manually refined with reference to the ToxA cocrystal structure. In particular, the rotation of the dithiolopyrrolone core was adjusted to reflect pi-stacking with Phe112, His238 was rotated and a water-molecule was modeled in the active site instead based on the ToxA structure (PDB 5JE1). Finally, the aliphatic tail was repositioned using Maestro’s built-in minimization feature.

### Graphical illustrations of X-ray structures

Graphical illustrations were created with PyMOL.^52^

### Differential scanning fluorimetry

DtpM was tested for its thermostability under different conditions in a 96-well thin-wall PCR plate (ThermoFisher). Each well contained 1 µl fluorescent dye indicator SYPRO Orange (Sigma Aldrich, 5000 x in DMSO, 1:40 diluted with H_2_O), 1 µl of 5 mg/ml DptM and 18 µl of buffer. The plate was sealed, centrifuged and then heated in a Bio-Rad CFX96 Real-Time PCR Detection System from 4 °C to 95°C in increments of 0.5 °C/20 sec, and the fluorescence was monitored. The results were normalized and tentatively fitted with GraphPad Prism 5. Data points before the minimum and after the maximum of the fluorescence intensity were excluded from fitting.^53^

### In vitro methylation assay of wild type and mutant XrdM/DtpM

The in vitro enzymatic reaction was performed using 100 µM XRD-271, 200 µM SAM, and 100 µM purified protein in Tris/HCl Buffer (20 mM Tris/HCl, 100 mM NaCl, pH 8.0 for wild type and mutant XrdM, pH 7.5 for wild type and mutant DtpM) at 30 °C for 30min or 60min. The reactions were quenched by adding an equal volume of cold MeOH, centrifuged at 13000 RPM for 20 min and subjected to LC-MS analysis.

### Phyletic distribution of XRD-like BGC and MT XrdM/DtpM

A cblaster database from all bacterial genomes in the antismash DB v4 (beta 2)^54^. XRD-like clusters were searched via cblaster^55^ using four separate query clusters of holomycin: MiBIG^56^ entries BGC0000373, BGC0002091, BGC0002412, and a detected cluster in *X. doucetiae* (GCF_000968195.1). The “required” flag was used to indicate homologs of XrdA-D (conserved 4 genes across all DTP producers in **Figure S1**) and all resulting hits were combined to form detected clusters in the database. Homologs of XrdM and DtpM were found via diamond with the “ultra-sensitive” option using the same database generated in cblaster and using XrdM and DtpM as queries. The same was performed for genomes that were not present in the antismash DB. All genomes that showed homologs of either cluster or XrdM/DtpM were collected and placed onto a GTDB tree using the GTDB-Tk^57^ command line tool. One genome, *P.* sp_SANK_73390, was placed at the genus level due to a lack of marker genes as indicated with the “g_” prefix. The resulting tree was pruned in dendroscope^58^ to highlight the relevant genomes and finalized in the Interactive Tree of Life (iTOL)^59^.

### In vivo methylation assay towards non-methylated toxoflavin 1,6-DDMT

The genes of *toxE* and *toxDBC* from *X. szentirmaii* were PCR amplified and fused together before being constructed into plasmid pCOLA (**Table S4**). To produce non-methylated toxoflavin 1,6-DDMT, *E. coli* strain containing pCOLA_ToxEDBC was cultivated in LB culture and supplemented with L-arabinose at a final concentration of 0.2% to activate the production. To test the methylation towards 1,6-DDMT, pToxA (**Table S4**) and pXrdM were separately co-expressed with pCOLA_ToxEDBC in *E.coli* TOP10. After 3 days of incubation at 25 °C, the culture broth was centrifuged at 6000 rpm for 10min. The supernatant was collected and dried by Biotage V-10 Touch evaporation system. The resulting residue was dissolved in H_2_O for LC-MS analysis.

### Site directed mutagenesis of MT13, MT14, and MT4 and *in vivo* methylation assay of resulting mutants towards non-methylated XRDs

The site directed mutagenesis mutants of MT4, MT13, and MT14 were constructed by PCR-mediated mutagenesis, as follows (exemplified by mutant pMT13-S18Y): 1) PCR amplification of the entire plasmid pMT13 (listed in **Table S4**) using primers in which the Ser codon (TCC) was substituted by that of Tyr (TAC) (**Table S5**, LS241 and LS242); 2) digestion of the PCR product with *Dpn*I restriction enzyme for elimination of the methylated DNA template, thereafter, 2 µL digestion mixture was used to transform competent cell of *E. coli* DH10B; 3) at least 3 positive colonies were picked out and cultivated in LB with chloramphenicol for plasmids extraction. The extracted plasmids were verified by DNA sequencing for the accuracy of the mutagenesis and then the right one was transformed into strain Δ*hfq* Δ*DC* P*_BAD_*_xrdA^36^, resulting the strain Δ*hfq* Δ*DC* P*_BAD_*_xrdA pMT13-S18Y (**Table S3**). The quadruple (S18Y-C115H-H116Y-D238H) site mutagenesis mutant of pMT4 were achieved step by step using this method with primers listed in **Table S4**.

## Data availability

Protein coordinates have been deposited in the RCSB Data Bank under the accession code 8RDL (XrdM:SAH), 8RDM (DptM:SAH), 8RDN (DptM:SAH:XRD-271), 8RDO (DtpM:SAH:XRD-271Me).

## Acknowledgments

Work in the Bode Group was supported by an ERC Advanced Grant (835108) and the Max-Planck-Society. E.M.H. and M.G. acknowledge financial support by the DFG – SFB 1309–325871075. We thank the staff of the beamline X06SA at the Paul-Scherrer-Institute, Swiss Light Source, Villigen Switzerland for assistance during data collection. The research leading to these results has received funding from the European Community’s Seventh Framework Programme (FP7/2007-2013) under BioStruct-X (grant agreement N°283570). We are grateful to K. Gärtner for technical assistance and to the students F. Reinhardt, Victoria Gebler and M. Hofberger for experimental support.

## Authors contributions

L.S. performed *in vivo* expression and screening, *in vitro* acticity assays, SSN analysis, generated and fermented recombinant strains, analyzed and interpreted HPLC-MS data. E.M.H. produced XrdM and DtpM proteins and together with M.G. performed crystallographic analysis. M.W. and E.B. contructed selected deletion mutants. L.S., M.W. and T.G. performed proteomic analysis. J.G. performed molecular modeling study. L.S., T.K. and D.S. performed the MIC assay. P.G. purified XRD-271 and XRD-271Me.

M.M.A performed phyletic distribution anlaysis. L.S. and E.M.H. wrote the manuscript, with contributions from all co-authors. M.G. and H.B.B supervised the project.

## Additional information

There are 3 Supplementary files accompany this manuscript: 1) Supplementary information (Figures S1–S24, and Tables S1-S11); 2) Extended_Excel_sheet_1; 3) Extended_Excel_sheet_2.

## Reference

1. Li, B., Wever, W.J., Walsh, C.T. & Bowers, A.A. Dithiolopyrrolones: biosynthesis, synthesis, and activity of a unique class of disulfide-containing antibiotics. Natural Product Reports 31, 905–923 (2014).

2. Li, B. & Walsh, C.T. Identification of the gene cluster for the dithiolopyrrolone antibiotic holomycin in Streptomyces clavuligerus. Proc Natl Acad Sci U S A 107, 19731–5 (2010).

3. Qin, Z., Huang, S., Yu, Y. & Deng, H. Dithiolopyrrolone natural products: isolation, synthesis and biosynthesis. Mar Drugs 11, 3970–97 (2013).

4. Oliva, B., O’Neill, A., Wilson, J.M., O’Hanlon, P.J. & Chopra, I. Antimicrobial properties and mode of action of the pyrrothine holomycin. Antimicrob Agents Chemother 45, 532–9 (2001).

5. McInerney, B.V. et al. Biologically active metabolites from Xenorhabdus spp., Part 1. Dithiolopyrrolone derivatives with antibiotic activity. J Nat Prod 54, 774–84 (1991).

6. Scharf, D.H., Habel, A., Heinekamp, T., Brakhage, A.A. & Hertweck, C. Opposed Effects of Enzymatic Gliotoxin N- and S-Methylations. Journal of the American Chemical Society 136, 11674–11679 (2014).

7. Duell, E.R. et al. Sequential Inactivation of Gliotoxin by the S-Methyltransferase TmtA. ACS Chemical Biology 11, 1082–1089 (2016).

8. Chen, X., Johnson, R.M. & Li, B. A Permissive Amide N-Methyltransferase for Dithiolopyrrolones. ACS Catalysis 13, 1899–1905 (2023).

9. Bode, E. et al. Simple “on-demand” production of bioactive natural products. Chembiochem 16, 1115–9 (2015).

10. Tobias, N.J. et al. Natural product diversity associated with the nematode symbionts Photorhabdus and Xenorhabdus. Nat Microbiol 2, 1676–1685 (2017).

11. Bode, E. et al. Promoter Activation in Δhfq Mutants as an Efficient Tool for Specialized Metabolite Production Enabling Direct Bioactivity Testing. Angew Chem Int Ed Engl 58, 18957–18963 (2019).

12. Bode, E. et al. easyPACId, a Simple Method for Induced Production, Isolation, Identification, and Testing of Natural Products from Proteobacteria. Bio Protoc 13, e4709 (2023).

13. Saker, S., Lebrihi, A. & Mathieu, F. Identification of two putative acyltransferase genes potentially implicated in dithiolopyrrolone biosyntheses in Saccharothrix algeriensis NRRL B- 24137. Appl Biochem Biotechnol 173, 787–802 (2014).

14. Saker, S., Chacar, S. & Mathieu, F. The final acylation step in aromatic dithiolopyrrolone biosyntheses: identification and characterization of the first bacterium N-benzoyltransferase from Saccharothrix algeriensis NRRL B-24137. Enzyme Microb Technol 72, 35–41 (2015).

15. Lamari, L. et al. New dithiolopyrrolone antibiotics from Saccharothrix sp. SA 233. I. Taxonomy, fermentation, isolation and biological activities. J Antibiot (Tokyo) 55, 696–701 (2002).

16. Jiang, J. et al. Functional and Structural Analysis of Phenazine O-Methyltransferase LaPhzM from Lysobacter antibioticus OH13 and One-Pot Enzymatic Synthesis of the Antibiotic Myxin. ACS Chem Biol 13, 1003–1012 (2018).

17. Zhu, Y. et al. Identification of caerulomycin A gene cluster implicates a tailoring amidohydrolase. Org Lett 14, 2666–9 (2012).

18. Ogasawara, Y. & Liu, H.-w. Biosynthetic Studies of Aziridine Formation in Azicemicins. Journal of the American Chemical Society 131, 18066–18068 (2009).

19. Tsodikov, O.V., Hou, C., Walsh, C.T. & Garneau-Tsodikova, S. Crystal structure of O- methyltransferase CalO6 from the calicheamicin biosynthetic pathway: a case of challenging structure determination at low resolution. BMC Struct Biol 15, 13 (2015).

20. Hillwig, M.L. et al. Identification and characterization of a welwitindolinone alkaloid biosynthetic gene cluster in the stigonematalean Cyanobacterium Hapalosiphon welwitschii. Chembiochem 15, 665–9 (2014).

21. Jumper, J. et al. Highly accurate protein structure prediction with AlphaFold. Nature 596, 583–589 (2021).

22. Fenwick, M.K., Philmus, B., Begley, T.P. & Ealick, S.E. Burkholderia glumae ToxA Is a Dual- Specificity Methyltransferase That Catalyzes the Last Two Steps of Toxoflavin Biosynthesis. Biochemistry 55, 2748–59 (2016).

23. Lee, Y.-R. et al. Structural Analysis of Glycine Sarcosine N-methyltransferase from Methanohalophilus portucalensis Reveals Mechanistic Insights into the Regulation of Methyltransferase Activity. Scientific Reports 6, 38071 (2016).

24. Liscombe, D.K., Louie, G.V. & Noel, J.P. Architectures, mechanisms and molecular evolution of natural product methyltransferases. Nat Prod Rep 29, 1238–50 (2012).

25. Laurino, P. et al. An Ancient Fingerprint Indicates the Common Ancestry of Rossmann-Fold Enzymes Utilizing Different Ribose-Based Cofactors. PLOS Biology 14, e1002396 (2016).

26. Krissinel, E. & Henrick, K. Detection of Protein Assemblies in Crystals. CompLife 2005 3695, 163–174 (2005).

27. Sun, Q., Huang, M. & Wei, Y. Diversity of the reaction mechanisms of SAM-dependent enzymes. Acta Pharm Sin B 11, 632–650 (2021).

28. Holm, L. & Rosenstrom, P. Dali server: conservation mapping in 3D. Nucleic acids research 38, W545–9 (2010).

29. Liscombe, D.K., Louie, G.V. & Noel, J.P. Architectures, mechanisms and molecular evolution of natural product methyltransferases. Natural product reports 29, 1238–50 (2012).

30. Takata, Y. et al. Catalytic mechanism of glycine N-methyltransferase. Biochemistry 42, 8394–402 (2003).

31. Lee, S.G., Kim, Y., Alpert, T.D., Nagata, A. & Jez, J.M. Structure and reaction mechanism of phosphoethanolamine methyltransferase from the malaria parasite Plasmodium falciparum: an antiparasitic drug target. J Biol Chem 287, 1426–34 (2012).

32. Ganguly, S. & Kundu, K.K. Protonation/deprotonation energetics of uracil, thymine, and cytosine in water from e.m.f./spectrophotometric measurements. Canadian Journal of Chemistry 72, 1120–1126 (1994).

33. Ilyina, M.G., Khamitov, E.M., Ivanov, S.P., Mustafin, A.G. & Khursan, S.L. Anions of uracils: N1 or N3? That is the question. Computational and Theoretical Chemistry 1078, 81–87 (2016).

34. Terwilliger, T.C. et al. AlphaFold predictions are valuable hypotheses and accelerate but do not replace experimental structure determination. Nature Methods (2023).

35. Neubacher, N. et al. Symbiosis, virulence and natural-product biosynthesis in entomopathogenic bacteria are regulated by a small RNA. Nat Microbiol 5, 1481–1489 (2020).

36. Zhai, Y. et al. Identification of an unusual type II thioesterase in the dithiolopyrrolone antibiotics biosynthetic pathway. Biochem Biophys Res Commun 473, 329–335 (2016).

37. Zallot, R., Oberg, N. & Gerlt, J.A. The EFI Web Resource for Genomic Enzymology Tools: Leveraging Protein, Genome, and Metagenome Databases to Discover Novel Enzymes and Metabolic Pathways. Biochemistry 58, 4169–4182 (2019).

38. Shannon, P. et al. Cytoscape: a software environment for integrated models of biomolecular interaction networks. Genome Res 13, 2498–504 (2003).

39. Bode, E. et al. Biosynthesis and function of simple amides in Xenorhabdus doucetiae. Environ Microbiol 19, 4564–4575 (2017).

40. Kabsch, W. Automatic processing of rotation diffraction data from crystals of initially unknown symmetry and cell constants. J. Appl. Cryst. 26, 795–800 (1993).

41. McCoy, A.J., et al. Phaser crystallographic software. J. Appl. Cryst. 40, 658–674 (2007).

42. Emsley, P., Lohkamp, B., Scott, W.G. & Cowtan, K. Features and development of Coot. Acta Crystallogr. Sect. D - Biol. Crystallogr. 66, 486–501 (2010).

43. Vagin, A.A. et al. REFMAC5 dictionary: organization of prior chemical knowledge and guidelines for its use. Acta Crystallogr. Sect. D - Biol. Crystallogr. 60, 2184–95 (2004).

44. Perrakis, A., Sixma, T.K., Wilson, K.S. & Lamzin, V.S. wARP: improvement and extension of crystallographic phases by weighted averaging of multiple-refined dummy atomic models. Acta crystallographica. Section D, Biological crystallography 53, 448–55 (1997).

45. Laskowski, R.A., Moss, D.S. & Thornton, J.M. Main-chain bond lengths and bond angles in protein structures. Journal of molecular biology 231, 1049–67 (1993).

46. Chen, V.B. et al. MolProbity: all-atom structure validation for macromolecular crystallography. *Acta crystallographica. Section D*, Biological crystallography 66, 12–21 (2010).

47. Langer, G., Cohen, S.X., Lamzin, V.S. & Perrakis, A. Automated macromolecular model building for X-ray crystallography using ARP/wARP version 7. Nat Protoc 3, 1171–9 (2008).

48. Kabsch, W. XDS. Acta Crystallogr. Sect. D - Biol. Crystallogr. 66, 125–32 (2010).

49. Eberhardt, J., Santos-Martins, D., Tillack, A.F. & Forli, S. AutoDock Vina 1.2.0: New Docking Methods, Expanded Force Field, and Python Bindings. J Chem Inf Model 61, 3891–3898 (2021).

50. Trott, O. & Olson, A.J. AutoDock Vina: improving the speed and accuracy of docking with a new scoring function, efficient optimization, and multithreading. J Comput Chem 31, 455–61 (2010).

51. RDKit: Open-source cheminformatics. https://www.rdkit.org

52. DeLano, W.L. The PyMOL Molecular Graphics System. *DeLano Scientific, San Carlos, CA*, USA. (2002).

53. Niesen, F.H., Berglund, H. & Vedadi, M. The use of differential scanning fluorimetry to detect ligand interactions that promote protein stability. Nature protocols 2, 2212–21 (2007).

54. Blin, K., Shaw, S., Medema, M.H. & Weber, T. The antiSMASH database version 4: additional genomes and BGCs, new sequence-based searches and more. Nucleic Acids Res (2023).

55. Gilchrist, C.L.M., et al. cblaster: a remote search tool for rapid identification and visualization of homologous gene clusters. Bioinform Adv 1, vbab016 (2021).

56. Terlouw, B.R. et al. MIBiG 3.0: a community-driven effort to annotate experimentally validated biosynthetic gene clusters. Nucleic Acids Res 51, D603–d610 (2023).

57. Chaumeil, P.A., Mussig, A.J., Hugenholtz, P. & Parks, D.H. GTDB-Tk: a toolkit to classify genomes with the Genome Taxonomy Database. Bioinformatics 36, 1925–7 (2019).

58. Huson, D.H. et al. Dendroscope: An interactive viewer for large phylogenetic trees. BMC Bioinformatics 8, 460 (2007).

59. Letunic, I. & Bork, P. Interactive Tree Of Life (iTOL) v5: an online tool for phylogenetic tree display and annotation. Nucleic Acids Res 49, W293–w296 (2021).

