## Supplementary Information for "Identification, structure and function of the methyltransferase involved in the biosynthesis of the dithiolopyrrolone antibiotic xenorhabdin"

[#] These authors contributed equally

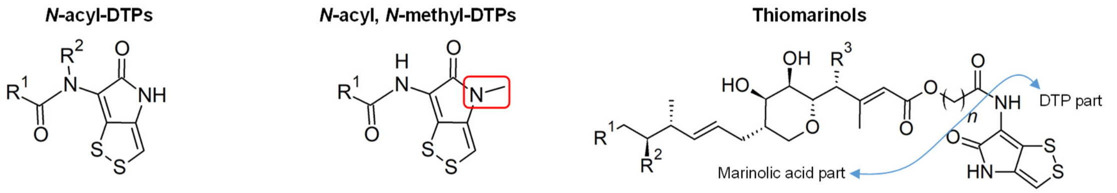


| | R1 | R2 |
|---|---|---|
| Holomycin | $CH_3$ | H |
| VD-844 | H | $CH_3$ |
| XRD-243 | $(CH_2)_2CH_3$ | H |
| XRD-271 | $(CH_2)_4CH_3$ | H |
| XRD-285 | $(CH_2)_3CH(CH_3)_2$ | H |
| XRD-299 | $(CH_2)_6CH_3$ | H |

| | R1 |
|---|---|
| Thiolutin | $CH_3$ |
| Aureothricin | $CH_2CH_3$ |
| XRD-243Me | $(CH_2)_2CH_3$ |
| XRD-271Me | $(CH_2)_4CH_3$ |
| XRD-285Me | $(CH_2)_3CH(CH_3)_2$ |
| XRD-299Me | $(CH_2)_6CH_3$ |

| | R1 | R2 | R3 | n |
|---|---|---|---|---|
| Thiomarinol A | H | OH | OH | 7 |
| Thiomarinol C | H | OH | H | 7 |
| Thiomarinol D | $CH_3$ | OH | H | 7 |
| Thiomarinol E | H | OH | OH | 9 |
| Thiomarinol F | H | =O | OH | 7 |

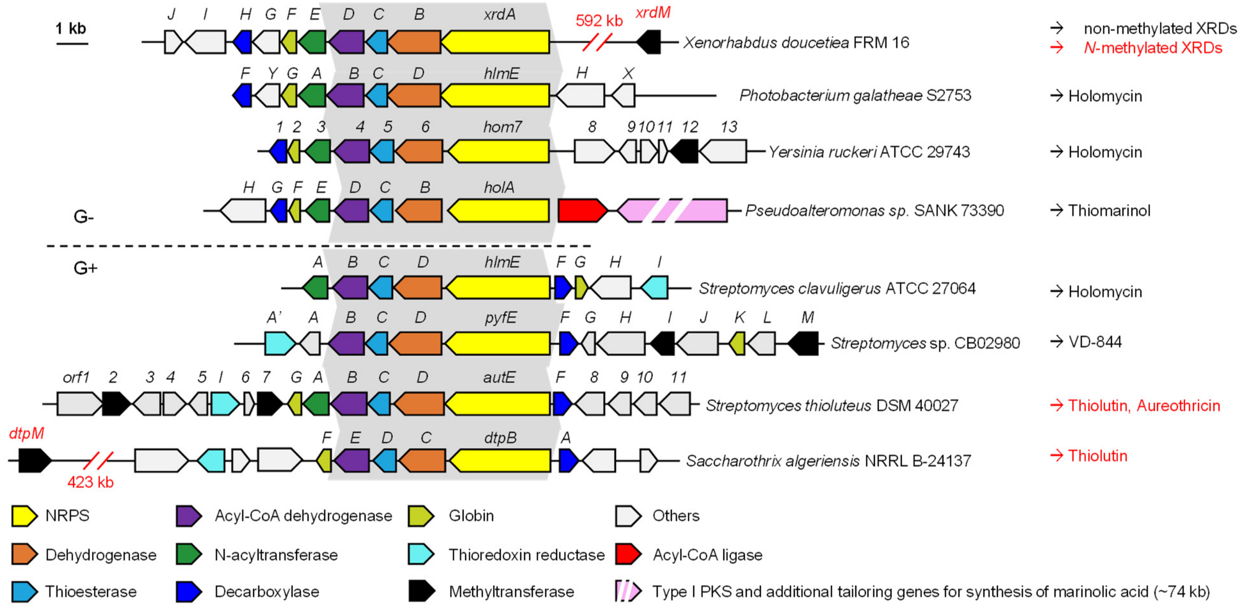


**Figure S1.** Summary of the structures, producers, and biosynthetic gene clusters (BGCs) of naturally occurring dithiolopyrrolones (DTPs). The BGC of Thiomarinol includes two parts: NRPS part for the synthesis of pyrrothine (*holA-H*), and type I PKS part for synthesis of marinolic acid which is not dedicated here.

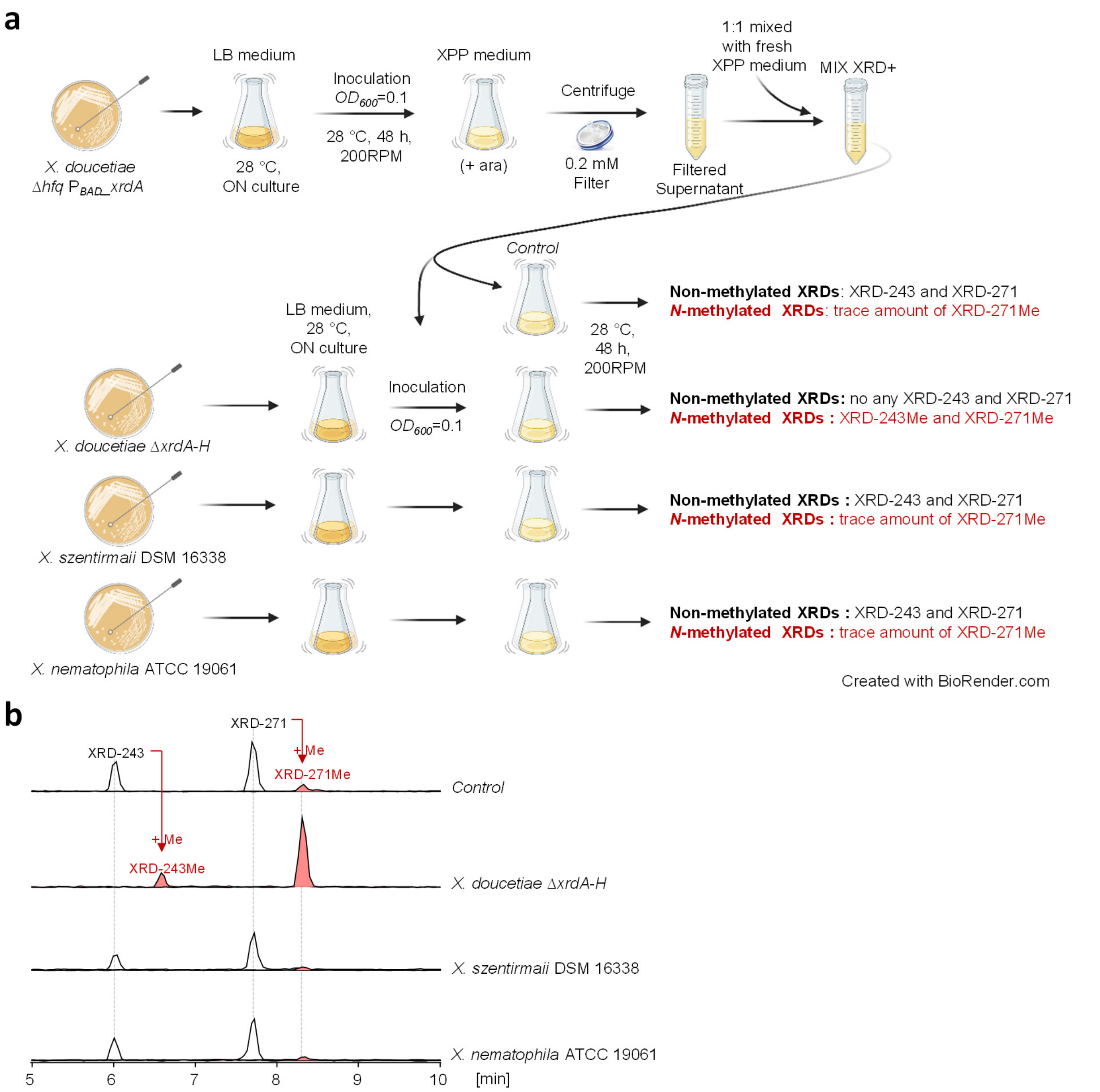


**Figure S2.** Feeding experiments of non-methylated XRDs toward *X. nematophila* and *X. szentirmaii*. (a) Workflow of the feeding experiments. Non-methylated XRDs (XRD-243 and XRD-271) and tiny amount of *N*-methylated XRD-271Me were produced by *X. doucetiae Δhfq* $P_{BAD}$*_xrdA* (a strain created previously following the easyPACiD approach[1] listed in **Tabel S3**) in XPP medium with arabinose added as the inducer. After 2 days, the culture was centrifuged, filtered, and mixed (1 to 1 ratio) with fresh XPP medium. The resulting mixture (MIX XRD+) was used for the production culture of Δ*xrdA-H* (XRD BGC deletion mutant)*, X. szentirmaii* DSM 16338, and *X. nematophila* ATCC 19061. The cultures were sampled and tested by LC-MS after 2 days. (**b**) Extracted ion chromatograms (EIC) from the LC-MS analysis of XRDs in culture extracts of these tested strains. In comparison to the control (no strain inoculated to MIX XRD+), XRD-243 and XRD-271 were completely converted to XRD-243Me and XRD-271Me in Δ*xrdA-H,* but remained unchanged in cultures of *X. szentirmaii* DSM 16338 and *X. nematophila* ATCC 19061, indicating that the latter two strains do not have or do not express the *N*-methyltransferase for XRDs.

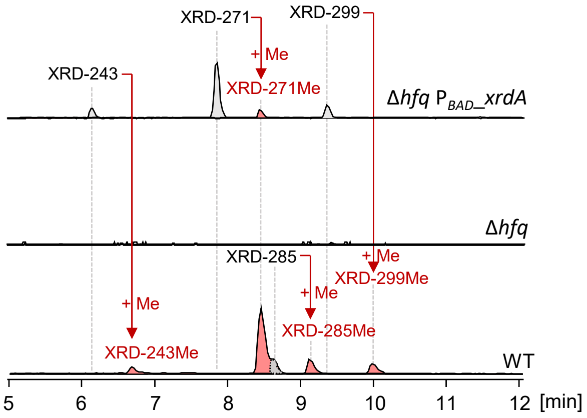


**Figure S3.** The XRD production profile of *X. doucetiae* WT, Δ*hfq,* and Δ*hfq* $P_{BAD}$_xrdA. Four *N*-methylated-XRD derivatives characterized by their different *N*-acyl groups such as XRD-243Me (butyl), XRD-271Me (hexanoyl), XRD-285Me (5-methyl hexanoyl), and XRD-299Me (octanoyl) were observed in WT. Production of XRDs was completely abolished after the deletion of *hfq*. $P_{BAD}$_*xrdA* activation in mutant Δ*hfq* rescues the production of XRDs but without methylation. In addition, XRD-285, which has a branched acyl chain, was not recovered by activation of $P_{BAD}$_*xrdA,* because the deletion of *hfq* disrupts iso-fatty acid formation[2].

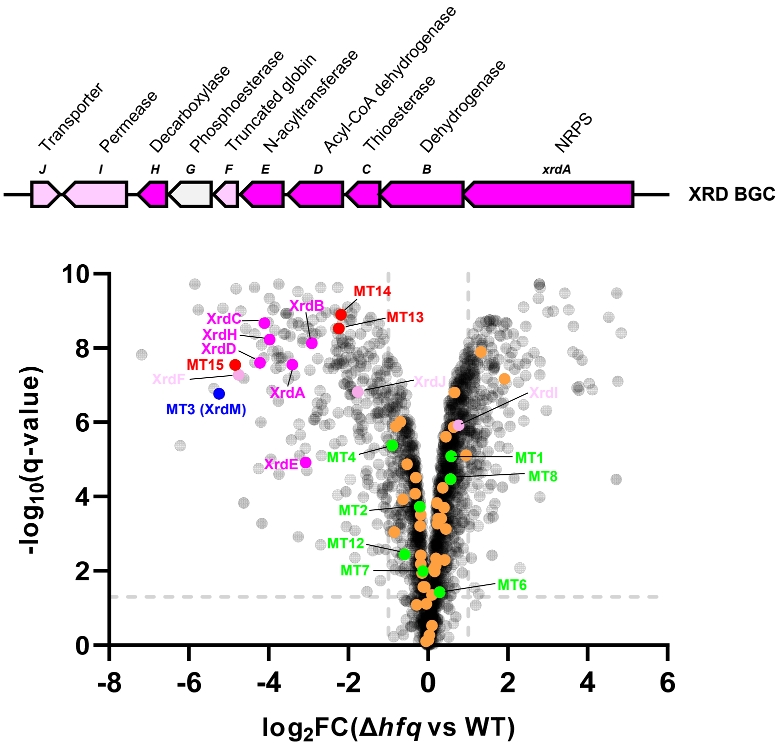


**Figure S4.** Volcano plot of the comparative proteomics analysis between *X. doucetiae Δhfq* mutant and WT. Two cutoffs (dashed lines) shown in the figure were q = 0.05, and FC (fold change) = 2. Data were calculated from four biological replicates. Dots in light and dark pink refer to proteins encoded by the XRD BGC shown above the volcano plot. The dark pink dots were XRDs biosynthesis proteins down-regulated in Δ*hfq* mutant compared to WT. Dots in blue, red, green, and orange refer to MTs. The blue, green and red dots were among the 15 MTs (MT5, 9, 10, and 11 were not detected) specific for *X. doucetiae* and chosen for protein expression in Δ*hfq* $P_{BAD}$_*xrdA*. In addition, the blue and red dots, referring to MT3, MT13, MT14, and MT15, were significantly down-regulated in Δ*hfq* and therefore selected for gene deletion in WT. Raw data from the proteomic analysis is shown in **Extended Excel Sheet_2**.

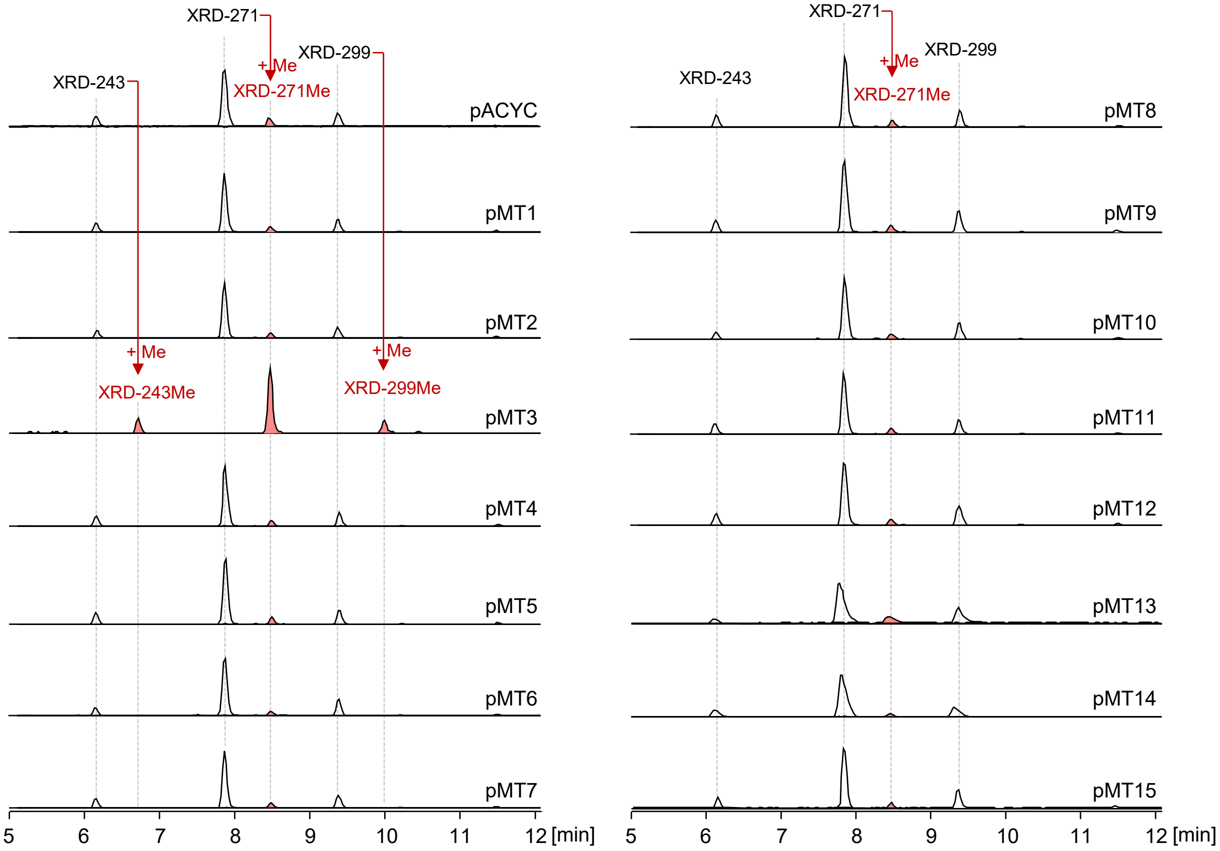


**Figure S5.** EICs of XRDs from the MT-expression strains. MT candidates MT1–MT15 were cloned into vector pACYC and transformed into Δ*hfq* $P_{BAD}$_xrdA, respectively. The negative control Δ*hfq* $P_{BAD}$_xrdA which was transformed with the empty plasmid pACYC can produce XRD-243, XRD-271, XRD-299, and a small amount of *N*-methylated XRD-271Me. In comparison to the control strain, Δ*hfq* $P_{BAD}$_xrdA pMT3 was the only strain producing *N*-methylated XRDs.

| Name | Query coverage | % Pairwise Identity | E Value | Description |
|---|---|---|---|---|
| ctg1_5739 | 86.59% | 22.7% | 2.13e-10 | CP072788.1 Saccharothrix algeriensis strain NRRL B-24137 chromosome |
| ctg1_428 | 69.11% | 24.1% | 4.55e-04 | CP072788.1 Saccharothrix algeriensis strain NRRL B-24137 chromosome |
| ctg1_5135 | 57.32% | 32.6% | 3.64e-14 | CP072788.1 Saccharothrix algeriensis strain NRRL B-24137 chromosome |
| ctg1_6018 | 54.88% | 29.3% | 4.92e-10 | CP072788.1 Saccharothrix algeriensis strain NRRL B-24137 chromosome |
| ctg1_1483 | 54.47% | 30.1% | 9.98e-08 | CP072788.1 Saccharothrix algeriensis strain NRRL B-24137 chromosome |
| ctg1_5263 | 49.19% | 26.6% | 1.82e-04 | CP072788.1 Saccharothrix algeriensis strain NRRL B-24137 chromosome |
| ctg1_4589 | 43.50% | 28.7% | 1.55e-06 | CP072788.1 Saccharothrix algeriensis strain NRRL B-24137 chromosome |
| ctg1_5259 | 41.46% | 30.1% | 4.31e-05 | CP072788.1 Saccharothrix algeriensis strain NRRL B-24137 chromosome |
| ctg1_4087 | 28.46% | 28.4% | 4.35e-02 | CP072788.1 Saccharothrix algeriensis strain NRRL B-24137 chromosome |

**Figure S6.** BLASTP search of XrdM against the genome of *S. algeriensis* NRRL B-24137 results in 9 hits, in which the top 1 hit is CTG1_5739, sharing a sequence coverage of 86.59% and sequence identity of 22.7% to XrdM.

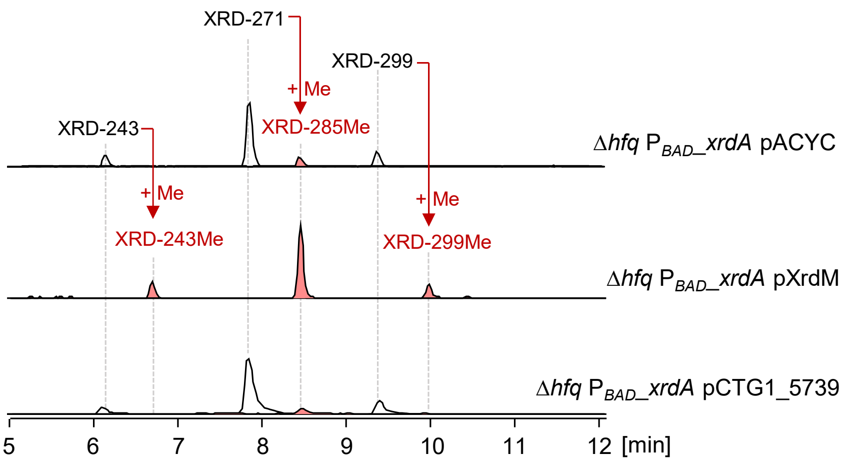


**Figure S7.** EICs of XRDs from Δ*hfq* $P_{BAD}$_xrdA pXrdM and Δ*hfq* $P_{BAD}$_xrdA pCTG1_5739. In comparison to XrdM, CTG1_5739 does not have the methylation activity towards non-methylated XRDs.

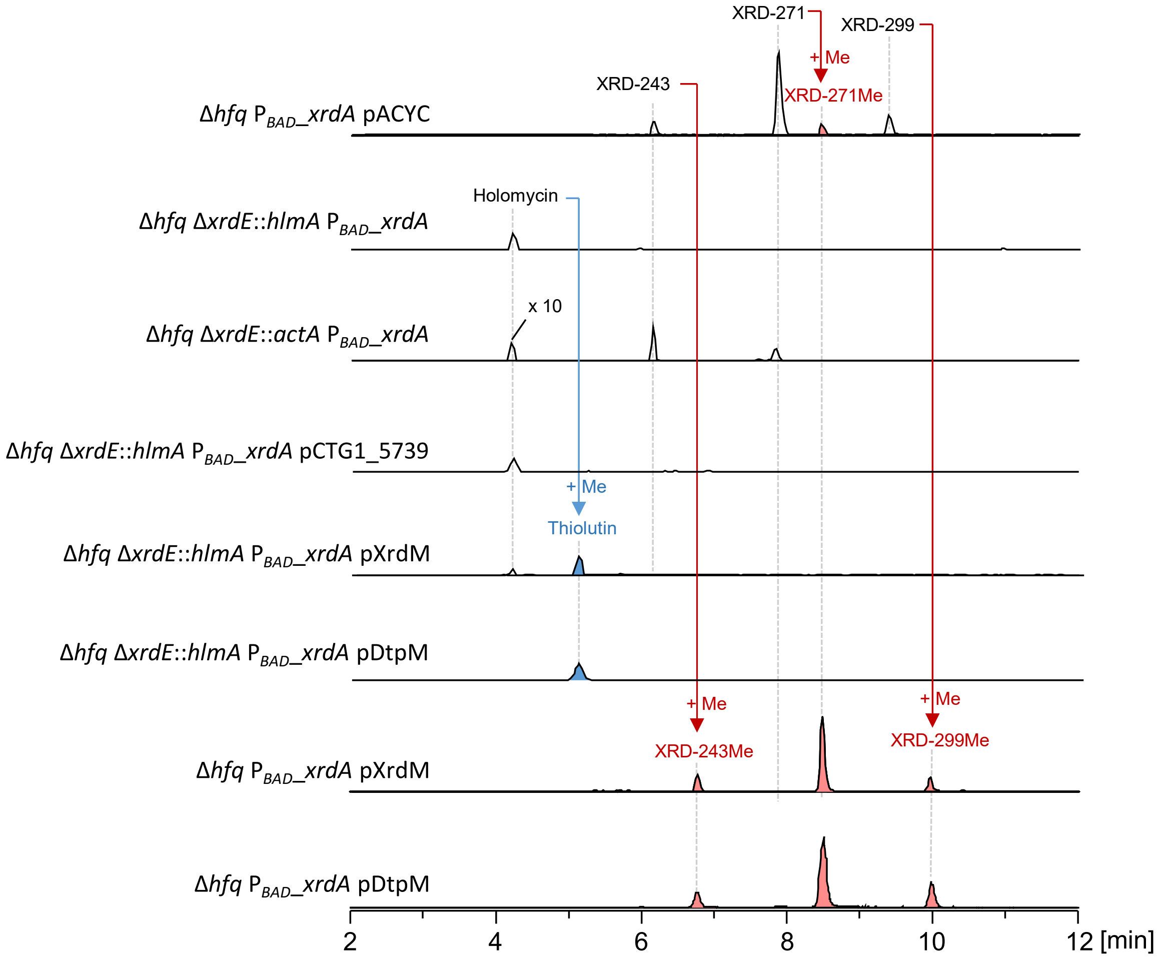


**Figure S8.** EICs of XRDs, holomycin, and thiolutin from swapped strains. Holomycin can be successfully produced by both *hlmA* and *actA* swapped strains. Compared to DtpM[3], CTG1_5739 was not able to methylate holomycin, while XrdM was able to convert holomycin to thiolutin similar as DtpM, even though the activity is lower. The sign of "x 10" indicates that the respective peaks were increased by 10 times in order to fit them to the chromatograms.

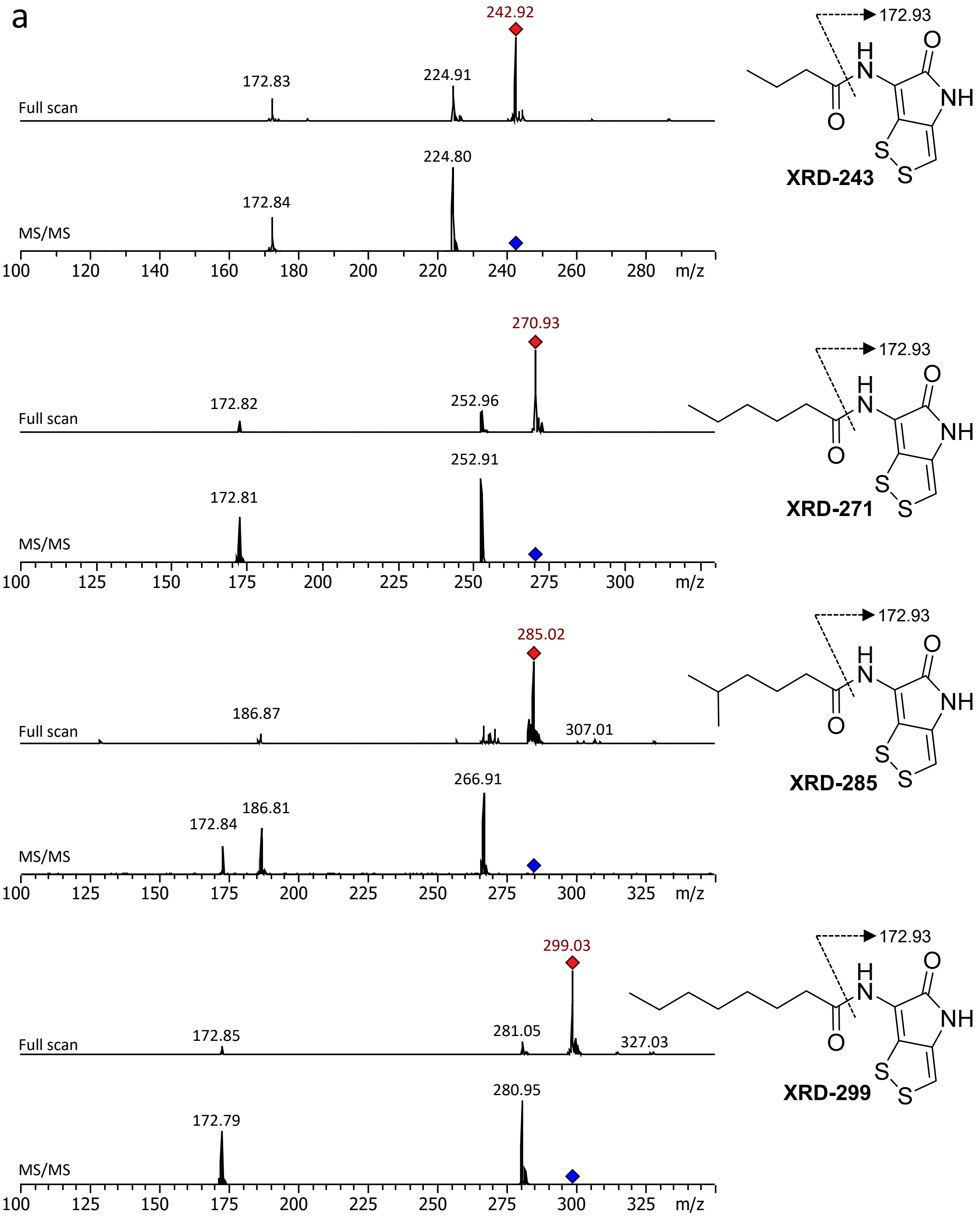
a
242.92
172.93
172.83
224.91
Full scan
224.80
172.84
MS/MS
100 120 140 160 180 200 220 240 260 280 m/z
XRD-243
270.93
172.82
252.96
Full scan
252.91
172.81
MS/MS
100 125 150 175 200 225 250 275 300 m/z
XRD-271
285.02
186.87
307.01
Full scan
266.91
186.81
172.84
MS/MS
100 125 150 175 200 225 250 275 300 325 m/z
XRD-285
299.03
172.85
281.05
327.03
Full scan
280.95
172.79
MS/MS
100 125 150 175 200 225 250 275 300 325 m/z
XRD-299

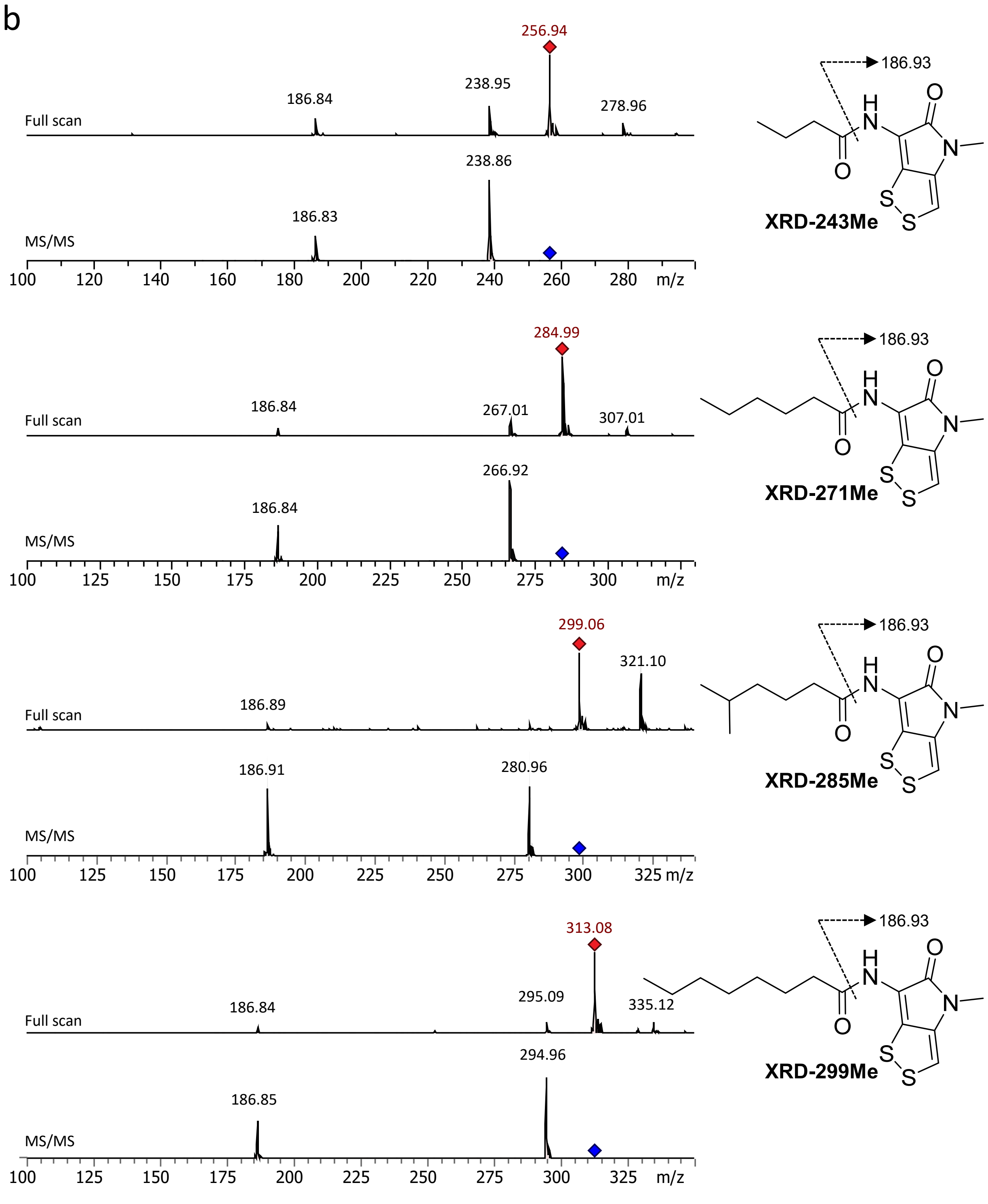
b
256.94
238.95
186.84
278.96
Full scan
238.86
186.83
MS/MS
100 120 140 160 180 200 220 240 260 280 m/z
186.93
XRD-243Me
284.99
186.84
267.01
307.01
Full scan
266.92
186.84
MS/MS
100 125 150 175 200 225 250 275 300 m/z
186.93
XRD-271Me
299.06
321.10
186.89
Full scan
186.91
280.96
MS/MS
100 125 150 175 200 225 250 275 300 325 m/z
186.93
XRD-285Me
313.08
186.84
295.09
335.12
Full scan
294.96
186.85
MS/MS
100 125 150 175 200 225 250 275 300 325 m/z
186.93
XRD-299Me

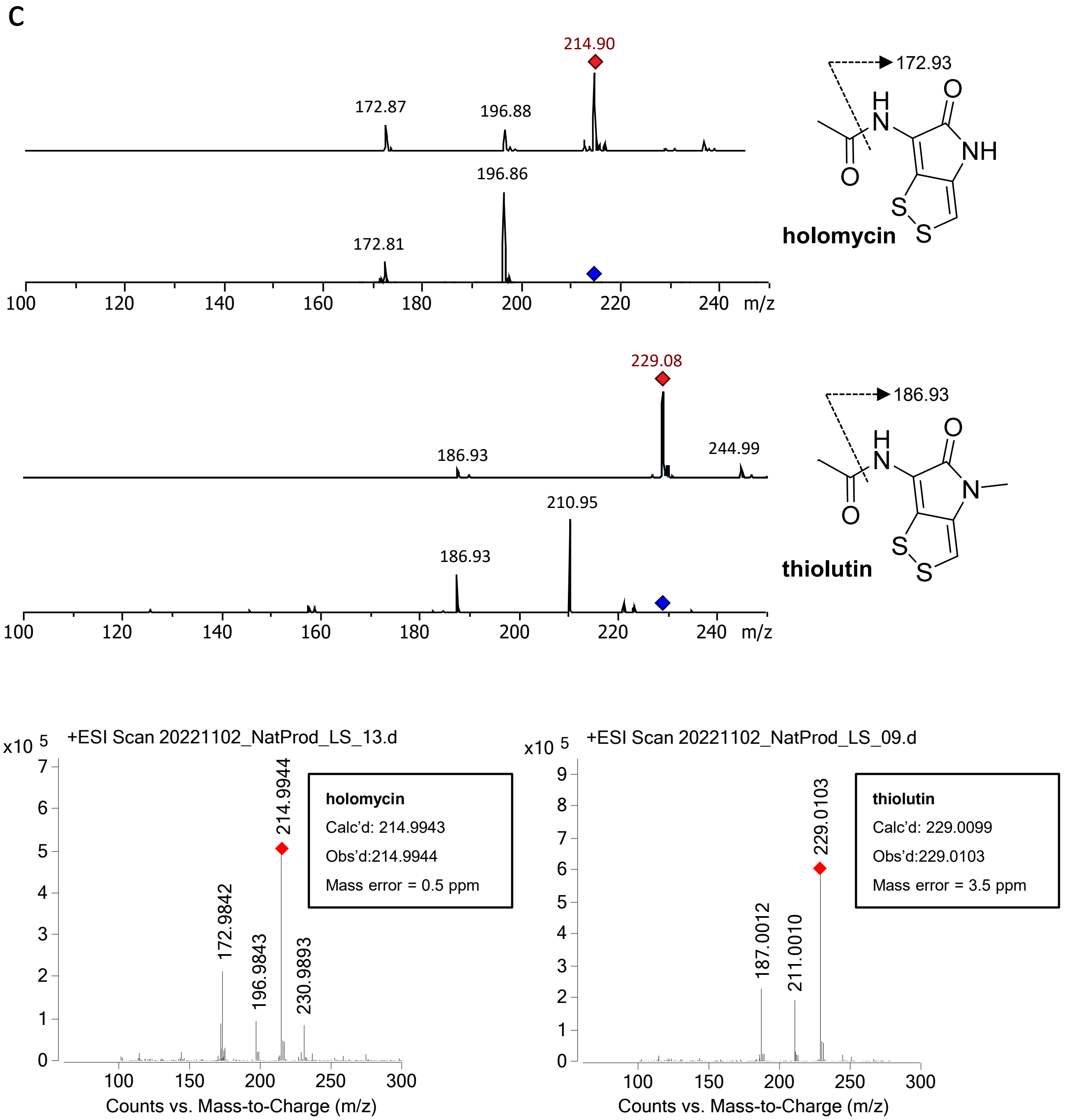


**Figure S9.** MS spectrum and MS/MS fragmentation pattern of non-methylated XRDs (**a**), *N*-methylated XRDs (**b**), holomycin, and thiolutin (**c**). As shown in **a**, the MS/MS spectrum of XRD-285 (frag. 172.84) overlaps with that of XRD-271Me (frag. 186.81). This is due to the fact that XRD-285 observed in WT eluted at almost the same time as XRD-271Me (**Figure S3**).

```
        1        10        20        30        40        50        60        70
DtpM    MAETDDE-RAGARSLLMQRLFGSRVTEVLAAMARLDLADAIGDGI-ADVHDLARSCDL-PADGLHRLLRALAGLGMCEESE
XrdM    MQEEIHYDSIGEGYEHFYNSVPQRKVEVR------TLFDMVGDVQGKSVLDLACGYGYFGRELYHRGASKVVGVDISEKM-
        * *  .    *     : . . .* .**         * * :**    .* *** . .     :  **     :.*:.:.*:

         80        90        100      110       120       130       140       150
DtpM    PGKFALTASGALLRKDHPESVYDFARFHTAPETTRPWTNLEQALRTGRPTFDEHFGSPLYE------YMAGHPELSARFAA
XrdM    ---------IALAKKKS-TEYGDNIEFHVANVSDM---QLNEKFDIITATFLFHYAKSIVELESMFRSVANHLKPSGKLVA
                 ** :*.    .  *   .**.*   :      :*:: :     .**   *:...: *        :*.* : *.::.*

            160       170       180       190       200       210       220       230
DtpM    AMRGESLATADTIAEHYDFSPYRTVTDVGGGDGTLITAILRRHPDLRGTIFETPEIAERAAERVRAAGLHDRCAVVSGDFF
XrdM    YMAAPDYQLEKGNCHNYGL------------------NILSEEPLQGGFIHQVEFIT------------------------
         *  . .     .   ..:*.:                    ** ..*    * *.:.   *:

            240       250       260       270       280       290       300       310
DtpM    DLVPGGADLYLVKSTLHNWDDEHVVRILSSCRTALADRGRLLVIDVVLPDRAEPDPAEL-NPYVKDLQMLVLLGGRERTRA
XrdM    --------TPPILLTFYRWDRETYKNAIHKAGFGHFEWRKP----MVLESDIERYPAGFWDTY------------------
                  :   *::.** *     . : ..   .   :    :    :** .   *   ** : :.*

            320       330       340       350
DtpM    HLDRLCARAGLVIDRVLPLPPHVGLSLTEVVPAPAGPTP
XrdM    --QQNCMHTGLTCW----MP
          :: * ::**.       *
```

**Figure S10.** Sequence alignment of DtpM and XrdM. Their pairwise identity is only 14.6%. Residues are numbered according to the DtpM sequence. Residues His251, Asp279, and Glu310 were supposed to be important for the catalytic function of DtpM.[3] Based on the here reported structural data, we exclude a functional role of Asp279 (gray shaded) for DtpM activity, while His251 and Glu310 (orange shaded) are likely catalytically relevant. In addition, Asn252 (red shaded) was identified to be crucial for DtpM activity. Amino acids shaping the substrate binding pocket of DtpM are shaded in green (from the same monomer) or in blue (from the neighboring monomer).

```
             1        10        20        30        40        50        60        70
DtpM         MAETDDERAGARSLLMQRLFGSRVTEVLAAMARLDLADAIGDGIADVHDLARSCDLPADGLHRLLRALAGLGMCE
WelM    MLSQEKPSTVPSIEASQTMLQMIIINPLVTRAIYAAAKLGIADLLKYGIKSYKELADAVSVKPLFLYRLLRALASVGVFA
MT15                        MGYTYQAALRAATVLGVADHLIDGSKTVYELAQAVGAQEQQLHRVLRLLATRNIFH
                            .     .: * : *.:** :   *     :** : .      *:*:** **   .:

         80        90        100      110       120       130       140       150
DtpM    ESEPGKFALTASGALLRKDHPESVYDFARFHTAPETTRPWTNLEQALRTGRPTFDEHFGSPLYEYMAGHPELSARFAAAM
WelM    EEQEGYFTLTPLANCLVSDIPGSLRALTIINNESELYQVRGNILYSLQNGCNAFEHLYGMPLFEYYIQNPELGKTFDEAM
MT15    EIEEKRFALTPAAEFLRTDVNNSLRAAVLMLTDKTFWQPLGEVVESVR-GNPAFKHIYGLPFFDYWAQTGMPADDFHVGM
        * :    *:**. .   * .*    *:    . : .      :    ::  ::: *   :*.. :* *:::*        .  *  .*

          160       170       180       190       200       210       220       230
DtpM    RGESLATADTIAEHYDFSPYRTVTDVGGGDGTLITAILRRHPDLRGTIFETPEIAERAAERVRAAGLHDRCAVVSGDFFD
WelM    TSVTAMDIAEIIANYDFSKVNKLVDVGAGQGKLLASILKAYPTCKGVLYELPSVSTGAVDLIKAEGLQDRCEIVGGNFLE
MT15    SSMSKVENLFLVHSYDFPKNATVVDVAGGFGGLLLRVLQDNTTLRGILFDRQHVLDRH--ELGALGDDTRWELASGDFFE
         . :      :    ***.    .:.**..* * *:  :*:   .  :* :::    :          : * * . *   :..*:*::

          240       250       260       270       280       290       300       310
DtpM    LVPGGADLYLVKSTLHNWDDEHVVRILSSCRTALADRGRLLVIDVVLPDRAEPDPAELNPYVKDL-QMLVLLGGRERTRA
WelM    SVPAGGDVYMIKSVIHNWDDENAITILKNCHRGMKENGKLLVVEAVIQPGNKPCAGK----FHDL-AMMLVLNGRERTEK
MT15    SCPQG-DIYLLKYIMHDWPDEQCISILRNCRKAMAPNGRVLVMDPVIPDGNISHAGK----EMDLLCMGIYEGGRERTRG
          * * *:*::*  :*:* **: : ** .*: .:  .*::**:: *:      . ..:       **  * :  .*****.

           320       330       340       350
DtpM    HLDRLCARAGLVIDRVLPLPPHVGLSLTEVVPAPAGPTP
WelM    EYQALFEASGFQLTRIIPTMSLM--SVIEGVRIG-----
MT15    ELQQLLTRAGLKLNRVIDTGCYV--SIVEAIAE------
        . : *   :*: : *::      :  *: * :
```

**Figure S11.** Sequence alignment of DtpM, WelM and MT15. MT15 has 34.9% sequence identity to WelM and 31.8% sequence identity to DtpM. Color coding and sequence numbering of residues is according to **Figure S10**.

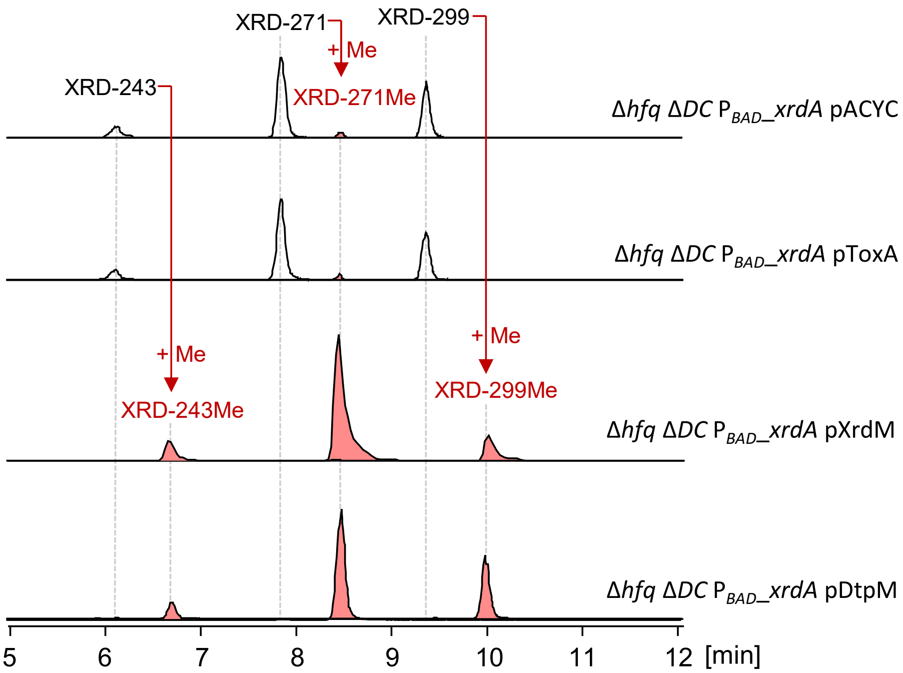


**Figure S12.** EICs of XRDs from Δ*hfq* Δ*DC* $P_{BAD}$_xrdA pACYC and Δ*hfq* Δ*DC* $P_{BAD}$_xrdA pToxA, Δ*hfq* Δ*DC* $P_{BAD}$_xrdA pXrdM, and Δ*hfq* Δ*DC* $P_{BAD}$_xrdA pDtpM. Both XrdM and DtpM can methylate XRDs, but ToxA can not.

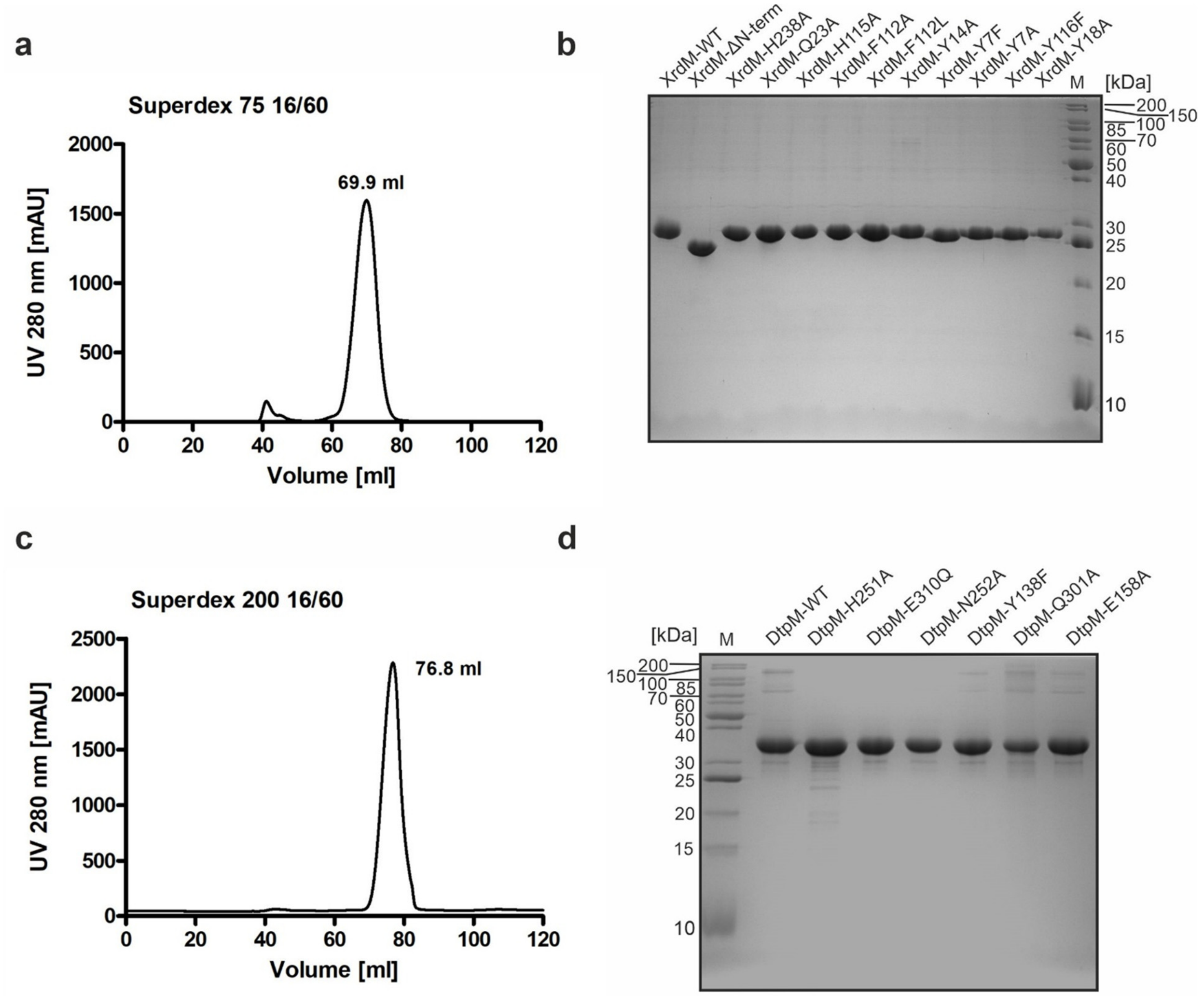


**Figure S13.** Heterologous production and purification of XrdM and DtpM. (**a**) Gel filtration chromatogram for XrdM. In solution, the protein is a monomer. (**b**)SDS-PAGE of purified XrdM and its mutants. (**c**) Size exclusion chromatography chromatogram for DtpM. (**d**) SDS-PAGE of purified DtpM and its mutants.

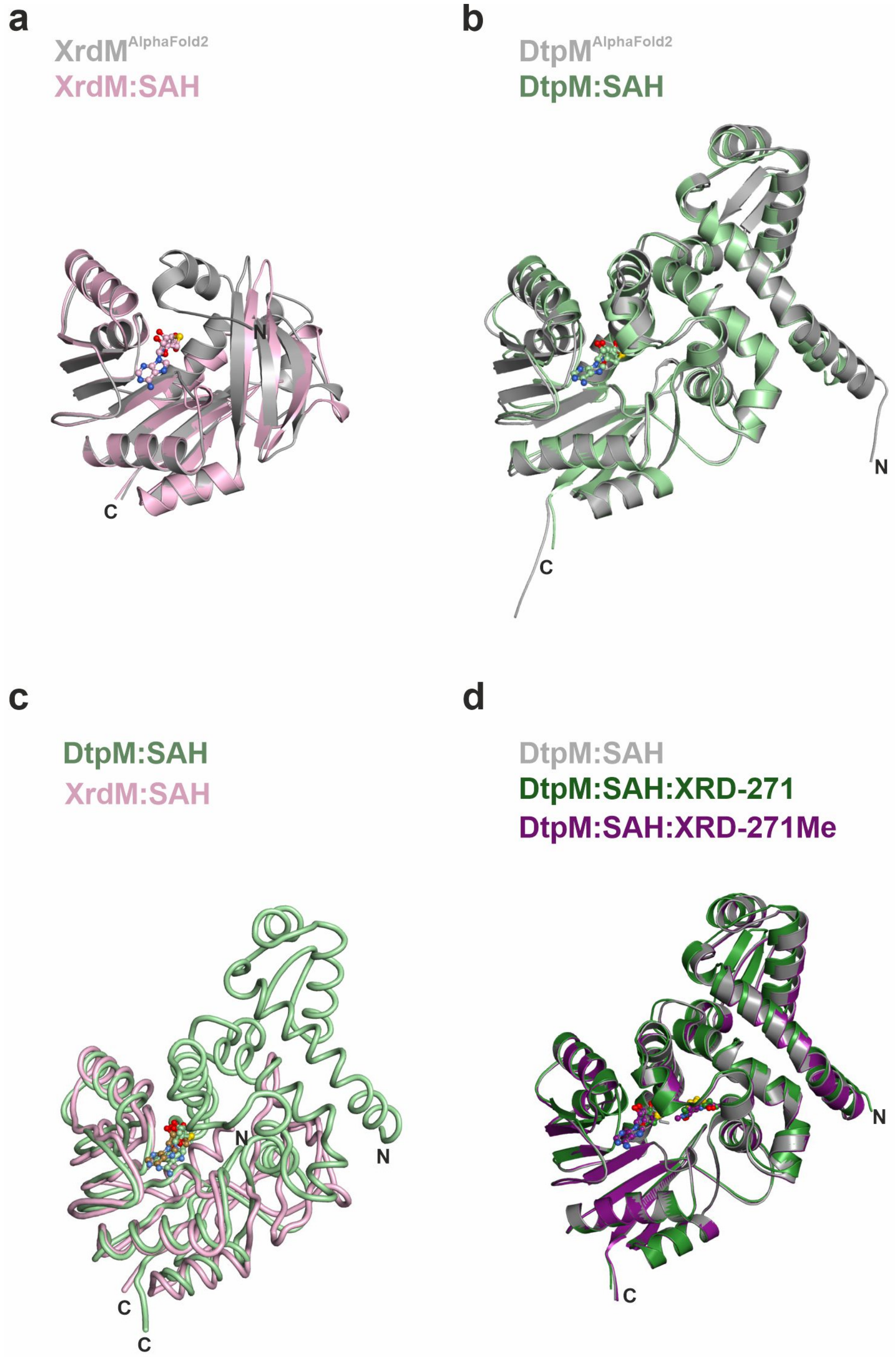


**Figure S14.** Comparison among experimental apo structures, ligand-bound structures and AlphaFold2 predictions. (**a**) Superposition of the X-ray structure of XrdM (lightpink) and the AlphaFold2 prediction of XrdM (gray). The prediction is highly accurate with a route mean square deviation (r.m.s.d.) of 0.39 Å over 147 atoms (align command within PyMOL). The N-terminus of XrdM (residues 4-23) is disordered in the crystal structure. (**b**) Superposition of the X-ray structure of a DtpM monomer (lightgreen) and its AlphaFold2 prediction (gray). The prediction deviates from the X-ray structure more significantly (r.m.s.d of 1.08 Å over 328 atoms, align command within PyMOL) and in consequence phasing of DtpM crystals required truncation of the AlphaFold2 model. For details see Methods section. (**c**) Superposition of a DtpM monomer onto XrdM. The structural alignment has an r.m.s.d. value of 1.67 Å over 518 atoms. (**d**) Superposition of DtpM monomers in the apo and ligand-bound states. No structural changes occur upon ligand binding to the DtpM active site. For better illustration only monomers of DtpM structures are shown.

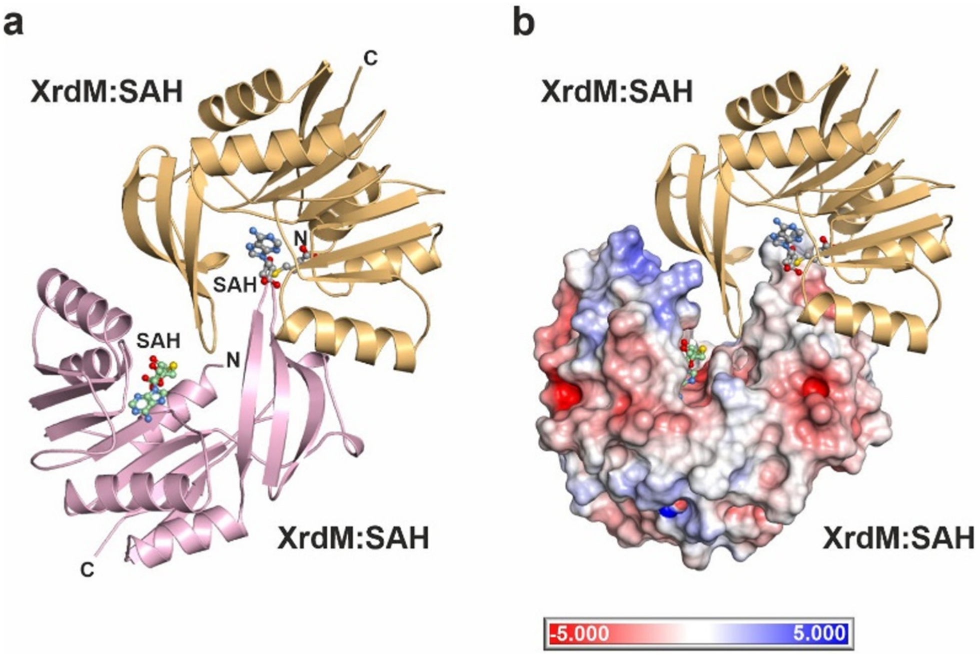


**Figure S15.** Crystal packing of XrdM crystals. (**a**) In the crystal lattice two XrdM molecules tightly interact with each other (both shown as ribbon and colored pink and brownish, respectively). Residues 179-185 of one monomer insert into the substrate binding cleft of the second XrdM monomer, hence preventing substrates from binding. (**b**) Same illustration as shown in panel **a** but with one XrdM molecule being shown as a surface illustration. The crystallographic symmetry mates mutually block their active sites.

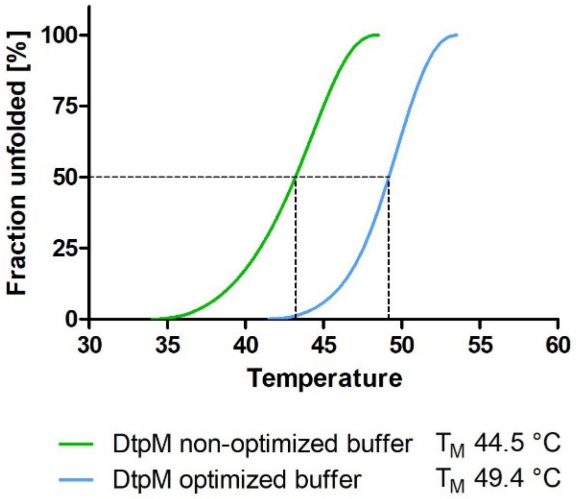


**Figure S16.** Differential scanning fluorimetry of purified DtpM. Different buffer conditions were tested for their impact on the thermal stability of DtpM. A solution containing 100 mM HEPES pH 7.0, 100 mM NaCl and 10% (v/v) glycerol was found to increase the melting temperature ($T_M$) of DtpM by 5 °C compared to a non-optimized buffer of 100 mM Tris/HCl pH 7.5, 100 mM NaCl.

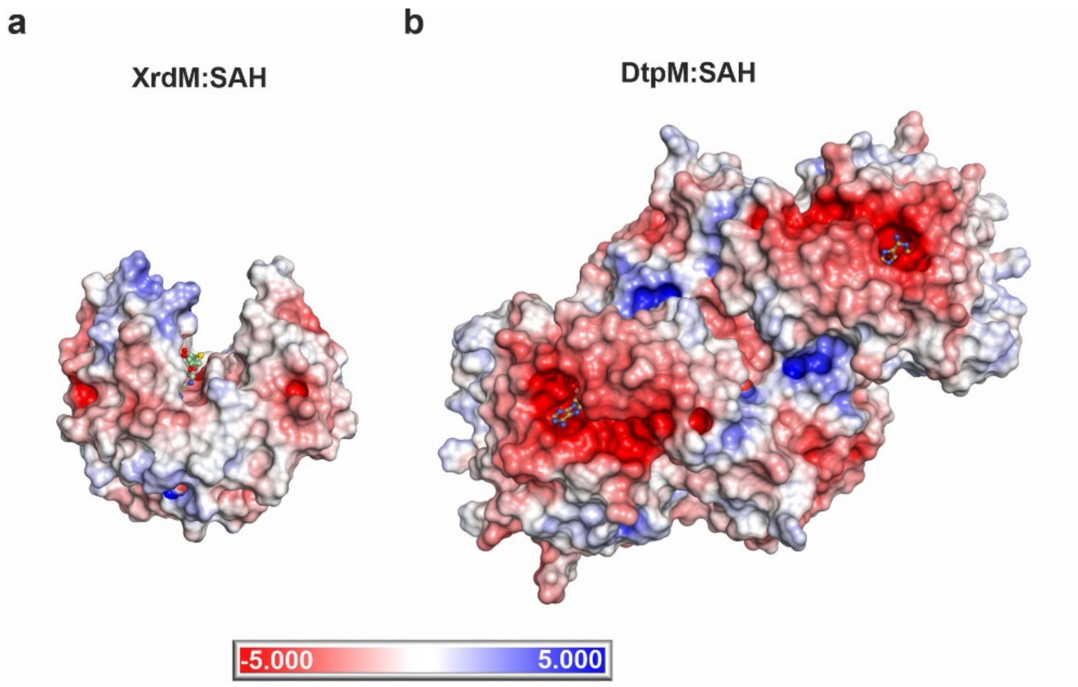


**Figure S17.** Connolly surface representations of monomeric XrdM (**a**) and dimeric DtpM (**b**). Colors indicate positive (blue) and negative (red) electrostatic surface potential, ranging from -5kT/e to + 5kT/e. A deep and solvent accessible substrate binding cleft is visible for XrdM, while the active site of DtpM is more shielded from the environment.

**a**

```
        1           10                    20                  30                 40                  50                  60                  70
DtpM    MAETDDERAGA---RSLLMQRLFGSRVTEVLAAMARLDLADAIGDGIADVHDLARSCDLPADGLHRLLRALAGLGMCEES
LaPhzM  MTEN--NRAGAVPLSSILLQMITGYWVTQSLYVAAKLGIADLVADAPKPIEELAAKTGAKAPLLKRVLRTIASIGVFTET
        *:*.  :****    *:*:* : *  **: * . *:*.:** :.*.    :.:** . .  *  *:*:**::*.:*:  *:

          80        90        100       110       120       130       140       150
DtpM    EPGKFALTASGALLRKDHPESVYDFARFHTAPETTRPWTNLEQALRTGRPTFDEHFGSPLYEYMAGHPELSARFAAAMRG
LaPhzM  EPGIFGITPLAALLRSGTPDSMRPQAIMHG-EEQYRAWADVLHNVQTGETAFEKEFGTSYFGYLAKHPEADRVFNEAQAG
        *** *.:*. .****.. *:*:    * :*    *  *.*::: : ::**..:*::.**:. : *:* *** .  *  *  *

          160       170       180       190       200       210       220       230
DtpM    ESLATADTIAEHYDFSPYRTVTDVGGGDGTLITAILRRHPDLRGTIFETPEIAERAAERVRAAGLHDRCAVVSGDFFDLV
LaPhzM  YTKQVAHAVVDAYDFSPFKTVIDIGAGYGPLLSAILRSQPEARGILFDQPHVAQAAGKRLAEAGVGDRCGTVGGDFFVEV
         :  .*.::.: *****::** *:*.* *.*::**** :*: ** :*: *.:*: *.:*:  **: ***..*.****  *

          240       250       260       270       280       290       300       310
DtpM    PGGADLYLVKSTLHNWDDEHVVRILSSCRTALADRGRLLVIDVVLPDRAEPDPAELNPYVKDLQMLVLLGGRERTRAHLD
LaPhzM  PADGDVYILSLLLHDWDDQRSIEILRNCRRAMPAHGKLLIVELVLPEGEEPFFGK----WLDLHMLVLLGAQERTADEFK
        *...*:*::.   **:***:: :.** .** *:. :*:**:::***:  **   .:       **:******.:***   .:.

          320       330       340       350
DtpM    RLCARAGLVIDRVLPLPPHVGLSLTEVVPAPAGPTP
LaPhzM  TLFAASGFALERVLPT--ASGLSIVEARPI------
         * * :*:.::****     ***:.*. *
```

**b** **c**

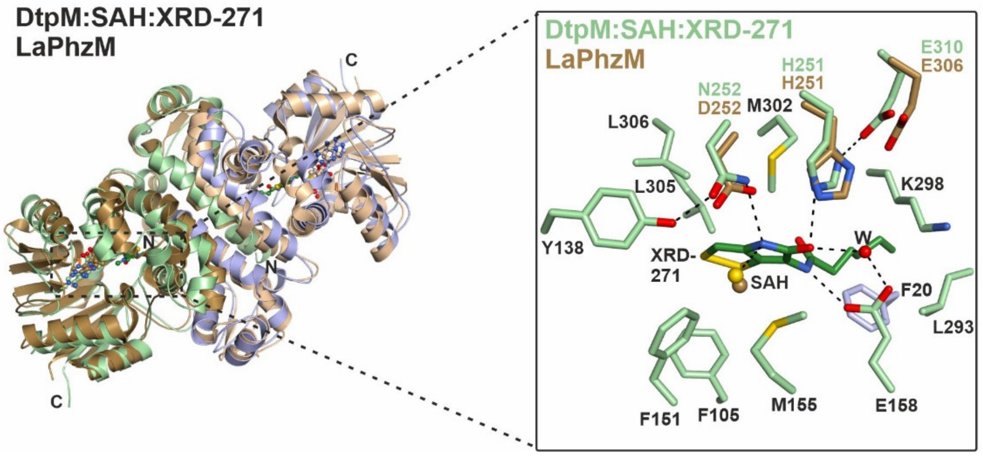


**Figure S18.** Comparison between DtpM and LaPhzM (PDB: 6C5B[4]). (**a**) Sequence alignment of DtpM and LaPhzM. Color coding and sequence numbering according to Figure S10. LaPhzM has the same/functionally equivalent catalytic residues as DptM (His251, N252 and E310 of DtpM correspond to His251, D252 and E306 in LaPhzM). (**b**) Superposition of dimeric DtpM (light green/purple) with dimeric LaPhzM (brownish colors) illustrates their structural similarity . (**c**) Stick presentation of the active site of DtpM and LaPhzM. For clarity only the assumed catalytic residues of LaPhzM (His251, D252 and E306; brown) are shown. They adopt the same position relative to the cofactor SAH as their equivalents in DptM. Hydrogen bonds are indicated as black dotted lines for DtpM according to **Figure 4c**.

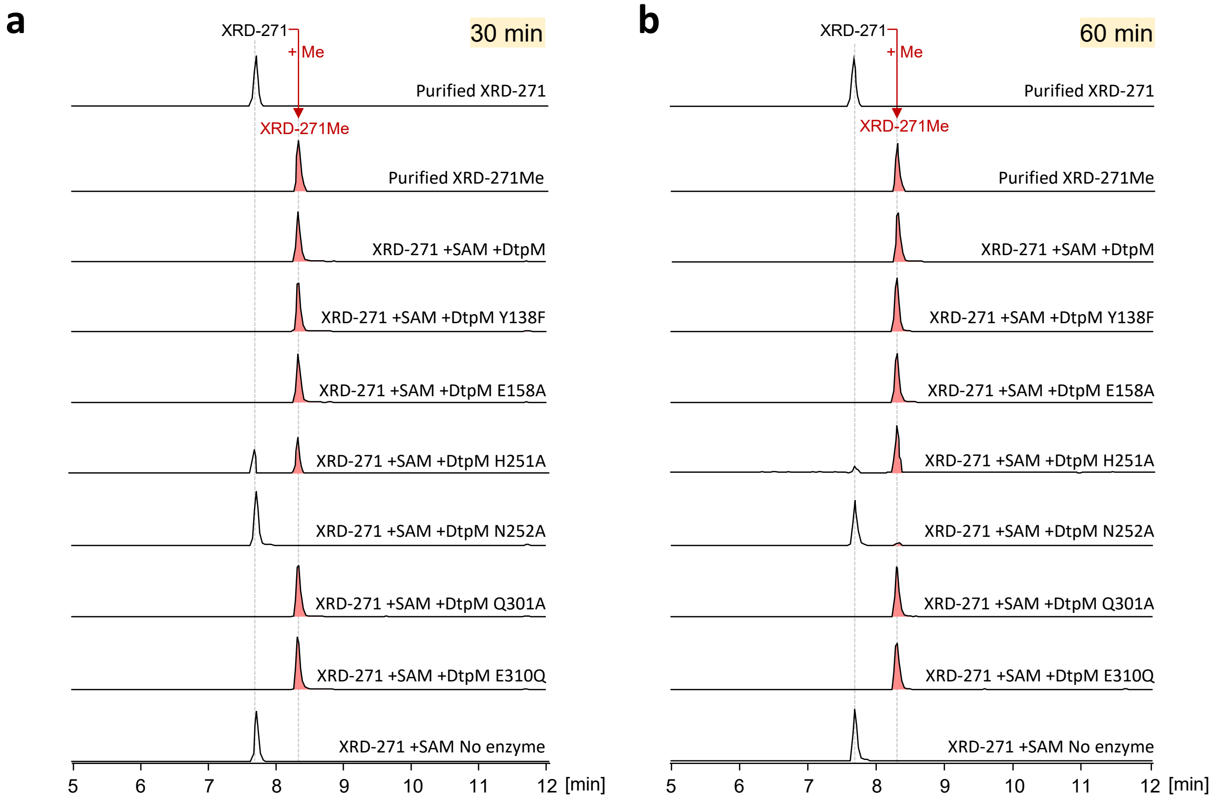


**Figure S19.** Activity assay of wild type and mutant DtpM using XRD-271 as substrate after incubation of 30min (**a**) and 60min (**b**). No enzyme-added reaction and the purified XRD-271 and XRD-271Me were used as control.

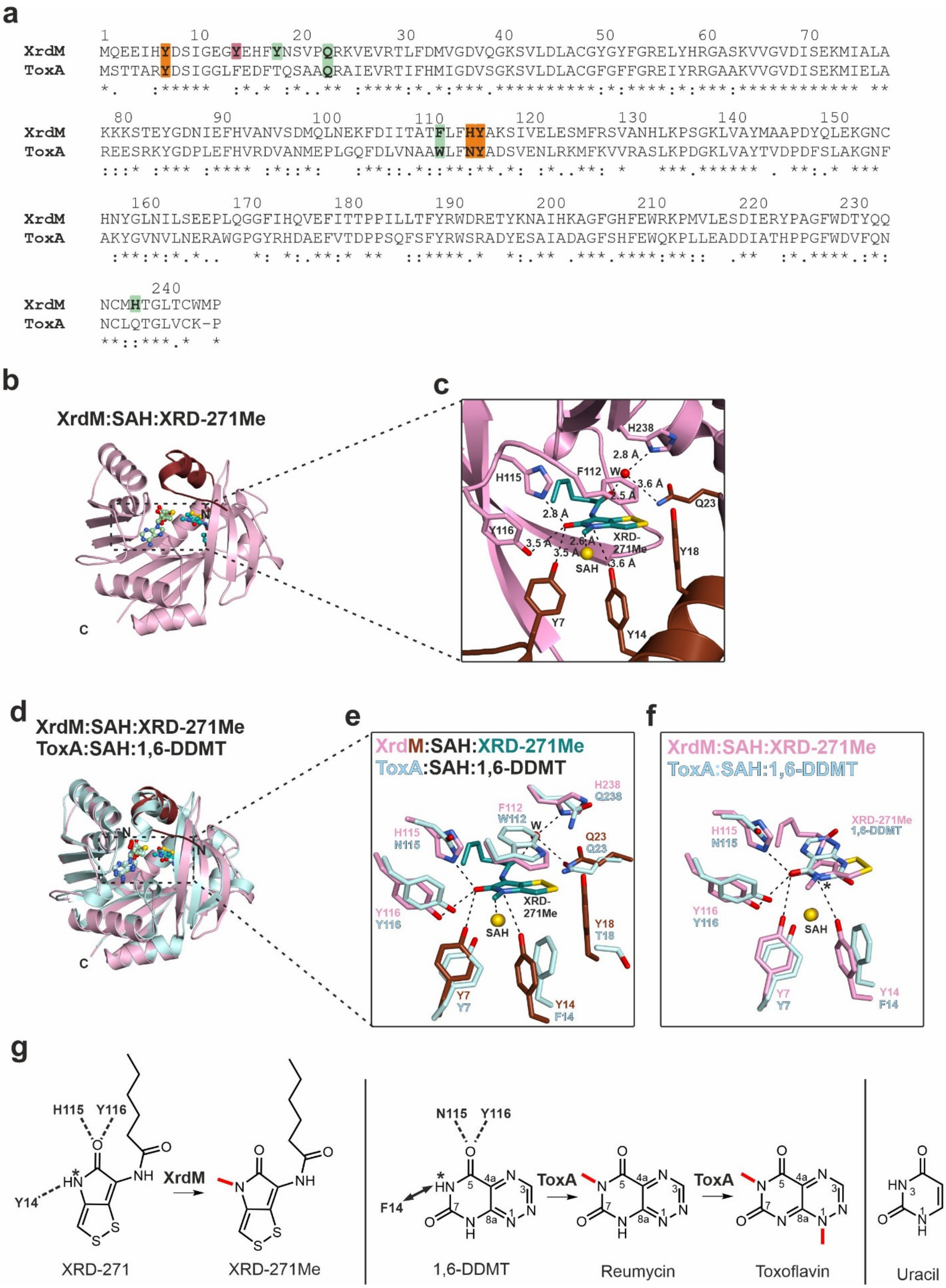
a
XrdM MQEEIHYDSIGEGYEHFYNSVPQRKVEVRTLFDMVGDVQGKSVLDLACGYGYFGRELYHRGASKVVGVDISEKMIALA
ToxA MSTTARYDSIGGLFEDFTQSAAQRAIEVRTIFHMIGDVSGKSVLDLACGFGFFGREIYRRGAAKVVGVDISEKMIELA
XrdM KKKSTEYGDNIEFHVANVSDMQLNEKFDIITATFLFHYAKSIVELESMFRSVANHLKPSGKLVAYMAAPDYQLEKGNC
ToxA REESRKYGDPLEFHVRDVANMEPLGQFDLVNAAWLFNYADSVENLRKMFKVVRASLKPDGKLVAYTVDPDFSLAKGNF
XrdM HNYGLNILSEEPLQGGFIHQVEFITTPPILLTFYRWDRETYKNAIHKAGFGHFEWRKPMVLESDIERYPAGFWDTYQQ
ToxA AKYGVNVLNERAWGPGYRHDAEFVTDPPSQFSFYRWSRADYESAIADAGFSHFEWQKPLLEADDIATHPPGFWDVFQN
XrdM NCMHTGLTCWMP
ToxA NCLQTGLVCK-P
b
XrdM:SAH:XRD-271Me
C
c
H238
2.8 Å
F112
W
3.6 Å
H115
2.5 Å
Q23
2.8 Å
Y116
2.6 Å
XRD-271Me
3.5 Å
3.5 Å
Y18
SAH
3.6 Å
Y7
Y14
d
XrdM:SAH:XRD-271Me
ToxA:SAH:1,6-DDMT
N
N
C
e
XrdM:SAH:XRD-271Me
ToxA:SAH:1,6-DDMT
H238
Q238
F112
W112
W
H115
N115
Q23
Q23
XRD-271Me
SAH
Y116
Y116
Y18
T18
Y7
Y7
Y14
F14
f
XrdM:SAH:XRD-271Me
ToxA:SAH:1,6-DDMT
H115
N115
XRD-271Me
1,6-DDMT
Y116
Y116
SAH
Y7
Y7
Y14
F14
g
H115
Y116
Y14
XrdM
XRD-271
XRD-271Me
N115
Y116
F14
ToxA
ToxA
1,6-DDMT
Reumycin
Toxoflavin
Uracil

**Figure S20.** *In silico* modeling of the XrdM:XRD-271Me complex and comparison of XrdM with ToxA. (**a**) Sequence alignment of XrdM and ToxA with key residues relevant for substrate binding (green) or catalysis (red (essential for XrdM), orange (impactful for XrdM)). Residue numbers are given for XrdM. (**b**) Ribbon illustration of the modelled XrdM:SAH:XRD-271Me complex. The N-terminal segment, that was not resolved in the X-ray structure shown in **Figure 4a**, is highlighted in dark red. (**c**) A zoom-in picture of the XrdM active site with bound XRD-271Me (teal) according to the modelling. Amino acid residues required for ligand binding or catalysis are labeled by the one-letter code and numbered. W indicates a defined water molecule that might contribute to ligand coordination. Predicted hydrogen bonds are indicated by black dotted lines and distances are given. (**d**) Superposition of the XrdM:SAH:XRD-271Me structure with ToxA:SAH:1,6-didemethyltoxoflavin (1,6-DDMT) (PDB: 5JE0[5]) shown as ribbon illustration with the ligands illustrated as ball-and-sticks model. (**e**) Zoom-in at the active site of the superposition shown in panel **d** with only key amino acid side chains shown as sticks. Residues 7, 23, 112, 115 and 116 are conserved in both methyltransferases and adopt similar positions relative to the cofactor SAH. Hydrogen bonds are indicated as black dotted lines for XrdM according to panel **c**. (**f**) Superposition of the active sites of XrdM (pink) and ToxA (light blue) with their respective ligands XRD-271Me and 1,6-DDMT. The site of methylation is indicated by an asterisk. Hydrogen bonds are indicated for the interaction of XrdM with XRD-271Me by black dotted lines. Note that ToxA encodes Phe14 instead of Tyr14. Activation of the carbonyl oxygen atom at position C5 (see also panel **g**) is ensured by Asn115 and Tyr116. (**g**) Chemical reactions catalyzed by ToxA (left) and XrdM (right). The substrates are oriented similarly and the sites of methylation are indicated by an asterisk. Residues that presumably activate the substrate by hydrogen bonds (black dotted lines) are indicated by the one-letter-code and numbered. ToxA does not encode Tyr14 but Phe14. Either methylation of 1,6-DDMT doesn't need activation by Tyr14, as 1,6-DDMT similarly to the structurally related uracil base acts as a week acid and can exist as a mixture of two anions deprotonated at positions N1 or N3 in weakly-alkaline aqueoussolution[6,7] or the aromatic character of residue 14, which is preserved in both ToxA and XrdM, serves exclusively to stabilize the transition state of the $S_N2$ reaction.

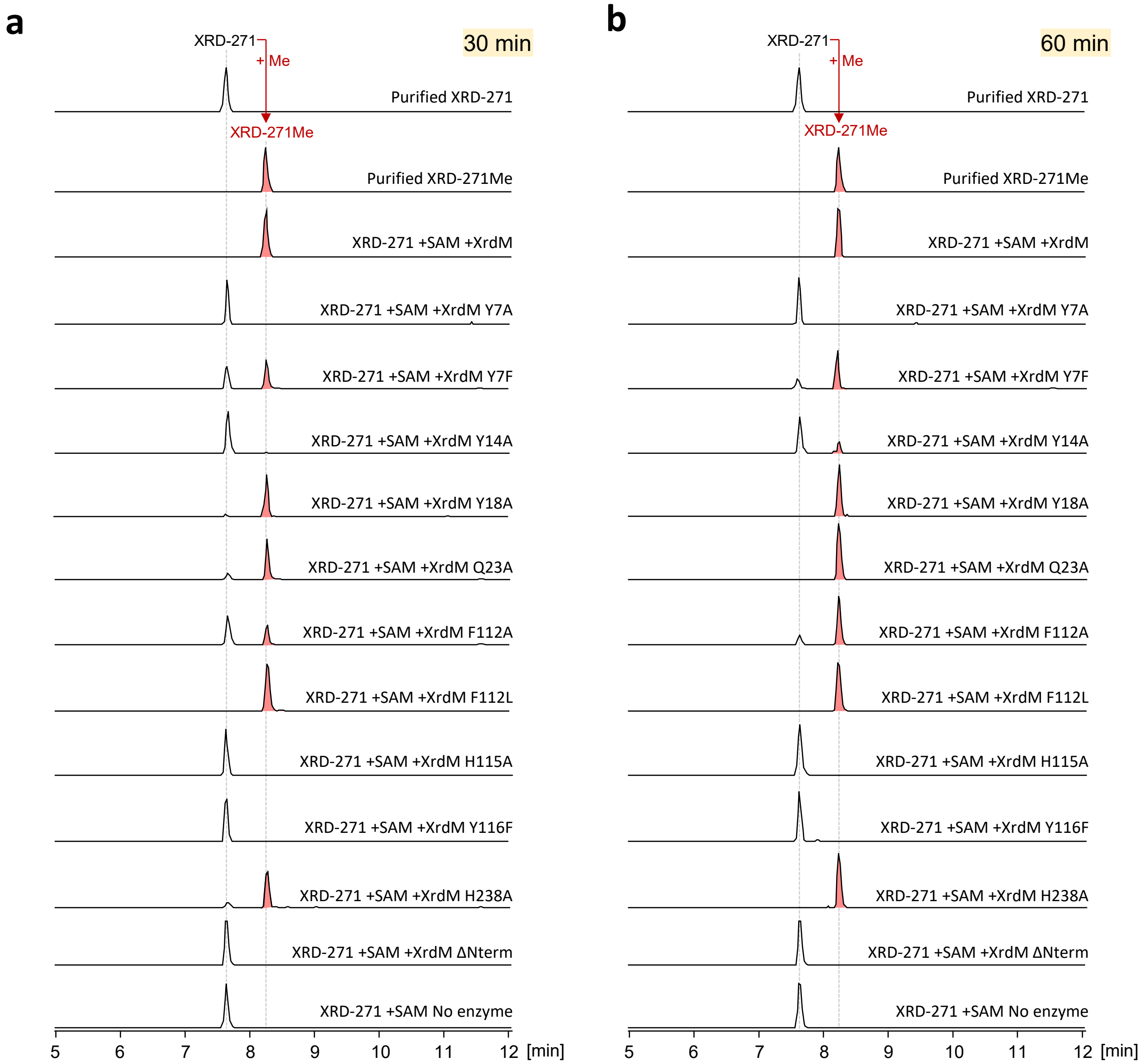


**Figure S21.** Activity assay of wild type and mutant XrdM using XRD-271 as substrate after incubation of 30min (**a**) and 60min (**b**). No enzyme-added reaction and the purified XRD-271 and XRD-271Me were used as control.

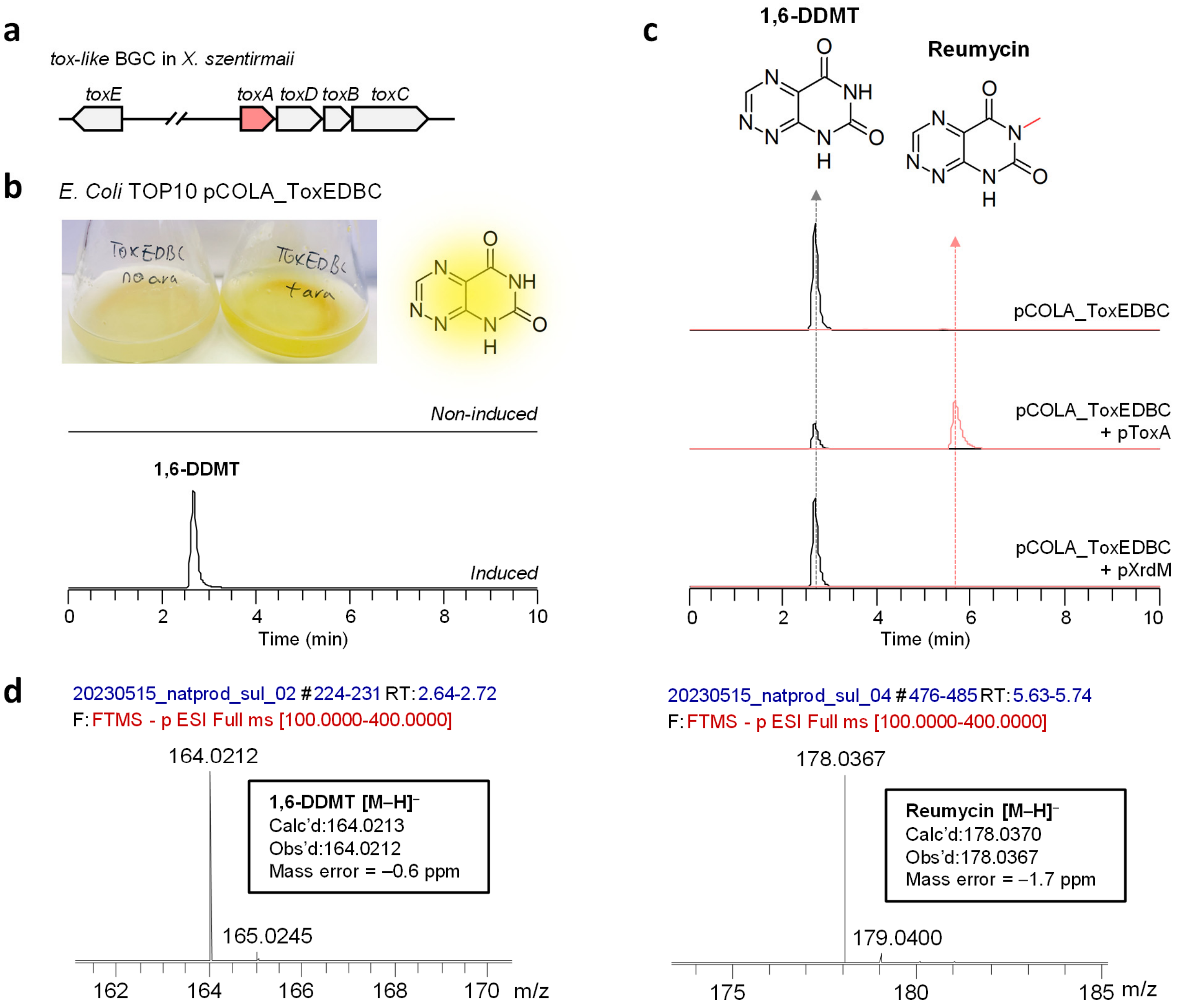


**Figure S22.** Methylation assay of ToxA and XrdM against 1,6-DDMT (write full name here as well). (**a**) The gene organization of a *tox-like* BGC from *X. szentirmaii.* (**b**) Production of the yellow product 1,6-DDMT in *E. coli* TOP10 pCOLA_ToxEDBC after induction with arabinose. EICs from LC-MS analysis of the induced culture confirmed the identity of 1,6-DDMT. (**c**) 1,6-DDMT can be methylated by ToxA to reumycin but not by XrdM. (**d**) LC-MS spectrum under negative mode revealed the mass value of 164.0212 and 178.0367 which is consistent with that of 1,6-DDMT (*m/z* 164.0213, [M-H]$^-$) and reumycin (*m/z* 178.0370, [M-H]$^-$).

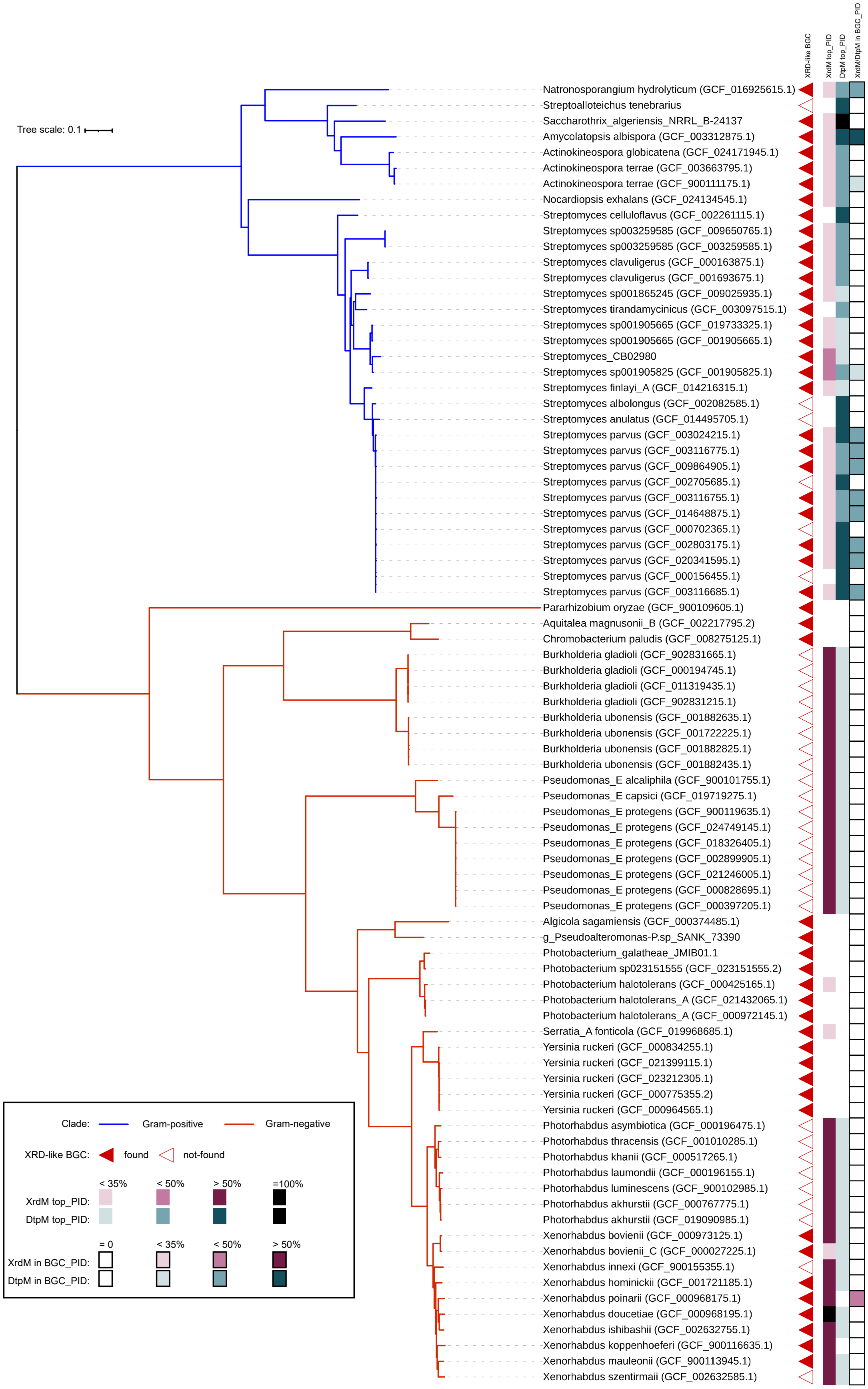
XRD-like BGC
XrdM top_PID
DtpM top_PID
XrdM/DtpM in BGC_PID
Tree scale: 0.1
Natronosporangium hydrolyticum (GCF_016925615.1)
Streptoalloteichus tenebrarius
Saccharothrix_algeriensis_NRRL_B-24137
Amycolatopsis albispora (GCF_003312875.1)
Actinokineospora globicatena (GCF_024171945.1)
Actinokineospora terrae (GCF_003663795.1)
Actinokineospora terrae (GCF_900111175.1)
Nocardiopsis exhalans (GCF_024134545.1)
Streptomyces celluloflavus (GCF_002261115.1)
Streptomyces sp003259585 (GCF_009650765.1)
Streptomyces sp003259585 (GCF_003259585.1)
Streptomyces clavuligerus (GCF_000163875.1)
Streptomyces clavuligerus (GCF_001693675.1)
Streptomyces sp001865245 (GCF_009025935.1)
Streptomyces tirandamycinicus (GCF_003097515.1)
Streptomyces sp001905665 (GCF_019733325.1)
Streptomyces sp001905665 (GCF_001905665.1)
Streptomyces_CB02980
Streptomyces sp001905825 (GCF_001905825.1)
Streptomyces finlayi_A (GCF_014216315.1)
Streptomyces albolongus (GCF_002082585.1)
Streptomyces anulatus (GCF_014495705.1)
Streptomyces parvus (GCF_003024215.1)
Streptomyces parvus (GCF_003116775.1)
Streptomyces parvus (GCF_009864905.1)
Streptomyces parvus (GCF_002705685.1)
Streptomyces parvus (GCF_003116755.1)
Streptomyces parvus (GCF_014648875.1)
Streptomyces parvus (GCF_000702365.1)
Streptomyces parvus (GCF_002803175.1)
Streptomyces parvus (GCF_020341595.1)
Streptomyces parvus (GCF_000156455.1)
Streptomyces parvus (GCF_003116685.1)
Pararhizobium oryzae (GCF_900109605.1)
Aquitalea magnusonii_B (GCF_002217795.2)
Chromobacterium paludis (GCF_008275125.1)
Burkholderia gladioli (GCF_902831665.1)
Burkholderia gladioli (GCF_000194745.1)
Burkholderia gladioli (GCF_011319435.1)
Burkholderia gladioli (GCF_902831215.1)
Burkholderia ubonensis (GCF_001882635.1)
Burkholderia ubonensis (GCF_001722225.1)
Burkholderia ubonensis (GCF_001882825.1)
Burkholderia ubonensis (GCF_001882435.1)
Pseudomonas_E alcaliphila (GCF_900101755.1)
Pseudomonas_E capsici (GCF_019719275.1)
Pseudomonas_E protegens (GCF_900119635.1)
Pseudomonas_E protegens (GCF_024749145.1)
Pseudomonas_E protegens (GCF_018326405.1)
Pseudomonas_E protegens (GCF_002899905.1)
Pseudomonas_E protegens (GCF_021246005.1)
Pseudomonas_E protegens (GCF_000828695.1)
Pseudomonas_E protegens (GCF_000397205.1)
Algicola sagamiensis (GCF_000374485.1)
g_Pseudoalteromonas-P.sp_SANK_73390
Photobacterium_galatheae_JMIB01.1
Photobacterium sp023151555 (GCF_023151555.2)
Photobacterium halotolerans (GCF_000425165.1)
Photobacterium halotolerans_A (GCF_021432065.1)
Photobacterium halotolerans_A (GCF_000972145.1)
Serratia_A fonticola (GCF_019968685.1)
Yersinia ruckeri (GCF_000834255.1)
Yersinia ruckeri (GCF_021399115.1)
Yersinia ruckeri (GCF_023212305.1)
Yersinia ruckeri (GCF_000775355.2)
Yersinia ruckeri (GCF_000964565.1)
Photorhabdus asymbiotica (GCF_000196475.1)
Photorhabdus thracensis (GCF_001010285.1)
Photorhabdus khanii (GCF_000517265.1)
Photorhabdus laumondii (GCF_000196155.1)
Photorhabdus luminescens (GCF_900102985.1)
Photorhabdus akhurstii (GCF_000767775.1)
Photorhabdus akhurstii (GCF_019090985.1)
Xenorhabdus bovienii (GCF_000973125.1)
Xenorhabdus bovienii_C (GCF_000027225.1)
Xenorhabdus innexi (GCF_900155355.1)
Xenorhabdus hominickii (GCF_001721185.1)
Xenorhabdus poinarii (GCF_000968175.1)
Xenorhabdus doucetiae (GCF_000968195.1)
Xenorhabdus ishibashii (GCF_002632755.1)
Xenorhabdus koppenhoeferi (GCF_900116635.1)
Xenorhabdus mauleonii (GCF_900113945.1)
Xenorhabdus szentirmaii (GCF_002632585.1)
Clade: Gram-positive Gram-negative
XRD-like BGC: found not-found
< 35% < 50% > 50% =100%
XrdM top_PID:
DtpM top_PID:
= 0 < 35% < 50% > 50%
XrdM in BGC_PID:
DtpM in BGC_PID:

**Figure S23.** Phylogenetic classification tree of relevant genomes that showed detected XRD-like BGCs or similar MT to XrdM/DtpM. Blue and red branch colors show phyla for Actinomycetota (Gram-positive) and Pseudomonadota (Gram-negative) respectively. Red triangles indicate a detected XRD-like BGC. Best homolog for XrdM/DtpM found anywhere in the genome are indicated by purple and teal squares respectively. The squares with a black border indicate a homolog found within the BGC borders. The varying shades indicate amino acid percent identity (PID) of the hit where white is not detected, light is below 35%, medium below 50%, dark above 50%, and black at 100%. With the exception of *Amycolatopsis albispora*, the best hits outside of the cluster are not the same as those in the cluster. This survey suggests that most of the highly similar homologs of DtpM are found in Gram-positive bacteria, whereas XrdM, in contrast, are found in Gram-negative bacteria.

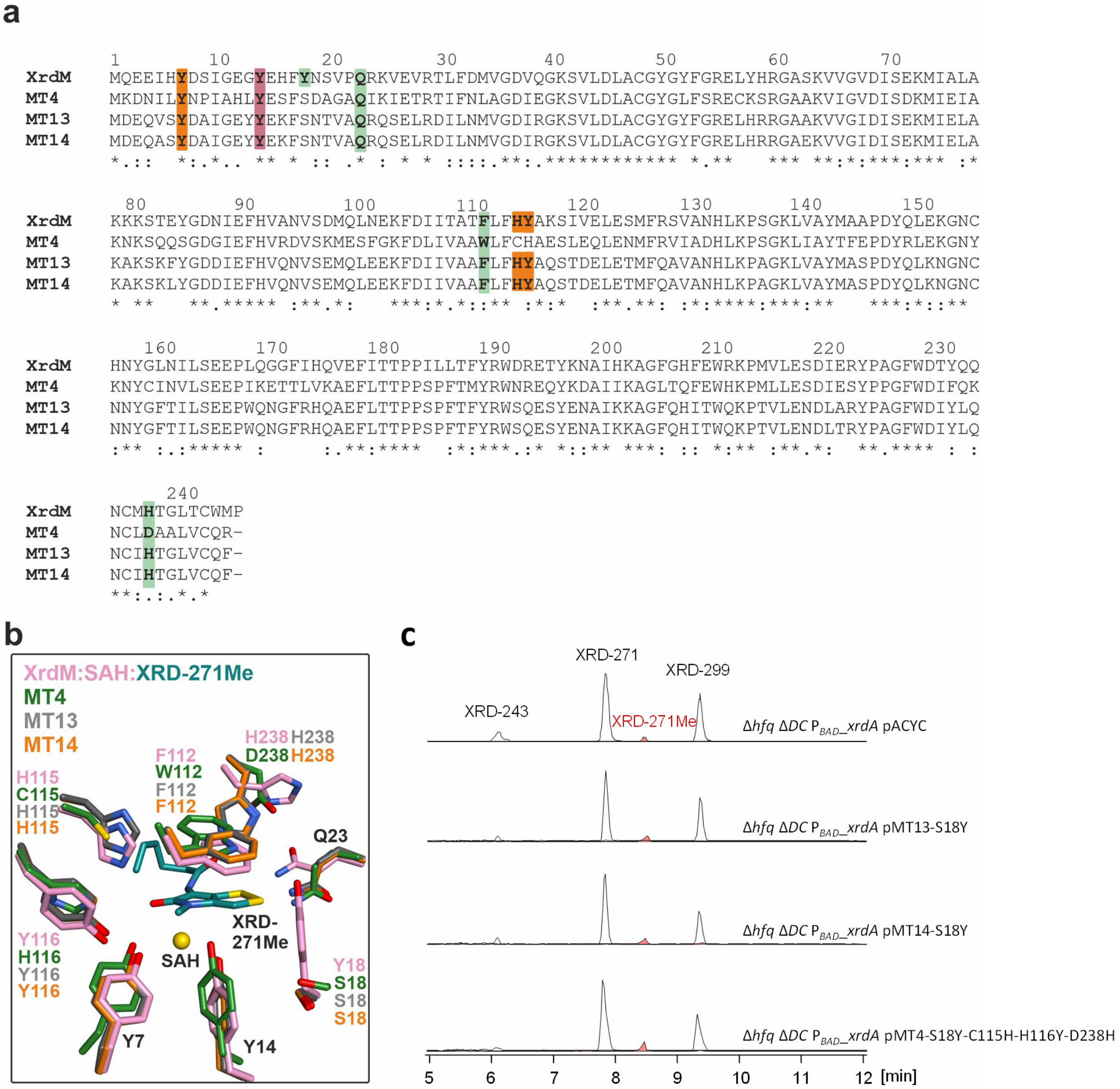


**Figure S24.** (**a**) Sequence alignment among XrdM, MT4, MT13 and MT14. The pairwise identity between XrdM and MT4, MT13 and MT14 is 57.1%, 70.2%, and 70.2%, respectively. Residue numbering and color coding is according to Figure S20. (**b**) Superposition of the XrdM:SAH:XRD-271Me model onto the AlphaFold2 predictions of MT4, MT13 and MT14. (**c**) EICs of XRDs from LC-MS analysis of mutant MT4, MT13 and MT14.

**Table S1.** Antimicrobial activity of XRD-271 and XRD-271Me.

| **Test Organism** | **MIC* (μg/mL) of XRD-271** | **MIC* (μg/mL) of XRD-271Me** |
|---|---|---|
| *Saccharomyces cerevisiae* CEN. PK2 | ≥8 | 2 |
| *Micrococcus luteus* DSM 20030 | 0.1 | 0.25 |
| *E. coli* MG1655 *ΔtolC Δbam* | 1 | 2 |

*The MIC was defined as the lowest concentration of compound completely inhibiting visible growth after 16 to 24 h of incubation at 30°C. Raw data from the MIC test was shown in **Extended Excel Sheet_1**.

**Table S2.** SAM-dependent methyltransferases in *X .doucetiae FRM16*.

| Locus tag | Annotation | Homolog in *X. szentirmaii* DSM 16338 | Homolog in *X. nematophila* ATCC 19061 |
|---|---|---|---|
| XDD1_0024 (MT1) | BREX-1 system adenine-specific DNA-Methyltransferase PglX | no | no |
| XDD1_0042 | Ribosomal RNA small subunit methyltransferase G, RsmG | WP_038240664.1 (86%) | WP_013183046.1 (90%) |
| XDD1_0068 | DNA adenine methyltransferase | WP_038237463.1 (91%) | WP_010847520.1 (93%) |
| XDD1_0097 | 16S rRNA (guanine(1516)-N(2))-methyltransferase RsmJ | WP_038237536.1 (89%) | WP_010848399.1 (92%) |
| XDD1_0159 | tRNA (cytidine/uridine-2'-O-)-methyltransferase TrmL | WP_038239915.1 (97%) | WP_010847601.1 (97%) |
| XDD1_0263 | tRNA (uracil-5-)-methyltransferase | WP_038240061.1 (93%) | WP_ 010847886.1(93%) |
| XDD1_0310 | Ribosomal RNA small subunit methyltransferase B, RsmB | WP_038240457.1 (84%) | WP_ 013183232.1 (89%) |
| XDD1_0361 | putative uroporphyrinogen-III C-methyltransferase | WP_071991849.1 (66%) | WP_ 010845306.1 (71%) |
| XDD1_0430 | 23S rRNA (guanosine-2'-O-)-methyltransferase RlmB | WP_038242355.1 (94%) | WP_ 010845248.1 (95%) |
| XDD1_0471 | Ribosomal RNA large subunit methyltransferase E, RlmE | WP_038237669.1 (97%) | WP_ 010845213.1 (99%) |
| XDD1_0699 (MT2) | Cytosine-specific methyltransferase | WP_038239551.1 (25%) | WP_ 010846361.1 (25%) |
| XDD1_0721 | Ribosomal RNA small subunit methyltransferase C, RsmC | WP_038235016.1 (87%) | WP_ 010846325.1 (91%) |
| XDD1_0981 | conserved protein of unknown function | WP_038237794.1 (77%) | WP_ 010846228.1 (79%) |
| XDD1_1102 (MT13) | Class I SAM-dependent Methyltransferase | WP_038240607.1 (82%) | WP_ 010845997.1 (52%) |
| XDD1_1130 | N-6 DNA methylase | WP_038241382.1 (78%) | WP_ 010846348.1 (77%) |
| XDD1_1136 | tRNA (guanine-N(7)-)-methyltransferase, TrmB | WP_038239339.1 (92%) | WP_ 013184854.1 (91%) |
| XDD1_1159 | Ribosomal RNA small subunit methyltransferase E, RsmE | WP_038235389.1 (86%) | WP_ 010846610.1 (88%) |
| XDD1_1209 | tRNA (guanine-N(1)-)-methyltransferase, trmE | WP_038235274.1 (97%) | WP_ 010845357.1 (96%) |
| XDD1_1234 | Uncharacterized tRNA/rRNA methyltransferase YfiF | WP_038238714.1 (86%) | WP_ 010848714.1 (86%) |
| XDD1_1259 | Methylated-DNA--protein-cysteine methyltransferase | WP_038238724.1 (89%) | WP_ 010848735.1 (93%) |
| XDD1_1272 | Ribosomal RNA large subunit methyltransferase H, RlmH | WP_038238766.1 (97%) | WP_ 010848746.1 (97%) |
| XDD1_1293 | (Dimethylallyl)adenosine tRNA methylthiotransferase MiaB | WP_038238801.1 (97%) | WP_ 010848760.1 (97%) |
| XDD1_1316 (MT3=XrdM) | Class I SAM-dependent Methyltransferase | WP_038240607.1 (67%) | WP_ 010845997.1 (55%) |
| XDD1_1319 (MT15) | Hydroxyneurosporene-O-methyltransferase | WP_038241970.1 (26%) | no |
| XDD1_1320 (MT4) | Similar to the TRP-1 protein | WP_038240039.1 (55%) | WP_ 010848326.1 (60%) |
| XDD1_1473 | Biotin synthesis protein BioC O-methyltransferase | WP_038238143.1 (83%) | WP_ 010848266.1 (86%) |
| XDD1_1508 (MT14) | Class I SAM-dependent Methyltransferase | WP_038240607.1 (83%) | WP_ 010848326.1 (52%) |
| XDD1_1539 | 23S rRNA (uracil-5-)-methyltransferase rumB | WP_038238023.1 (86%) | WP_ 010846882.1 (88%) |
| XDD1_1582 | tRNA uridine 5-oxyacetic acid(34) methyltransferase CmoM | WP_038237220.1 (91%) | WP_ 010846921.1 (93%) |
| XDD1_1599 | ribosomal RNA large subunit methyltransferase K/L | WP_038237256.1 (92%) | WP_ 013183938.1 (91%) |
| XDD1_1726 | 23S rRNA (guanine(745)-N(1))-methyltransferase | WP_038234018.1 (86%) | WP_ 010847383.1 (87%) |
| XDD1_2005 | Protein methyltransferase HemK | WP_038234838.1 (82%) | WP_ 010845680.1 (87%) |
| XDD1_2010 (MT5) | bifunctional, ab-hydrolase (2-205), and class I SAM-dependent methyltransferase | WP_038234880.1 (85%) | WP_ 010845286.1 (24%) |
| XDD1_2136 (MT6) | Bifunctional tRNA large subunit methyltransferase RlmN | WP_038234531.1 (31%) | WP_ 010847179.1 (31%) |
| XDD1_2137 (MT7) | Methyltransferase | WP_038237856.1 (28%) | WP_010848266.1 (28%) |
| XDD1_2141 (MT8) | putative methyltransferase | WP_038233467.1 (71%) | WP_ 010848326.1 (28%) |
| XDD1_2370 | Chemotaxis protein methyltransferase, CheR | WP_038241409.1 (90%) | AYA40528.1 (92%) |
| XDD1_2558 | 3-demethylubiquinone-9 3-methyltransferase | WP_038235733.1 (93%) | CCV29406.1 (93%) |
| XDD1_2631 | 50S ribosomal protein L3 N(5)-glutamine methyltransferase YfcB | WP_038236979.1 (87%) | WP_ 010846999.1 (87%) |
| XDD1_2639 | tRNA 5-methylaminomethyl-2-thiouridine biosynthesis bifunctional protein MnmC | WP_038236963.1 (80%) | WP_ 010846991.1 (82%) |
| XDD1_2773 | Ribosomal RNA large subunit methyltransferase N | WP_038234531.1 (96%) | WP_ 010847179.1 (97%) |
| XDD1_2789 | tRNA (cytidine/uridine-2'-O-)-methyltransferase TrmJ | WP_038234569.1 (90%) | WP_ 013084954.1 (91%) |
| XDD1_2817 | tRNA (adenine-N(6)-)-methyltransferase | WP_038234634.1 (78%) | WP_ 010847130.1 (81%) |
| XDD1_2936 (MT9) | class I SAM-dependent methyltransferase | WP_038237392.1 (30%) | WP_ 161783469.1 (24%) |
| XDD1_2970 (MT10) | class I SAM-dependent methyltransferase | WP_038237392.1 (28%) | WP_ 010845997.1 (27%) |
| XDD1_3026 | Ribosomal RNA small subunit methyltransferase H, RsmH | WP_038241916.1 (93%) | WP_ 010845567.1 (94%) |
| XDD1_3125 | 23S rRNA (uracil-5-)-methyltransferase RumA | WP_038242077.1 (87%) | WP_ 010845131.1 (86%) |
| XDD1_3160 | Protein-L-isoaspartate O-methyltransferase | WP_038236222.1 (90%) | WP_ 010845607.1 (91%) |
| XDD1_3199 | Ubiquinone/menaquinone biosynthesis methyltransferase ubiE | WP_038235523.1 (96%) | WP_ 010848662.1 (95%) |
| XDD1_3214 | 5-methyltetrahydropteroyltriglutamate-homocysteine S-methyltransferase | WP_038235647.1 (92%) | WP_ 010848669.1 (93%) |
| XDD1_3283 | tRNA (Thr-GGU) A37 N-methylase and TrmO C-terminal domain | WP_038236301.1 (89%) | WP_ 013185411.1 (91%) |
| XDD1_3333 (MT11) | N5-glutamine S-adenosyl-L-methionine-dependent methyltransferase | WP_038234838.1 (29%) | WP_ 010845680.1 (32%) |
| XDD1_3346 | Ribosomal RNA large subunit methyltransferase M, RlmM | WP_038236402.1 (93%) | WP_ 010848594.1 (94%) |
| XDD1_3364 | Prepilin peptidase-dependent protein C | WP_051462345.1 (72%) | WP_ 010848571.1 (71%) |
| XDD1_3389 | Ribosomal RNA small subunit methyltransferase A, RsmA | WP_038236482.1 (90%) | WP_ 010847809.1 (92%) |
| XDD1_3459 (MT12) | SAM-dependent O-methyltransferase | WP_038236584.1 (93%) | no |
| XDD1_3490 | Ribosomal RNA small subunit methyltransferase I | WP_038236668.1 (93%) | WP_ 010847779.1 (95%) |
| XDD1_3561 | Ribosomal protein L11 methyltransferase | WP_038240686.1 (95%) | WP_ 010847914.1 (94%) |
| XDD1_3579 | Ribosomal RNA small subunit methyltransferase D | WP_038240730.1 (87%) | WP_ 010848030.1 (90%) |
| XDD1_3681 | 23S rRNA A2030 N6-methylase RlmJ | WP_038238908.1 (89%) | WP_ 010848500.1 (89%) |

**Note**: 45 SAM-dependent MT were excluded based on their presence and high sequence identity (cutoff > 60%) in *X. nematophila* ATCC 19061 and *X. szentirmaii* DSM 16338. The resting 15 MT candidates were highlighted in blue and selected for the protein expression test.

**Table S3.** Strains used in this study.

| Strain | Genotype/Description | Reference |
|---|---|---|
| ***E. coli*** | | |
| DH10B | Cloning strain; F$^-$ *mcr*A, Δ(*mrr-hsdRMS-mcrBC*) Φ80*lacZ* Δ*M15* Δ*lac*X74 *rec*A1 *end*A1,araD139, Δ(*ara-leu*)7697 *gal*U, *gal*K, *rps*L,*nup*G, λ- | Invitrogen |
| TOP10 | Heterologous expression strain; *F*$^-$ *mcr*A Δ*(mrr-hsdRMS-mcrBC)* Φ80*lacZ* Δ*M15* Δ *lacX74 recA1 ara* Δ*139* Δ*(ara-leu)7697 gal*U *gal*K *rps*L *(Str*$^R$*) end*A1 *nup*G | Invitrogen |
| ST18 | *E. coli* S17-1 *λpir* Δ*hem*A | Ref[8] |
| ST18 pEB_de-MT3 (MT3=XrdM) | ST18 contains the gene *MT3* (*XDD1_1316*) deletion plasmid pEB_de-MT3 | This study |
| ST18 pEB_de-MT13 | ST18 contains the gene *MT13* (*XDD1_1102*) deletion plasmid pEB_de-MT13 | This study |
| ST18 pEB_de-MT14 | ST18 contains the gene *MT14* (*XDD1_1508*) deletion plasmid pEB_de-MT14 | This study |
| ST18 pEB_de-MT15 | ST18 contains the gene *MT15* (*XDD1_1319*) deletion plasmid pEB_de-MT15 | This study |
| TOP10 pCOLA_ToxEDBC | Heterologous expression of *tox* BGC (without ToxA) in *E.coli* TOP10 | This study |
| TOP10 pCOLA_ToxEDBC pToxA | Heterologous expression strain of *tox* BGC (without ToxA) with plasmid pToxA co-expressed | This study |
| TOP10 pCOLA_ToxEDBC pXrdM | Heterologous expression strain of *tox* BGC (without ToxA) with plasmid pXrdE co-expressed | This study |
| XL10gold | Cloning strain; Tet$^r$ Δ(*mcrA*)*183* Δ(*mcrCB-hsdSMR-mrr*)*173 end*A1 *sup*E44 *thi-1 rec*A1 *gyr*A96 *rel*A1 *lac* Hte [F´ *proAB lacI*$^q$*Z* Δ*M15* Tn*10* (Tet$^r$) Amy Cam$^r$] | Agilent |
| BL21-Gold (DE3) | Expression strain; *E. coli* B F$^-$ *omp*T *hsd*S ($r_B{}^-$ $m_B{}^-$) *dcm*$^+$ Tet$^r$ *gal* λ(DE3) *end*A Hte | Agilent |
| BL21-Gold (DE3) pETDuet1-SUMO-XrdM | Expression strain of XrdM | This study |
| BL21-Gold (DE3) pETDuet1-SUMO-XrdM-ΔNterm | Expression strain of mutant XrdM with the deletion of N-terminal residues 4-23 | This study |
| BL21-Gold (DE3) pETDuet1-SUMO-XrdM-ΔLoop | Expression strain of mutant XrdM with the deletion of residues 181-184 | This study |
| BL21-Gold (DE3) pETDuet1-SUMO-XrdM-Y7A | Expression strain of mutant XrdM with the Tyr7 substituted by Ala | This study |
| BL21-Gold (DE3) pETDuet1-SUMO-XrdM-Y7F | Expression strain of mutant XrdM with the Tyr7 substituted by Phe | This study |
| BL21-Gold (DE3) pETDuet1-SUMO-XrdM-Y14A | Expression strain of mutant XrdM with the Tyr14 substituted by Ala | This study |
| BL21-Gold (DE3) pETDuet1-SUMO-XrdM-Y18A | Expression strain of mutant XrdM with the Tyr18 substituted by Ala | This study |
| BL21-Gold (DE3) pETDuet1-SUMO-XrdM-Q23A | Expression strain of mutant XrdM with the Gln23 substituted by Ala | This study |
| BL21-Gold (DE3) pETDuet1-SUMO-XrdM-F112A | Expression strain of mutant XrdM with the Phe112 substituted by Ala | This study |
| BL21-Gold (DE3) pETDuet1-SUMO-XrdM-F112L | Expression strain of mutant XrdM with the Phe112 substituted by Leu | This study |
| BL21-Gold (DE3) pETDuet1-SUMO-XrdM-H115A | Expression strain of mutant XrdM with the His115 substituted by Ala | This study |
| BL21-Gold (DE3) pETDuet1-SUMO-XrdM-Y116F | Expression strain of mutant XrdM with the Tyr116 substituted by Phe | This study |
| BL21-Gold (DE3) pETDuet1-SUMO-XrdM-H238A | Expression strain of mutant XrdM with the His238 substituted by Ala | This study |
| BL21-Gold (DE3) pETDuet1-SUMO-DtpM | Expression strain of DtpM | This study |
| BL21-Gold (DE3) pETDuet1-SUMO-DtpM-Y138F | Expression strain of mutant DtpM with the Tyr138 substituted by Phe | This study |
| BL21-Gold (DE3) pETDuet1-SUMO-DtpM-E158A | Expression strain of mutant DtpM with the Glu158 substituted by Ala | This study |
| BL21-Gold (DE3) pETDuet1-SUMO-DtpM-H251A | Expression strain of mutant DtpM with the His251 substituted by Ala | This study |
| BL21-Gold (DE3) pETDuet1-SUMO-DtpM-N252A | Expression strain of mutant DtpM with the Asn252 substituted by Ala | This study |
| BL21-Gold (DE3) pETDuet1-SUMO-DtpM-Q301A | Expression strain of mutant DtpM with the Gln301 substituted by Ala | This study |
| BL21-Gold (DE3) pETDuet1-SUMO-DtpM-E310Q | Expression strain of mutant DtpM with the Glu310 substituted by Gln | This study |

| | | |
|---|---|---|
| ***Xenorhabdus*** | | |
| *X. nematophila* ATCC 19061 | *X. nematophila* ATCC 19061 wild type | Ref[9] |
| *X. szentirmaii* DSM 16338 | *X. szentirmaii* DSM 16338 wild type | Ref[9] |
| ***Xenorhabdus doucetiae* FRM16** | | |
| WT | *X. doucetiae* FRM16 wild type | Ref[10] |
| Δ*hfq* $P_{BAD}$_*xrdA* | *X. doucetiae* FRM16 with the deletion of *hfq* (*XDD1_0420*) and $P_{BAD}$ promoter exchanged in front of gene *xrdA* | Ref[2] |
| Δ*xrdA-H* | *X. doucetiae* FRM16 with the deletion of XRD BGC | This study |
| Δ*MT3* | *X. doucetiae* FRM16 with the deletion of *MT3* (*XDD1_1316)* | This study |
| Δ*MT13* | *X. doucetiae ΔMT13* with the deletion of *MT13* (*XDD1_1102)* | This study |
| Δ*MT14* | *X. doucetiae* with the deletion of *MT14* (*XDD1_1508)* | This study |
| Δ*MT15* | *X. doucetiae* with the deletion of *MT15* (*XDD1_1319)* | This study |
| Δ*hfq* $P_{BAD}$_*xrdA* pMT1 | *Δhfq* $P_{BAD}$_*xrdA* with the protein expression plasmid pMT1 | This study |
| Δ*hfq* $P_{BAD}$_*xrdA* pMT2 | *Δhfq* $P_{BAD}$_*xrdA* with the protein expression plasmid pMT2 | This study |
| Δ*hfq* $P_{BAD}$_*xrdA* pMT3 (MT3=XrdM) | *Δhfq* $P_{BAD}$_*xrdA* with the protein expression plasmid pMT3 | This study |
| Δ*hfq* $P_{BAD}$_*xrdA* pMT4 | *Δhfq* $P_{BAD}$_*xrdA* with the protein expression plasmid pMT4 | This study |
| Δ*hfq* $P_{BAD}$_*xrdA* pMT5 | *Δhfq* $P_{BAD}$_*xrdA* with the protein expression plasmid pMT5 | This study |
| Δ*hfq* $P_{BAD}$_*xrdA* pMT6 | *Δhfq* $P_{BAD}$_*xrdA* with the protein expression plasmid pMT6 | This study |
| Δ*hfq* $P_{BAD}$_*xrdA* pMT7 | *Δhfq* $P_{BAD}$_*xrdA* with the protein expression plasmid pMT7 | This study |
| Δ*hfq* $P_{BAD}$_*xrdA* pMT8 | *Δhfq* $P_{BAD}$_*xrdA* with the protein expression plasmid pMT8 | This study |
| Δ*hfq* $P_{BAD}$_*xrdA* pMT9 | *Δhfq* $P_{BAD}$_*xrdA* with the protein expression plasmid pMT9 | This study |
| Δ*hfq* $P_{BAD}$_*xrdA* pMT10 | *Δhfq* $P_{BAD}$_*xrdA* with the protein expression plasmid pMT10 | This study |
| Δ*hfq* $P_{BAD}$_*xrdA* pMT11 | *Δhfq* $P_{BAD}$_*xrdA* with the protein expression plasmid pMT11 | This study |
| Δ*hfq* $P_{BAD}$_*xrdA* pMT12 | *Δhfq* $P_{BAD}$_*xrdA* with the protein expression plasmid pMT12 | This study |
| Δ*hfq* $P_{BAD}$_*xrdA* pMT13 | *Δhfq* $P_{BAD}$_*xrdA* with the protein expression plasmid pMT13 | This study |
| Δ*hfq* $P_{BAD}$_*xrdA* pMT14 | *Δhfq* $P_{BAD}$_*xrdA* with the protein expression plasmid pMT14 | This study |
| Δ*hfq* $P_{BAD}$_*xrdA* pMT15 | *Δhfq* $P_{BAD}$_*xrdA* with the protein expression plasmid pMT15 | This study |
| Δ*hfq* $P_{BAD}$_*xrdA* pCTG1_5739 | *Δhfq* $P_{BAD}$_*xrdA* with the protein expression plasmid pCTG1_5739 (QTR03164.1) | This study |
| Δ*hfq* $P_{BAD}$_*xrdA* pACYC | *Δhfq* $P_{BAD}$_*xrdA* with the empty plasmid pACYC | This study |
| Δ*hfq* Δ*xrdE::hlmA* $P_{BAD}$_*xrdA* | *Δhfq* with the gene *xrdE* (XRD acetyltransferase, *XDD1_0766*) replaced by *hlmA* (HlmA, the holothin acetyltransferase WP_003955866.1), and the $P_{BAD}$ promoter exchanged in front of gene *xrdA* | This study |
| Δ*hfq* Δ*xrdE::hlmA* $P_{BAD}$_*xrdA* pXrdM | Δ*hfq* Δ*xrdE::hlmA* $P_{BAD}$_*xrdA* with the protein expression plasmid pMT3 (MT3=XrdM) | This study |
| Δ*hfq* Δ*xrdE::hlmA* $P_{BAD}$_*xrdA* pCTG1_5739 | Δ*hfq* Δ*xrdE::hlmA* $P_{BAD}$_*xrdA* with the protein expression plasmid pCTG1_5739 | This study |
| Δ*hfq* Δ*xrdE::hlmA* $P_{BAD}$_*xrdA* pDtpM | Δ*hfq* Δ*xrdE::hlmA* $P_{BAD}$_*xrdA* with the protein expression plasmid pDtpM | This study |
| Δ*hfq* Δ*xrdE::actA* $P_{BAD}$_*xrdA* | *Δhfq* with the gene *xrdE* replaced by *actA* (ActA, the putative thiolutin acetyltransferase KF719091.1), and the $P_{BAD}$ promoter exchanged in front of gene *xrdA* | This study |
| Δ*hfq* Δ*DC* $P_{BAD}$_*xrdA* | *X. doucetiea* with the deletion of *hfq* (*XDD1_0420*) and decarboxylase (DC, *XDD1_2132*) and with the $P_{BAD}$ promoter exchange in front of gene *xrdA* | This study |
| Δ*hfq* Δ*DC* $P_{BAD}$_*xrdA* pACYC | Δ*hfq* Δ*DC* $P_{BAD}$_*xrdA* with the empty plasmid pACYC | This study |
| Δ*hfq* Δ*DC* $P_{BAD}$_*xrdA* pXrdM | Δ*hfq* Δ*DC* $P_{BAD}$_*xrdA* with the protein expression plasmid pXrdM | This study |
| Δ*hfq* Δ*DC* $P_{BAD}$_*xrdA* pToxA | Δ*hfq* Δ*DC* $P_{BAD}$_*xrdA* with the protein expression plasmid pToxA | This study |
| Δ*hfq* Δ*DC* $P_{BAD}$_*xrdA* pDtpM | Δ*hfq* Δ*DC* $P_{BAD}$_*xrdA* with the protein expression plasmid pDtpM | This study |
| Δ*hfq* Δ*DC* $P_{BAD}$_*xrdA* pMT13-S18Y | Δ*hfq* Δ*DC* $P_{BAD}$_*xrdA* with the plasmid mutant pMT13-S18Y | This study |
| Δ*hfq* Δ*DC* $P_{BAD}$_*xrdA* pMT14-S18Y | Δ*hfq* Δ*DC* $P_{BAD}$_*xrdA* with the plasmid mutant pMT14-S18Y | This study |
| Δ*hfq* Δ*DC* $P_{BAD}$_*xrdA* pMT4-S18Y-C115H-H116Y-D238H | Δ*hfq* Δ*DC* $P_{BAD}$_*xrdA* with the plasmid mutant pMT4-S18Y-D238H-C115H-H116Y | This study |

**Table S4.** Plasmids used in this study.

| Plasmid | Genotype/Description | Reference |
|---|---|---|
| pACYC | p15A ori, *araC*-$P_{BAD}$, *tacI*, $Cm^R$ | Ref[11] |
| pCOLA | CollA ori, *araC*-$P_{BAD}$, *tacI*, $Kana^R$ | Ref[11] |
| pEB17 | pDS132 based, R6K ori, oriT, *cipB*, *sacB,* $kan^R$ | Ref[2] |
| pMT1 | The pACYC backbone (PCR amplified using the primer CK1007+CK1010) and gene *MT1* (PCR amplified using LS81+LS82) were ligated by Gibson assembly | This study |
| pMT2 | The pACYC backbone (PCR amplified using the primer CK1007+CK1010) and gene *MT2* (PCR amplified using LS83+LS84) were ligated by Gibson assembly | This study |
| pMT3 (=pXrdM) | The pACYC backbone (PCR amplified using the primer CK1007+CK1010) and gene *MT3* (PCR amplified using LS85+LS86) were ligated by Gibson assembly | This study |
| pMT4 | The pACYC backbone (PCR amplified using the primer CK1007+CK1010) and gene *MT4* (PCR amplified using LS87+LS88) were ligated by Gibson assembly | This study |
| pMT5 | The pACYC backbone (PCR amplified using the primer CK1007+CK1010) and gene *MT5* (PCR amplified using LS89+LS90) were ligated by Gibson assembly | This study |
| pMT6 | The pACYC backbone (PCR amplified using the primer CK1007+CK1010) and gene *MT6* (PCR amplified using LS91+LS92) were ligated by Gibson assembly | This study |
| pMT7 | The pACYC backbone (PCR amplified using the primer CK1007+CK1010) and gene *MT7* (PCR amplified using LS93+LS94) were ligated by Gibson assembly | This study |
| pMT8 | The pACYC backbone (PCR amplified using the primer CK1007+CK1010) and gene *MT8* (PCR amplified using LS95+LS96) were ligated by Gibson assembly | This study |
| pMT9 | The pACYC backbone (PCR amplified using the primer CK1007+CK1010) and gene *MT9* (PCR amplified using LS97+LS98) were ligated by Gibson assembly | This study |
| pMT10 | The pACYC backbone (PCR amplified using the primer CK1007+CK1010) and gene *MT10* (PCR amplified using LS99+LS100) were ligated by Gibson assembly | This study |
| pMT11 | The pACYC backbone (PCR amplified using the primer CK1007+CK1010) and gene *MT11* (PCR amplified using LS101+LS102) were ligated by Gibson assembly | This study |
| pMT12 | The pACYC backbone (PCR amplified using the primer CK1007+CK1010) and gene *MT12* (PCR amplified using LS103+LS104) were ligated by Gibson assembly | This study |
| pMT13 | The pACYC backbone (PCR amplified using the primer CK1007+CK1010) and gene *MT13* (PCR amplified using LS126+LS127) were ligated by Gibson assembly | This study |
| pMT14 | The pACYC backbone (PCR amplified using the primer CK1007+CK1010) and gene *MT14* (PCR amplified using LS128+LS129) were ligated by Gibson assembly | This study |
| pMT15 | The pACYC backbone (PCR amplified using the primer CK1007+CK1010) and gene *MT1*5 (PCR amplified using LS105+LS106) were ligated by Gibson assembly | This study |
| pCTG1_5739 | The pACYC backbone (PCR amplified using the primer CK1007+CK1010) and gene *ctg1_5739* (PCR amplified using LS147+LS148) were ligated by Gibson assembly | This study |
| pDtpM | The pACYC backbone (PCR amplified using the primer CK1007+CK1010) and gene *dtpM* (PCR amplified using LS222+LS223) were ligated by Gibson assembly | This study |
| pToxA | The pACYC backbone (PCR amplified using the primer CK1007+CK1010) and gene *toxA* (PCR amplified using LS226+LS227) were ligated by Gibson assembly | This study |
| pCOLA_ToxEDBC | The pCOLA backbone (PCR amplified using the primer CK1007+CK1010) and gene *toxE* (PCR amplified using LS228+LS229) and genes *toxDBC* (PCR amplified using LS230+LS231) were ligated by Gibson assembly | This study |
| pCEP_$P_{BAD}$_*xrdA* | pCEP_kan vector with the first 620 bp of xrdA was inserted through two restriction enzyme sites *Pst*I and *Xba*I | Ref[1] |
| pEB_de-MT3 (XrdM) | The pEB17 backbone (PCR amplified using primer LS142+LS143) and two homologous arms (HAL and HAR, PCR amplified using primers LS130+LS131 and LS132+LS133, respectively) were ligated by Gibson assembly | This study |
| pEB_de-MT13 | The pEB17 backbone (obtained by digestion with restriction enzyme *Bgl*II and *Pst*I) and two homologous arms (HAL and HAR, PCR amplified using primers MW636+MW637 and MW638+MW639, respectively) were ligated by Gibson assembly | This study |
| pEB_de-MT14 | The pEB17 backbone (obtained by digestion with restriction enzyme *Bgl*II and *Pst*I) and two homologous arms (HAL and HAR, PCR amplified using primers MW632+MW633 and MW634+MW635, respectively) were ligated by Gibson assembly | This study |

| | | |
|---|---|---|
| pEB_de-MT15 | The pEB17 backbone (obtained by digestion with restriction enzyme *Bgl*II and *Pst*I) and two homologous arms (HAL and HAR, PCR amplified using primers MW528+MW529 and MW530+MW531, respectively) were ligated by Gibson assembly | This study |
| pEB01 (pEB_de-DC) | The plasmid construct for deletion of decarboxylase (DC) | Ref[10] |
| pEB_de *xrdA-H* | The pEB17 backbone (obtained by digestion with restriction enzyme *Bgl*II and *Pst*I) and two homologous arms (HAL and HAR, PCR amplified using primers PEB195+PEB196 and PEB197+PEB198, respectively) were ligated by Gibson assembly | This study |
| pEB_*hlmA swap* | The pEB17 backbone (PCR amplified using primer LS142+LS143), HAL (LS171+LS172), gene *hlmA* (LS173+LS174), and HAR (LS175+LS176) were ligated by Gibson assembly | This study |
| pEB_*actA swap* | The pEB17 backbone (PCR amplified using primer LS142+LS143), HAL (LS165+LS166), gene *actA* (LS167+LS168), and HAR (LS169+LS170) were ligated by Gibson assembly | This study |
| pETDuet1-SUMO-XrdM | The pETDuet-1 backbone with a His6-SUMO-Tag inserted into the MCS1 and the gene for XrdM in frame behind. | This study |
| pETDuet1-SUMO-XrdM-ΔLoop | Construction as above for Mutant XrdM with the deletion of residues 181-184 | This study |
| pETDuet1-SUMO-XrdM-ΔNterm | Construction as above for Mutant XrdM with the deletion of residues 4-23 | This study |
| pETDuet1-SUMO-XrdM Y7A | Construction as above for Mutant XrdM with the Tyr7 substituted by Ala | This study |
| pETDuet1-SUMO-XrdM Y7F | Construction as above for Mutant XrdM with the Tyr7 substituted by Ala | This study |
| pETDuet1-SUMO-XrdM Y14A | Construction as above for Mutant XrdM with the Tyr7 substituted by Ala | This study |
| pETDuet1-SUMO-XrdM Y18A | Construction as above for Mutant XrdM with the Tyr7 substituted by Ala | This study |
| pETDuet1-SUMO-XrdM Q23A | Construction as above for Mutant XrdM with the Tyr7 substituted by Ala | This study |
| pETDuet1-SUMO-XrdM F112A | Construction as above for Mutant XrdM with the Tyr7 substituted by Ala | This study |
| pETDuet1-SUMO-XrdM F112L | Construction as above for Mutant XrdM with the Tyr7 substituted by Ala | This study |
| pETDuet1-SUMO-XrdM H115A | Construction as above for Mutant XrdM with the Tyr7 substituted by Ala | This study |
| pETDuet1-SUMO-XrdM Y116F | Construction as above for Mutant XrdM with the Tyr7 substituted by Ala | This study |
| pETDuet1-SUMO-XrdM H238A | Construction as above for Mutant XrdM with the Tyr7 substituted by Ala | This study |
| pETDuet1-SUMO-DtpM | The pETDuet-1 backbone with a His6-SUMO-Tag inserted into the MCS1 and the gene for DtpM in frame behind. | This study |
| pETDuet1-SUMO-DtpM Y138F | Construction as above for Mutant DtpM with the Tyr138 substituted by Phe | This study |
| pETDuet1-SUMO-DtpM E158A | Construction as above for Mutant DtpM with the Glu158 substituted by Ala | This study |
| pETDuet1-SUMO-DtpM H251A | Construction as above for Mutant DtpM with the His251 substituted by Ala | This study |
| pETDuet1-SUMO-DtpM N252A | Construction as above for Mutant DtpM with the Asn252 substituted by Ala | This study |
| pETDuet1-SUMO-DtpM Q301A | Construction as above for Mutant DtpM with the Gln301 substituted by Ala | This study |
| pETDuet1-SUMO-DtpM E310Q | Construction as above for Mutant DtpM with the Glu310 substituted by Gln | This study |
| pMT13-S18Y | Mutant of pMT13 with the Ser18 substituted by Tyr | This study |
| pMT14-S18Y | Mutant of pMT14 with the Ser18 substituted by Tyr | This study |
| pMT4-S18Y-C115H-H116Y-D238H | Mutant of pMT4 with the Ser18, Asp238, Cys115, and His116 substituted by Tyr and His, His, and Tyr, respectively | This study |

**Table S5.** Primers used in this study.

| Primer | Sequence | usage |
|---|---|---|
| CK1007 | CATGGAATTCCTCCTGTTAGCCCAAAAAAAC | pACYC(and pCOLA) backbone amplification |
| CK1010 | TTAATTAACCTAGGCTGCTGCCACCG | |
| LS142 | AGATCTGAGCTCTCCCGGGAATTCCAC | pEB17 backbone amplification |
| LS143 | CTGCAGGTCGACTCTAGAGGATCGATCC | |
| LS79 | CACACTTTGCTATGCCATAG | Verification of all plasmid constructs based on pACYC and pCOLA |
| LS80 | CTACCTTAGGACCGTTATAG | |
| LS081 | GTTTTTTTGGGCTAACAGGAGGAATTCCATGAACACTTCCAACATCAAAAAATATGC | Amplification of MT1 for expression |
| LS082 | CGGTGGCAGCAGCCTAGGTTAATTAACTATTCCTCCTGAGCCTTATTGCCCG | |
| LS083 | GTTTTTTTGGGCTAACAGGAGGAATTCCATGAAAGCGATAGATTTGTTTGCAGGCGC | Amplification of MT2 for expression |
| LS084 | CGGTGGCAGCAGCCTAGGTTAATTAATTATTTGTCGGTGTTTTTTCTTCCG | |
| LS085 | GTTTTTTTGGGCTAACAGGAGGAATTCCATGCAAGAAGAGATACATTATGATTCTATTG | Amplification of MT3 for expression |
| LS086 | CGGTGGCAGCAGCCTAGGTTAATTAACTATGGCATCCAACAGGTCAGTC | |
| LS087 | GTTTTTTTGGGCTAACAGGAGGAATTCCATGAAAGATAACATCTTATATAATCCTATTG | Amplification of MT4 for expression |
| LS088 | CGGTGGCAGCAGCCTAGGTTAATTAATTAGCGTTGACAAACAAGAGCGGCATC | |
| LS089 | GTTTTTTTGGGCTAACAGGAGGAATTCCATGGAAAATATCGTGGTCATAGGTCAGAG | Amplification of MT5 for expression |
| LS090 | CGGTGGCAGCAGCCTAGGTTAATTAATCATTCTCGGCGACGAATGCGTTTGG | |
| LS091 | GTTTTTTTGGGCTAACAGGAGGAATTCCATGAAAATCAGCGAAATCATGGAGTCTG | Amplification of MT6 for expression |
| LS092 | CGGTGGCAGCAGCCTAGGTTAATTAATTACATAATTCGATATTTACTGCGAA | |
| LS093 | GTTTTTTTGGGCTAACAGGAGGAATTCCATGCCAATCAACTTTAATGCCCGTGAAAATAG | Amplification of MT7 for expression |
| LS094 | CGGTGGCAGCAGCCTAGGTTAATTAATTATTTTATGCCTACCCATACCGTCC | |
| LS095 | GTTTTTTTGGGCTAACAGGAGGAATTCCATGGCAATTGACCGACTTTATCAAGACAG | Amplification of MT8 for expression |
| LS096 | CGGTGGCAGCAGCCTAGGTTAATTAACTATTTATTCTTCCTGACGATAAAAATC | |
| LS097 | GTTTTTTTGGGCTAACAGGAGGAATTCCATGAAACAATTTAATAAGAATGCTCAGGC | Amplification of MT9 for expression |
| LS098 | CGGTGGCAGCAGCCTAGGTTAATTAATTACCTAAGTTTGACCGCAGTAAACG | |
| LS099 | GTTTTTTTGGGCTAACAGGAGGAATTCCATGACAGCTATCGATTATTCTTCTCTATC | Amplification of MT10 for expression |
| LS100 | CGGTGGCAGCAGCCTAGGTTAATTAATTAATACTTTTGAGCATGTAGTAAAAAC | |
| LS101 | GTTTTTTTGGGCTAACAGGAGGAATTCCATGAATTTCTCTCATGAATATCAAATTTT | Amplification of MT11 for expression |
| LS102 | CGGTGGCAGCAGCCTAGGTTAATTAATTAAAAGTTATTTATACCAATAGTAATAAC | |
| LS103 | GTTTTTTTGGGCTAACAGGAGGAATTCCATGACCTCACCCTATAATTATGACAAAGATG | Amplification of MT12 for expression |
| LS104 | CGGTGGCAGCAGCCTAGGTTAATTAATTACTGCTTCTTGCAGATCAGTAATG | |
| LS126 | GTTTTTTTGGGCTAACAGGAGGAATTCCATGGACGAGCAAGTAAGCTATG | Amplification of MT13 for expression |
| LS127 | CGGTGGCAGCAGCCTAGGTTAATTAATTAAAACTGACAAACAAGTCCGGTGTG | |
| LS128 | GTTTTTTTGGGCTAACAGGAGGAATTCCATGGACGAGCAAGCAAGCTATG | Amplification of MT14 for expression |
| LS129 | CGGTGGCAGCAGCCTAGGTTAATTAATTAAAACTGACAAACAAGCCCGGTGTG | |
| LS105 | GTTTTTTTGGGCTAACAGGAGGAATTCCATGGGCATTGATATGAACACACTCAATTATAAC | Amplification of MT15 for expression |
| LS106 | CGGTGGCAGCAGCCTAGGTTAATTAATTATTCGGCAATGGCTTCGACAATC | |
| LS147 | GTTTTTTTGGGCTAACAGGAGGAATTCCATGGTAAGTACATCAGCTGAAGAGATAGCGG | Amplification of CTG1_5739 for expression |
| LS148 | CGGTGGCAGCAGCCTAGGTTAATTAATTATTTAACGCCGCTCGTGACCAGGAAC | |
| LS130 | GATCGATCCTCTAGAGTCGACCTGCAGTTTCTACCCAATCAGTTGAGCTG | Amplification of HAL and HAR for deletion of *MT3* |
| LS131 | CCGGGAGAGCTCAGATCTCCAAATATAACTTTTTCCCGACTG | |
| LS132 | CTAAAATATATCCCCCAAATTATTGGGGGCCATAATCATGGTTCCCTTATTAAG | |
| LS133 | CTTAATAAGGGAACCATGATTATGGCCCCCAATAATTTGGGGGATATATTTTAG | |
| LS134 | CTATATGCCTGGCTCAATTATTATTG | Verification of the gene deletion of *MT3* |
| LS135 | GATCCATGTTAAACAGAGGGATATG | |
| LS165 | CGATCCTCTAGAGTCGACCTGCAGAACTCGTCAGAGCTAATGACCCC | Amplification of HAL, gene *actA*, and HAR for replacement of gene *xrdE* |
| LS166 | CAATTGCCAGCTTGGAGCTCGGTTAATCATCATTAACAAAGATATTCCTATTGAG | |
| LS167 | GATTAACCGAGCTCCAAGCTGGCAATTGCATATACATTACG | |
| LS168 | GATATAAGGGTTATGACTACGACAGATGTTAATCAACATCACG | |
| LS169 | CGTGATGTTGATTAACATCTGTCGTAGTCATAACCCTTATATCCTCTCCAACAACGATC | |
| LS170 | CCCGGGAGAGCTCAGATCTGATCGTTGCCCATAATGATGTGGTG | |
| LS171 | GATCGATCCTCTAGAGTCGACCTGCAGAACTCGTCAGAGCTAATGACCCCAC | Amplification of HAL, gene *hlmA*, and HAR for replacement of gene *xrdE* |
| LS172 | GCCATTACCACCCTGGAACTGGGCTAATCATCATTAACAAAGATATTC | |
| LS173 | TTAGCCCAGTTCCAGGGTGGTAATGGCAAAG | |
| LS174 | ATGAATAGTCACGATGCTTTTACAGTATCAACGGCGACC | |
| LS175 | GTTGATACTGTAAAAGCATCGTGACTATTCATAACCCTTATATCCTCTCCAACAAC | |

| | | |
|---|---|---|
| LS176 | GGAATTCCCGGGAGAGCTCAGATCTGATCGTTGCCCATAATGATGTGGTG | |
| LS155 | CACCAATCGACAATATTTCGGTGGGG | Verification of the gene replacement of *xrdE* |
| LS156 | CCGTGATCCTGACCGAAGAGCTGATC | |
| LS222 | GTTTTTTTGGGCTAACAGGAGGAATTCCATGGCGGAAACCGATGATGAACGTGCTGGCG | Amplification of DtpM for expression |
| LS223 | CGGTGGCAGCAGCCTAGGTTAATTAATCACGGGGTCGGGCCGGCCGGTGCAGGTAC | |
| LS226 | GTTTTTTTGGGCTAACAGGAGGAATTCCATGAAAAATGAAGAGTCCTATGATTCTATTG | Amplification of ToxA for expression |
| LS227 | CGGTGGCAGCAGCCTAGGTTAATTAACTAAAATTGGCAGATAAAACCAGTTTG | |
| LS228 | GTTTTTTTGGGCTAACAGGAGGAATTCCATGACACTCGATGAGATATATATGTCTCGTG | Amplification of ToxE |
| LS229 | CTAAACGGGACGCAAACGAAGACGGACATCAGGCCCAATTTGTTTTAC | |
| LS230 | CCGTCTTCGTTTGCGTCCCGTTTAGATTTGACATCTGTTATTTATATTGACCGAATC | Amplification of ToxDBC |
| LS231 | CGGTGGCAGCAGCCTAGGTTAATTAATTATGTTCTAACTTCAATTGGATATAG | |
| EV1 | CGTAAAGTTGAAGTTCGTAC | Mutagenesis of XrdM; Deletion of residues 4-23 |
| EV2 | GGATCCACCGATCTGTTC | |
| EV3 | ATCCTGCTGACCTTCTAC | Mutagenesis of XrdM; Deletion of residues 181-184 |
| EV4 | GATGAATTCAACCTGGTG | |
| EV5 | CGAAATCCACGCGGACTCTATCGGTGAAGG | Mutagenesis of XrdM: Y7A |
| EV6 | GATCCACCGATCTGTTCAC | Mutagenesis of XrdM: Y7A |
| EV7 | CGAAATCCACTTTGACTCTATCGGTG | Mutagenesis of XrdM: Y7F |
| EV8 | GATCCACCGATCTGTTCAC | Mutagenesis of XrdM: Y7F |
| EV9 | CGGTGAAGGTGCGGAACACTTCTACAAC | Mutagenesis of XrdM: Y14A |
| EV10 | ATAGAGTCGTAGTGGATTTC | Mutagenesis of XrdM: Y14A |
| EV11 | CGAACACTTCGCGAACTCTGTTCCGCAG | Mutagenesis of XrdM: Y18A |
| EV12 | TAACCTTCACCGATAGAG | Mutagenesis of XrdM: Y18A |
| EV13 | CTCTGTTCCGGCGCGTAAAGTTG | Mutagenesis of XrdM: Q23A |
| EV14 | TTGTAGAAGTGTTCGTAAC | Mutagenesis of XrdM: Q23A |
| EV15 | CACCGCTACCGCGCTGTTCCACTAC | Mutagenesis of XrdM: F112A |
| EV16 | ATGATGTCGAATTTTTCGTTC | Mutagenesis of XrdM: F112A |
| EV17 | CACCGCTACCCTGCTGTTCCACT | Mutagenesis of XrdM: F112L |
| EV18 | ATGATGTCGAATTTTTCGTTCAGC | Mutagenesis of XrdM: F112L |
| EV19 | CTTCCTGTTCGCGTACGCTAAATCTATCGTTGAACTG | Mutagenesis of XrdM: H115A |
| EV20 | GTAGCGGTGATGATGTCG | Mutagenesis of XrdM: H115A |
| EV21 | CCTGTTCCACTTTGCTAAATCTATCG | Mutagenesis of XrdM: Y116F |
| EV22 | AAGGTAGCGGTGATGATG | Mutagenesis of XrdM: Y116F |
| EV23 | GAACTGCATGGCGACCGGTCTGACCTGC | Mutagenesis of XrdM: H238A |
| EV24 | TGCTGGTAGGTGTCCCAG | Mutagenesis of XrdM: H238A |
| EV25 | TTCTCCGCTGTTTGAATACATGGC | Mutagenesis of DtpM: Y138F |
| EV26 | CCGAAGTGTTCGTCGAAG | Mutagenesis of DtpM: Y138F |
| EV27 | TATGCGTGGTGCGTCTCTGGCTACCGC | Mutagenesis of DtpM: E158A |
| EV28 | GCAGCAGCGAAACGAGCA | Mutagenesis of DtpM: E158A |
| EV29 | ATCTACCCTGGCGAACTGGGACGACG | Mutagenesis of DtpM: H251A |
| EV30 | TTAACCAGGTACAGGTCAG | Mutagenesis of DtpM: H251A |
| EV31 | TACCCTGCACGCGTGGGACGACG | Mutagenesis of DtpM: N252A |
| EV32 | GATTTAACCAGGTACAGG | Mutagenesis of DtpM: N252A |
| EV33 | TAAAGACCTGGCGATGCTGGTTCTGC | Mutagenesis of DtpM: Q301A |
| EV34 | ACGTACGGGTTCAGTTCA | Mutagenesis of DtpM: Q301A |
| EV35 | GGGTGGTCGTCAGCGTACCCGTG | Mutagenesis of DtpM: E310Q |
| EV35 | AGCAGAACCAGCATTTGCAG | Mutagenesis of DtpM: E310Q |
| PEB195 | CCTCTAGAGTCGACCTGCAGCACAACGGCAGCCTATATGG | Amplification of HAL and HAR for deletion of genes *xrdA-H* |
| PEB196 | CTCTTCCACGAGGCGATTAATCTCGCTCCTACATTTGACCGAGG | |
| PEB197 | AATCCTCGGTCAAATGTAGGAGCGAGATTAATCGCCTCGTGGA | |
| PEB198 | TCCCGGGAGAGCTCAGATCTGAGAGGTAATCAGGGTGGCG | |
| PEB199 | CAGCAGGAACAGATACCTGC | Verification of the genes deletion of *xrdA-H* |
| PEB200 | GTTCATTGTCGGGATAGGTAG | |
| PEB201 | CGGATAACTAACCAAACCG | |
| PEB202 | TATGTCAGCAATAGGGTGAC | |
| MW636 | CCTCTAGAGTCGACCTGCAGCGATTTGGGCTGTATTGATGG | Amplification of HAL and HAR for deletion of gene *MT13* |
| MW637 | GAGAGCATGAGCGGTGAGCGTCCATTATTATAACTCCTGATAAAG | |
| MW638 | GTCCATTATTATAACTCCTGATAAAGGCTCACCGCTCATGCTC | |

| | | |
|---|---|---|
| MW639 | TCCCGGGAGAGCTCAGATCTGCACATAACATTGCCGCG | |
| MW632 | CCTCTAGAGTCGACCTGCAGGATTTGGGCGGTGTTGATG | Amplification of HAL and HAR for deletion of gene *MT14* |
| MW633 | GAGAGCATGAGCAGTGAGCGTCCATTATTATAACTCCTGATAAAG | |
| MW634 | CTTTATCAGGAGTTATAATAATGGACGCTCACTGCTCATGCTCTC | |
| MW635 | TCCCGGGAGAGCTCAGATCTGCACATAACATCGCCGCG | |
| MW528 | CCTCTAGAGTCGACCTGCAGGGTGCTGCAAAAGTGATTGG | Amplification of HAL and HAR for deletion of gene *MT15* |
| MW529 | CTATCTGCCTTAATGTAGAAGGAATTGTTTGATGAAGTAGGTGTAAAGC | |
| MW530 | GCTTTACACCTACTTCATCAAACAATTCCTTCTACATTAAGGCAGATAG | |
| MW531 | TCCCGGGAGAGCTCAGATCTCTCTGCAATCGATATTTCTTCAC | |
| LS241 | GTATTATGAAAAATTTTACAATACGGTCGCTCAACG | Site mutagenesis of MT13 and MT14 (S18Y) |
| LS242 | CGTTGAGCGACCGTATTGTAAAATTTTTCATAATAC | |
| LS243 | GAGAGTTTTTACGACGCCGGCG | Site mutagenesis of MT4 (S18Y-C115H-H116Y-D2438H) |
| LS244 | CGCCGGCGTCGTAAAAACTCTC | |
| LS245 | CCACTATGCGGAATCACTTGAACAACTTG | |
| LS246 | CATAGTGGAATAACCAGGCTGCAACTATC | |
| LS247 | GAAAAACTGTCTTCATGCCGCTCTTG | |
| LS248 | CAAGAGCGGCATGAAGACAGTTTTTC | |
| LS249 | CCACCATGCGGAATCACTTGAACAACTTG | |
| LS250 | CATGGTGGAATAACCAGGCTGCAACTATC | |

**Table S6.** Synthesized genes.

| Gene | Sequence (codon-optimized) |
|---|---|
| *hlmA* | ATGAATAGTCACGATGCTTTTACAGTATCAACGGCGACCTTGGAGGACTGGTATCAGGTTGCGGAGTGGGCAGACGGCG<br>AAGGTTGGAACGTGGGCGACGGCGATGTTGCCTGCTTCCACCCGACTGACCCGGCGGGTTTCTTTATCGGCCGCCTGGG<br>TGCTCGCCCTGTGGCCGCTGTGAGCATTGTTAATTACGATGATCGTTATGCAGTGCTGGGTCATTATCTGACGGATCCG<br>GAGTTTCGTGGTCGTGGCTACGGCTTGGCGACCTGGAAAGCAGCATTTCCGCATAGCGGTAACCGTACCGTAGGCCTTG<br>ACGCGATGCCGGCGCAACGTGCGAACTACGAAACCCATGGTTTCAAGGCAGCGCATGATACCGTTCATTTCGCGGGCTC<br>GCCGGCGCGCCCAACGGGCCCGGTCCAGGGCGTTGCTCCGGTTACCCCGGAGCACGCGGAAGCACTCGCGGCGTACGAC<br>CGTGGTTGTTTCCCGGCGGACCGTTCTGGTTTCGTCGGCCGCTGGCTGACCGCTCCGGGTCGCACCGCTCGCGTGCGTC<br>TGCGCGATGGCGCGGTGGCTGGTTACGGCGTGATCCGTCCGGCTGGCAGAGGCCACCGTATTGGTCCGCTGTTCGCGGA<br>CACCCCGGAGGACGCCGCCGCGCTGTTCGATGGTTTGGTTGGTCACTTTGGTCCGGGTGAAGAAGTTTCCCTGGATATC<br>CCGGGTACTCACGCAGCGAGCGCAGACTTGCTGACGAGCCGTGGTTTAGCTGCCCAATTTCACACCGTGCGTATGTATA<br>CCGGTCCAGTGCCGGAGACTGCCGAAGAGCGTGTCTTTGCCATTACCACCCTGGAACTGGGCTAA |
| *actA* | GTAACTACGACAGATGTTAATCAACATCACGACCTGGAAGATTTCACCGTGGACACGGCTACCGTTGACGATTGGCGTCA<br>GATTACCGCTTGGGGCAACCAGGAGGGCTGGAACATTGGTTTCCACGATGCAGAGTGCTTTTTCGCTGCAGACCCGGGCG<br>GTTTCTTCATCGGCCGTGTCGGTGGCCGCCCAGTGTCGGCGGTGTCTATGGTGAACTATAGCGATGAATTTAGCGCGTGG<br>GGTCATTATCTGGTGGACCCGCGTTTACGCGGTCGTGGCTACGGTCAGGGTGTGTGGGAAGTCGCGGTTCGTCATGCAGG<br>ATCTCGTGCGGCGAGCGGTGATGCCATGCCGGGGGTCGTGGGCTTCTACCGCCGTGAAGGTATGGTTCCGGTTCATCATA<br>CCGTTCATTGGGCAGGCCGGCTGGCACGTGCGGGCCGCGAGGTGGACGGCGTTGAACCGGTTAGACCGGAGCAACTGGGT<br>GCGGTTGCTGACTATGATGGTGAATGTTTCCCGGCGCACCGTCCGGGCTTCCTGGGCCGCTGGCTGTTTGCACCGGGCCA<br>CGTGGCCAGAGCTCGCTGGGCGGGTGGTCGTGTTACGGGTTACGGCGTGCTACGCCCAGCACCGGCTGGCTACCGCATCG<br>GTCCGTTGGTTGCGGACACTCCGCGTATCGCCGCGGAGGTGTTTGATGCATTGACCGCGCACCTGCGTCCGGGCGCCGAG<br>GTCAGCGCGTTTTCCCCGGAAGTGCAAGAGGCGGCGGCTCCGCTGCTGTCCGCGGGTGGTCTGGCGGAACGTTTTCGTTT<br>GGTTCGTGTGCACCGTGGTGCGCCTCCGGCTCACCGCGCTCGTAATGTATATGCAATTGCCAGCTTGGAGCTCGGTTAA |
| *ctg1_5739* | GTAAGTACATCAGCTGAAGAGATAGCGGCAGGCTACGAAGAATATGATGATCTGCCGGAACGTGTGTTGGGCTACCCGAC<br>CGCATTTTCCGCGTTGCGCCTGGGTGCTCCGGACGTTCGCACCGTTTTGGATTACGGTTGTGGTCCGGGGAAGGTGGCCC<br>TGCGCGTGGCGGAGGGTTATGGCGCCAGAGTGCACGCGGTCGACATCAGCGCTCGCATGCTGGCAATTGCGCGCGCTCGC<br>CGTGCGCATCCGCTGGTGGCGTACCATCTGGTGGAAGGCGCCTCCCTGCGTTTCCTGCCGGACGACAGCGTTGATGCGGC<br>ATTGTGCTGCTTTGTTTTCGTGACGGTTGCGGAGGCAGACCGTCTCCGCGAAGTTTGTGCAGAGGTTCACCGTGTTTTAC<br>GTCCGGGCGGTCGTTTCGCCGTGCTTGACGTGAACCCGGATGCTACCGGTATTCGTTTCTCGACCTTTCGTACCGGCGAG<br>CCGGGCGCACGTTATCGTCCGGGTCAGCAACGTCCGACTTGGCTGCACTTGCCGGATGGTGGCGTGCTGGACTTGGCTGA<br>TCATCACTGGCCGCGTTCTGCGTACCGTGCGGCTCTGGCTGGCGCGGGTTTTGGTGATATCGCCTGCGCGGCGCCATTAC<br>TGGGTGGTGCGGACGCGCCTGCGGAAGCGCCACCGGGTAGCCTGTTGGACGTCGAGCGCCCGGCGGAGGCACTGCACCCG<br>CCGTTCCTGGTCACGAGCGGCGTTAAATAA |
| *dtpM* | ATGGCGGAAACCGATGATGAACGTGCTGGCGCTCGCTCCCTGCTGATGCAACGCCTGTTCGGCTCCCGTGTTACCGAAGT<br>GCTGGCGGCGATGGCGCGTCTGGATCTGGCAGATGCCATCGGCGATGGTATTGCTGACGTACATGACCTGGCGCGTTCTT<br>GTGACCTGCCGGCTGACGGCCTGCACCGTCTGCTGCGCGCTCTGGCTGGTCTGGGTATGTGCGAAGAATCCGAGCCGGGT<br>AAATTCGCGCTGACCGCAAGCGGTGCCCTGCTGCGTAAAGACCATCCGGAGTCCGTTTATGATTTCGCTCGCTTCCATAC<br>TGCCCCGGAGACCACTCGTCCGTGGACGAATCTGGAACAGGCGCTGCGTACCGGCCGTCCTACCTTCGACGAGCATTTCG<br>GTTCTCCGCTGTACGAATACATGGCTGGCCACCCGGAGCTGAGCGCACGTTTCGCTGCTGCGATGCGCGGTGAATCTCTG<br>GCGACCGCTGACACCATTGCTGAGCACTACGACTTCTCTCCATACCGTACTGTGACCGACGTCGGTGGCGGCGACGGCAC<br>CCTGATCACCGCAATTCTGCGCCGTCACCCGGACCTGCGTGGCACCATCTTCGAAACCCCGGAAATTGCTGAACGTGCAG<br>CTGAGCGTGTACGTGCAGCTGGCCTGCATGACCGCTGCGCCGTTGTAAGCGGCGACTTTTTCGACCTGGTACCGGGCGGC<br>GCTGACCTGTACCTGGTCAAATCCACCCTGCACAACTGGGACGACGAACACGTAGTGCGCATCCTGAGCAGCTGCCGCAC<br>CGCGCTGGCAGATCGTGGTCGTCTGCTGGTTATCGACGTGGTTCTGCCAGACCGTGCCGAACCGGATCCGGCGGAGCTGA<br>ACCCGTACGTTAAGGACCTGCAGATGCTGGTTCTGCTGGGTGGCCGTGAACGTACCCGTGCACATCTGGATCGCCTGTGT<br>GCACGTGCTGGTCTGGTTATCGACCGTGTGTCCTGCCGCTGCCTCCGCATGTTGGTCTGAGCCTGACCGAAGTTGTACCTGC<br>ACCGGCCGGCCCGACCCCGTGA |
| *xrdM* | GAAATCCACTACGACTCTATCGGTGAAGGTTACGAACACTTCTACAACTCTGTTCCGCAGCGTAAAGTTGAAGTTCGTAC<br>CCTGTTCGACATGGTTGGTGACGTTCAGGGTAAATCTGTTCTGGACCTGGCTTGCGGTTACGGTTACTTCGGTCGTGAAC<br>TGTACCACCGTGGTGCTTCTAAAGTTGTTGGTGTTGACATCTCTGAAAAAATGATCGCTCTGGCTAAAAAGAAATCTACC<br>GAATACGGTGACAACATCGAATTCCACGTAGCGAACGTAAGCGACATGCAGCTGAACGAAAAAATTCGACATCATCACCGC<br>TACCTTCCTGTTCCACTACGCTAAATCTATCGTTGAACTGGAATCTATGTTCCGTTCTGTTGCTAACCACCTGAAACCGT<br>CTGGTAAACTGGTTGCTTACATGGCTGCTCCGGACTACCAGCTGGAAAAAGGTAACTGCCACAACTACGGTCTGAACATC<br>CTGTCTGAAGAACCGCTGCAAGGTGGTTTCATCCACCAGGTTGAATTCATCACCACCCCGCCGATCCTGCTGACCTTCTA<br>CCGTTGGGACCGTGAAACCTACAAAAACGCTATCCACAAAGCTGGTTTCGGTCACTTCGAATGGCGTAAACCGATGGTTC<br>TGGAATCTGACATCGAACGTTACCCGGCTGGTTTCTGGGACACCTACCAGCAGAACTGCATGCACACCGGTCTGACCTGC<br>TGGATGCCGTAA |

**Table S7.** NCBI accession number of proteins in this study.

| Protein | NCBI accession number | Locus tag |
|---|---|---|
| MT1 | WP_001095581.1 | XDD1_0024 |
| MT2 | WP_000643690.1 | XDD1_0699 |
| MT3 (XrdM) | WP_010845997.1 | XDD1_1316 |
| MT4 | WP_010845997.1 | XDD1_1320 |
| MT5 | WP_000431866.1 | XDD1_2010 |
| MT6 | WP_045970778.1 | XDD1_2136 |
| MT7 | WP_045970780.1 | XDD1_2137 |
| MT8 | WP_045970788.1 | XDD1_2141 |
| MT9 | WP_045971961.1 | XDD1_2836 |
| MT10 | WP_231854520.1 | XDD1_2970 |
| MT11 | WP_045972546.1 | XDD1_3333 |
| MT12 | WP_045972703.1 | XDD1_3459 |
| MT13 | WP_045969284.1 | XDD1_1102 |
| MT14 | WP_045969875.1 | XDD1_1508 |
| MT15 | WP_084721119.1 | XDD1_1319 |
| CTG1_5739 | QTR03164.1 | |
| HlmA | WP_003955866.1 | |
| ActA | KF719091.1 | |
| DtpM | QTR03060 | |
| ToxA | PHM34299.1 | Xsze_00722 |
| ToxB | PHM34300.1 | Xsze_00723 |
| ToxC | PHM34301.1 | Xsze_00724 |
| ToxD | PHM34302.1 | Xsze_00725 |
| ToxE | PHM33766.1 | Xsze_00151 |

**Table S8.** Isolation conditions for the purification of XRD-271 and XRD-271Me.

| **Isolation of XRD-271** | |
|---|---|
| *1. Prep* | |
| Solvents | ACN/$H_2O$ +0.1%FA |
| 0 min | 15% |
| 2 min | 15% |
| 28 min | 60% |
| Retention time of XRD-271 | 19.1 min |
| Column | waters XBridge BEH C18 19x250mm |
| Detection UV | 50 mAU |
| *2. Semi-prep* | |
| Solvents | ACN/$H_2O$ +0.1%FA |
| 0 min | 38% |
| 2 min | 38% |
| 28 min | 38% |
| Retention time of XRD-271 | 24.8 min |
| Column | Phenomenex Phenyl-Hexyl 10x250mm |
| Detection UV | 20 mAU |
| **Isolation of XRD-271Me** | |
| *1. Prep* | |
| Solvents | ACN/$H_2O$ +0.1%FA |
| 0 min | 15% |
| 2 min | 15% |
| 33 min | 40% |
| Retention time of XRD-271Me | 19.5 min |
| Column | waters XBridge BEH C18 19x250mm |
| Detection UV | 60 mAU |
| *2. Semi-prep* | |
| Solvents | ACN/$H_2O$ +0.1%FA |
| 0 min | 38% |
| 2 min | 38% |
| 28 min | 40% |
| Retention time of XRD-271Me | 24.3 min |
| Column | Phenomenex Phenyl-Hexyl 10x250mm |
| Detection UV | 35 mAU |
| **Amount of pure compounds** | |
| XRD-271 | 64.4 mg |
| XRD-271Me | 84.2 mg |

**Table S9.** X-ray data collection and refinement statistics for XrdM and DtpM structures.

| | **XrdM:SAH** | **DtpM:SAH** | **DtpM:SAH :XRD-271** | **DtpM:SAH :XRD-271Me** |
|---|---|---|---|---|
| ***Crystal parameters*** | | | | |
| Space group | $P4_32_12$ | $P4_122$ | $P2_12_12$ | $P4_122$ |
| Cell constants | a= b= 72.5 Å<br>c= 105.0 Å | a= 53.2 Å<br>b= 53.2 Å<br>c= 207.9 Å | a= 112.4 Å<br>b= 218.8 Å<br>c= 55.3 Å | a= 53.4 Å<br>b= 53.4 Å<br>c= 209.3 Å |
| Subunits / AU[a] | 1 | 1 | 4 | 1 |
| ***Data collection*** | | | | |
| Beam line | X06SA, SLS | X06SA, SLS | X06SA, SLS | X06SA, SLS |
| Wavelength (Å) | 1.0 | 1.0 | 1.0 | 1.0 |
| Resolution range (Å)[b] | 46-2.1<br>(2.2-2.1) | 47-1.75<br>(1.85-1.75) | 49-2.2<br>(2.3-2.2) | 47-1.75<br>(1.85-1.75) |
| No. observations | 92135 | 162253 | 303383 | 141583 |
| No. unique reflections[c] | 16946[#] | 30822[#] | 64433[#] | 31122[#] |
| Completeness (%)[b] | 99.6 (99.8) | 98.4 (98.5) | 91.5 (94.4) | 97.6 (97.2) |
| $R_{merge}$ (%)[b, d] | 4.1 (69.7) | 7.0 (67.5) | 9.2 (67.6) | 4.9 (66.9) |
| $I/\sigma$ (I)[b] | 18.6 (2.3) | 11.5 (1.9) | 10.7 (2.1) | 16.7 (2.2) |
| ***Refinement (REFMAC5)*** | | | | |
| Resolution range (Å) | 30-2.1 | 30-1.75 | 30-2.2 | 30-1.75 |
| No. refl. working set | 16059 | 29272 | 61187 | 29553 |
| No. refl. test set | 845 | 1540 | 3220 | 1555 |
| No. non hydrogen | 1667 | 2797 | 10902 | 2782 |
| Solvent ($H_2O$) | 22 | 93 | 228 | 72 |
| $R_{work}/R_{free}$ (%)[e] | 20.0/22.9 | 17.6/21.9 | 19.3/22.2 | 20.3/22.0 |
| r.m.s.d. bond (Å) / angle (°)[f] | 0.003/1.168 | 0.002/1.206 | 0.005/1.382 | 0.004/1.313 |
| Protein | 66.8 | 34.3 | 43.6 | 31.4 |
| SAH | 53.1 | 28.2 | 33.0 | 25.0 |
| Ligand | - | - | 42.1 | 29.0 |
| Ramachandran Plot (%)[g] | 97.5/2.5/0.0 | 98.8/1.2/0.0 | 99.2/0.8/0.0 | 98.8/1.2/0.0 |
| PDB accession code | 8RDL | 8RDM | 8RDN | 8RDO |

[a] Asymmetric unit

[b] The values in parentheses for resolution range, completeness, $R_{merge}$ and $I/\sigma$ (I) correspond to the highest resolution shell

[c] Data reduction was carried out with XDS and from a single crystal.

[#] Friedel pairs were treated as identical reflections

[d] $R_{merge}(I) = \Sigma_{hkl}\Sigma_j \mid I(hkl)_j - \langle I(hkl)\rangle \mid / \Sigma_{hkl}\, \Sigma_j\, I(hkl)_j$, where $I(hkl)_j$ is the $j^{th}$ measurement of the intensity of reflection hkl and $\langle I(hkl)\rangle$ is the average intensity

[e] $R = \Sigma_{hkl} \mid |F_{obs}| - |F_{calc}| \mid / \Sigma_{hkl} |F_{obs}|$, where $R_{free}$ is calculated without a sigma cut off for a randomly chosen 5% of reflections, which were not used for structure refinement, and $R_{work}$ is calculated for the remaining reflections

[f] Deviations from ideal bond lengths/angles

[g] Percentage of residues in favored / allowed / outlier region

**Table S10.** Proteins structurally related to XrdM according to Dali search.

| PDB entry code | Z-Score | R.m.s.d. [Å] | Identity [%] | Protein |
|---|---|---|---|---|
| 5je4 | 30.5 | 1.5 | 55 | Methyltransferase ToxA |
| 5hik | 22.1 | 2.1 | 18 | Glycine Sarcosine *N*-methyltransferase |
| 3bkw | 21.5 | 2.8 | 24 | *S*-adenosylmethionine dependent methyltransferase (NP_104914.1) from *Mesorhizobium loti* |
| 1z3c | 21.2 | 2.4 | 22 | *Encephalitozooan cuniculi* mRNA Cap (Guanine-N7) methyltransferase |
| 1ve3 | 20.7 | 2.8 | 25 | PH0226 protein from *Pyrococcus horikoshii* OT3 |
| 3d2l | 20.2 | 2.9 | 22 | SAM-dependent methyltransferase (ZP_ 00538691.1) from *Exiguobacterium sp.* 255-15 |
| 4hgz | 20.1 | 2.7 | 17 | CcbJ methyltransferase from *Streptomyces caelestis* |
| 3g5l | 20.1 | 3.3 | 23 | Putative *S*-adenosylmethionine dependent methyltransferase from *Listeria monocytogenes* |
| 1vl5 | 20.0 | 2.7 | 17 | Putative methyltransferase (BH2331) from *Bacillus halodurans* C-125 |
| 6uk5 | 20.0 | 2.2 | 19 | Structure of SAM bound CalS10, an amino pentose methyltransferase from *Micromonospora echinaspora* involved in calicheamicin biosynthesis |

The Dali server identified numerous structures that display 3D similarity to XrdM. The best 10 hits are shown and listed according to their Z-score. Redundant protein hits have been excluded from the list.

**Table S11.** Proteins structurally related to DtpM according to Dali search.

| PDB entry code | Z-Score | R.m.s.d. [Å] | Identity [%] | Protein |
|---|---|---|---|---|
| 6c5b | 41.6 | 1.9 | 40 | LaPhzM |
| 3gwz | 39.5 | 2.4 | 36 | Mitomycin 7-O-methyltransferase MmcR |
| 1tw3 | 37.0 | 2.7 | 37 | Carminomycin-4-O-methyltransferase (DnrK) |
| 3i5u | 36.6 | 3.1 | 33 | O-methyltransferase (NcsB1) from neocarzinostatin biosynthesis |
| 6clx | 36.4 | 3.2 | 34 | TnmH in complex with SAM |
| 5ice | 36.3 | 2.8 | 25 | (S)-norcoclaurine 6-O-methyltransferase |
| 4a6d | 36.1 | 2.1 | 24 | Human N-acetylserotonin methyltransferase (ASMT) in complex with SAM |
| 5cvv | 35.8 | 3.0 | 26 | coniferyl alcohol bound monolignol 4-O-methyltransferase 9 |
| 6i73 | 35.8 | 2.9 | 27 | Structure of Fragaria ananassa O-methyltransferase in complex with S-adenosylhomocysteine and protocatechuic aldehyde |
| 3p9i | 35.5 | 3.2 | 24 | Crystal structure of perennial ryegrass LpOMT1 complexed with S-adenosyl-L-homocysteine and sinapaldehyde |

The Dali server identified numerous structures that display 3D similarity to DtpM. The best 10 hits are shown and listed according to their Z-score. Redundant protein hits have been excluded from the list.

**Supplementary Reference**


1. Bode, E. et al. Simple "on-demand" production of bioactive natural products. *Chembiochem* **16**, 1115-9 (2015).
2. Bode, E. et al. Promoter Activation in Δhfq Mutants as an Efficient Tool for Specialized Metabolite Production Enabling Direct Bioactivity Testing. *Angew Chem Int Ed Engl* **58**, 18957-18963 (2019).
3. Chen, X., Johnson, R.M. & Li, B. A Permissive Amide N-Methyltransferase for Dithiolopyrrolones. *ACS Catalysis* **13**, 1899-1905 (2023).
4. Jiang, J. et al. Functional and Structural Analysis of Phenazine O-Methyltransferase LaPhzM from Lysobacter antibioticus OH13 and One-Pot Enzymatic Synthesis of the Antibiotic Myxin. *ACS Chem Biol* **13**, 1003-1012 (2018).
5. Fenwick, M.K., Philmus, B., Begley, T.P. & Ealick, S.E. Burkholderia glumae ToxA Is a Dual-Specificity Methyltransferase That Catalyzes the Last Two Steps of Toxoflavin Biosynthesis. *Biochemistry* **55**, 2748-59 (2016).
6. Ilyina, M.G., Khamitov, E.M., Ivanov, S.P., Mustafin, A.G. & Khursan, S.L. Anions of uracils: N1 or N3? That is the question. *Computational and Theoretical Chemistry* **1078**, 81-87 (2016).
7. Ganguly, S. & Kundu, K.K. Protonation/deprotonation energetics of uracil, thymine, and cytosine in water from e.m.f./spectrophotometric measurements. *Canadian Journal of Chemistry* **72**, 1120-1126 (1994).
8. Thoma, S. & Schobert, M. An improved Escherichia coli donor strain for diparental mating. *FEMS Microbiol Lett* **294**, 127-32 (2009).
9. Neubacher, N. et al. Symbiosis, virulence and natural-product biosynthesis in entomopathogenic bacteria are regulated by a small RNA. *Nat Microbiol* **5**, 1481-1489 (2020).
10. Bode, E. et al. Biosynthesis and function of simple amides in Xenorhabdus doucetiae. *Environ Microbiol* **19**, 4564-4575 (2017).
11. Lorenzen, W., Ahrendt, T., Bozhüyük, K.A. & Bode, H.B. A multifunctional enzyme is involved in bacterial ether lipid biosynthesis. *Nat Chem Biol* **10**, 425-7 (2014).
